# Quantifying single cell lipid signaling kinetics after photo-stimulation

**DOI:** 10.1101/2023.01.27.525833

**Authors:** David T. Gonzales, Milena Schuhmacher, H. Mathilda Lennartz, Juan M. Iglesias-Artola, Sascha M. Kuhn, Pavel Barahtjan, Christoph Zechner, André Nadler

**Affiliations:** Swiss Federal Institute of Technology Lausanne, 1015 Lausanne, Switzerland; Max Planck Institute of Molecular Cell Biology and Genetics, 01307 Dresden, Germany; Center for Systems Biology Dresden, 01307 Dresden, Germany

**Author notes:** These authors contributed equally (David T. Gonzales, Milena Schuhmacher).

**Keywords:** lipid signaling, cell-to-cell variability, parameter identifiability, profile likelihood

## Abstract

Studying the role of molecularly distinct lipid species in cell signaling remains challenging due to a scarcity of methods for performing quantitative lipid biochemistry in living cells. We have recently used lipid uncaging to quantify lipid-protein affinities and rates of lipid transbilayer movement and turnover in the diacylglycerol signaling pathway using population average time series data. So far, this approach does not allow to account for the cell-to-cell variability of cellular signaling responses. We here report a framework that allows to uniquely identify model parameters such diacylglycerol-protein affinities and transbilayer movement rates at the single cell level for a broad variety of structurally different diacylglycerol species. We find that lipid unsaturation degree and longer side chains generally correlate with faster lipid transbilayer movement and turnover and higher lipid-protein affinities. In summary, our work demonstrates how rate parameters and lipid-protein affinities can be quantified from single cell signaling trajectories with sufficient sensitivity to resolve the subtle kinetic differences caused by the chemical diversity of cellular signaling lipid pools.

## Introduction

Biological signaling networks process information through biochemical reaction networks. However, individual cells differ in shape, size, and molecular composition and are also subject to different sources of noise and variability [1]. An accurate description of cell signaling thus requires methodologies to quantitatively analyze the kinetics of the underlying biochemical reactions in single cells [2–4]. To investigate fast biological processes in the second to minute timescale, light-induced perturbations such as optogenetics, photo-switching, or uncaging approaches are commonly employed due to their high spatial and temporal precision [5,6]. While initially pioneered for soluble metabolites and cytoplasmic proteins, optical perturbations have been increasingly used to investigate dynamic processes in lipid and membrane biology over the last decade. Photo-switching and uncaging experiments have been instrumental for analyzing cellular lipid signaling events, *e.g*. revealing important details about the PIP2-synaptotagmin interaction during synaptic vesicle fusion [7] or the role of arachidonic acid signaling in insulin secretion [8]. The cellular functions of the second messenger diacylglycerol (DAG) represent a particularly intriguing case. Various DAG species which differ in acyl chain length, unsaturation and positioning at the glycerol backbone are generated through the action of phospholipases C and D (PLC and PLD) after receptor activation. These DAGs recruit cytosolic effector proteins such as protein kinase C (PKC) isoforms [9] or MUNC13 [10] proteins to the plasma membrane through C1-domain-DAG interactions, ultimately triggering downstream signaling responses. Early reports suggested that the structural diversity is functionally relevant for the signaling outcome [11–13]. We recently provided the mechanistic underpinning for this notion using a combination of plasma-membrane specific DAG uncaging and mathematical modelling of C1-domain recruitment dynamics to the plasma membrane [14]. In these experiments, we equipped native DAG species with a photoremovable group that blocked their biological functions and pre-localized the resulting caged DAGs to the outer leaflet of the plasma membrane. Light-induced cleavage of the photo-removable group induces a well-defined concentration increase of a specific DAG species at the plasma membrane. We demonstrated that this approach can be used to extract quantitative lipid-protein affinities and rates of lipid trans-bilayer movement across the plasma membrane from population-average time trace data. However, it does not account for cell-to-cell heterogeneity in DAG signaling events or variation of photoreaction yields among single cells, which might cause a loss of information and less accurate parameter estimates.

In the present study, we exploit the full information content of single-cell trajectories to infer lipid-protein affinities and kinetic parameters. The key challenge for quantifying photo-induced biochemical reactions in single cells is the fact that photoreaction yields are influenced by cell architecture, cell cycle stages, and other factors that contribute to cellular heterogeneity. While average photoreaction yields for lipid uncaging at the plasma membrane can be determined in principle, it is currently not possible to simultaneously measure signaling trajectories and photoreaction yield in the same cell because the utilized assays require independent experimental approaches. One way to address this problem is to include the number of photo-released molecules for each cell as an unknown model parameter and infer it together with all other parameters. How well this approach works in practice, however, depends on whether the additional degree of freedom will affect parameter identifiability. This can be systematically determined using identifiability analyses such as the profile likelihood method. Based on the profile likelihood analysis, model parameters can then be classified into three categories: (1) identifiable parameters that can be uniquely determined from a given set of measurements, (2) *practically* non-identifiable parameters due to insufficient data amount or quality, and (3) *structurally* non-identifiable parameters resulting from the mathematical structure of the underlying model [15]. Note that the last type of non-identifiability, typically resulting from an overparameterization of the model, cannot be resolved by collecting more or more accurate data.

Using simulated single cell timelapse data, we first determined the DAG signaling model structure and data characteristics that are needed to satisfy structural and practical identifiability. We found that all parameters of the models we previously proposed for lipid signaling dynamics were structurally identifiable despite treating single cell photoreaction yields as unknown parameters. However, we found that model parameters easily become practically non-identifiable with a decreasing signal-to-noise ratio. Practical identifiability can be partially recovered by including a greater number of single cell traces in the analysis, so we used this approach to determine rate parameters and photoreaction yields from experimental single-cell traces of six structurally different DAG species. Two of the considered DAG species (stearoyl-arachidonylglycerol, SAG and stearoyl-oleoylglycerol, SOG), were also featured in our previous work [14] and thus allowed a direct comparison of the respective methodologies utilized for data analysis. From our results, we found that increased acyl chain unsaturation led to faster rates of lipid trans-bilayer movement and turnover. Lipid protein affinities followed a similar trend with the notable exception of SOG, suggesting that individual DAG species interact differently with the lipid binding domain. Taken together, our study demonstrates the feasibility of inferring the photoreaction yield of uncaging experiments together with model parameters while accounting for cell-to-cell heterogeneity. Our analytical approach should be generally applicable to other optical perturbation experiments in living cells where photoreaction yields are experimentally difficult to obtain - making quantitative analysis of dynamic processes in cell biology more readily accessible.

## Results

### Structural and practical identifiability of a model of DAG-driven protein recruitment to the plasma membrane

We used simulated data and profile likelihood analysis to determine the structural and practical identifiability of the parameters of a minimal model of DAG-driven protein recruitment to the plasma membrane. DAG signaling typically involves the production of the signaling DAG species at the plasma membrane after receptor activation. Subsequently, the generated DAGs either activate integral membrane proteins (*e.g*. TRP channels) by allosteric modulation or recruit C1-domain containing cytoplasmic effector proteins (*e.g*. novel PKC isoforms). We consider the latter process here. Receptor-induced production of DAGs can be mimicked by photo-release of native DAGs at the plasma membrane, which allows to study the effects of individual, structurally unique DAG species. Experimentally, the signaling event can then be monitored in time by observing the intracellular localization of DAG reporter proteins such as the C1-EGFP-NES construct [14]. To analyze such data, we consider a minimal model of DAG signaling as shown in Fig. 1. Cells are initially loaded with a caged DAG (cgDAG) in the outer leaflet of the cell membrane. The caging group prevents the DAG to flip into the inner leaflet of the membrane. Upon photoactivation by exposure to UV light, the caging group is cleaved and the DAG is liberated. Free DAG can move between the inner and outer leaflet of the membrane. Once DAG is in the inner leaflet, it can be metabolized by the cell or recruit the C1-EGFP-NES reporter from the cytosol. This process can be mathematically described by

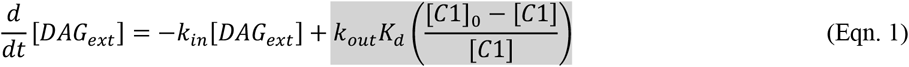

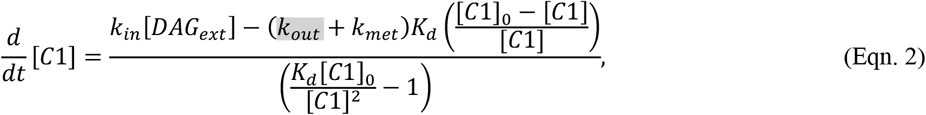

where DAGext is the uncaged DAG in the outer leaflet, C1 is the C1-EGFP-NES reporter in the cytosol, k_in_ and k_out_ are the rate constants of DAG flip-flop between the inner and outer leaflet, k_met_ is the rate constant of DAG turnover in the inner leaflet, and K_d_ is the equilibrium constant of DAG with C1 (*see Section 1 of the Supplementary Materials for model derivation*).

**Figure 1.**
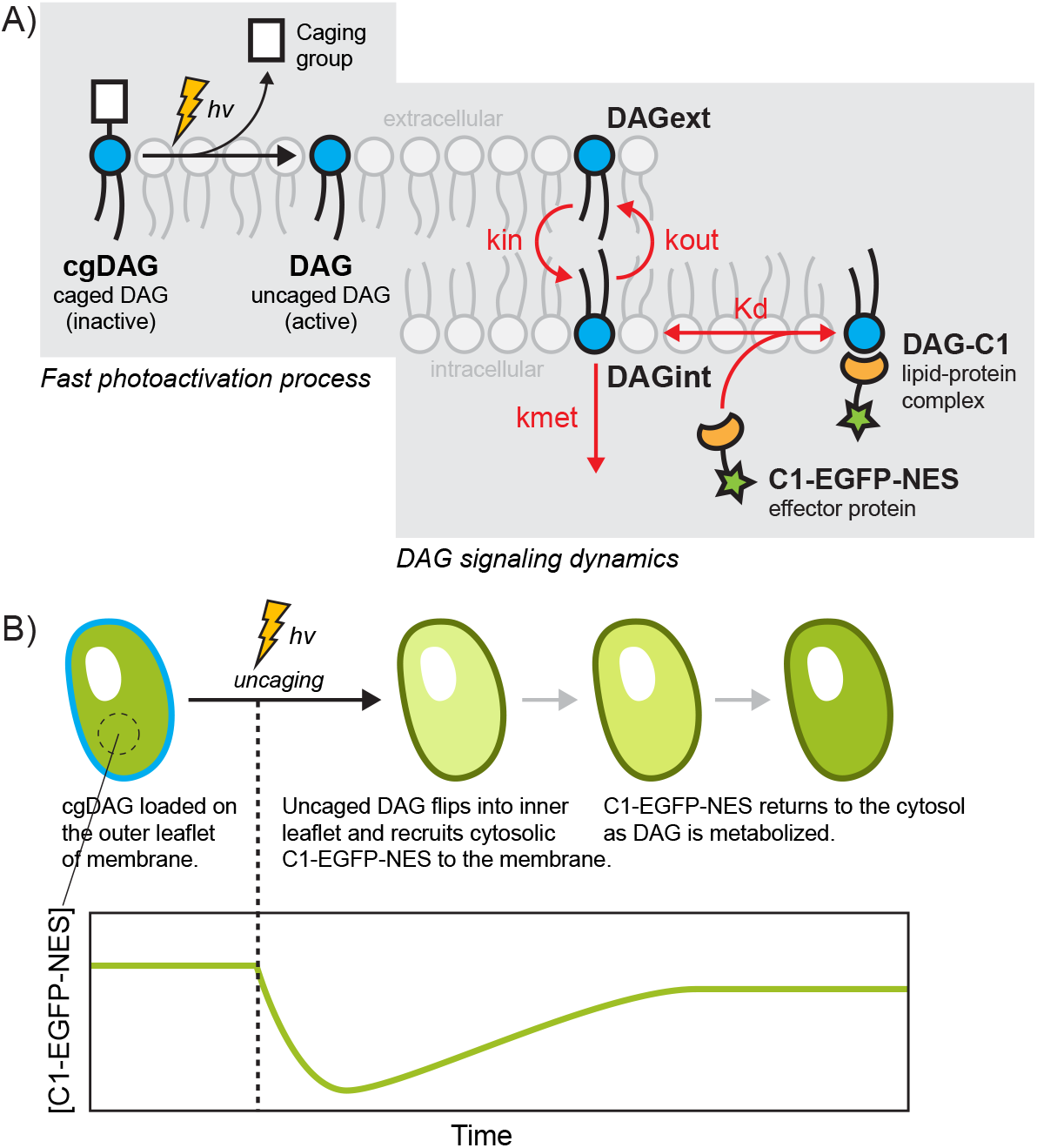
Photoactivation and DAG signaling dynamics. (A) Photoactivation or uncaging of caged diacylglycerols (cgDAG) in the outer leaflet of the cell membrane initiates the DAG signaling dynamics. Activated DAG can flip-flop between the outer and inner leaflets of the membrane. In the inner leaflet, DAG can either be metabolized by the cell or recruit the C1-EGFP-NES effector protein to form the DAG-C1 lipid-protein complex. (B) In an uncaging experiment, cells expressing C1-EGFP-NES are pre-loaded with cgDAG on the outer leaflet of the cell membrane. C1-EGFP-NES concentrations in the cytosol are monitored over time by confocal microscopy during the uncaging experiment. As cgDAG is uncaged, DAG molecules flip into the inner leaflet and recruits C1-EGFP-NES from the cytosol. This results in a drop of C1-EGFP-NES in the cytosol. As the cell metabolizes DAG molecules, C1-EGFP-NES returns into the cytosol.

In our previous work, we observed that, for some DAG species, the inside-out trans-bilayer movement (k_out_) is virtually zero (*e.g*. for SOG) and can be omitted from the model [14]. In the model version without k_out_, terms with k_out_ (in grey boxes in Eqns. 1-2) are omitted. Brackets denote the effective concentration of the species over the cell volume. Subscripts of zero indicate initial conditions at t = 0 s. After uncaging DAGs by UV exposure, cytosolic C1-EGFP-NES levels drop as DAG flips into the inner leaflet and recruits C1-EGFP-NES to the membrane. The cytosolic C1-EGFP-NES level then slowly recovers as DAGs are metabolized by the cell and C1- EGFP-NES is returned to the cytosol (Fig. 1B). Experimentally, we can easily monitor the cytosolic levels of the C1-EGFP-NES reporter in single cells over time using confocal time lapse microscopy. In contrast, DAG levels and C1-EGFP-NES densities on the inner leaflet cannot be quantified easily in a time-dependent fashion. In our previous study, identifying the model parameters k_in_, k_out_, k_met_ and K_d_ in Eqns. 1-2 [14] required measuring both initial DAG concentrations and C1-EGFP-NES levels as population-average time traces. This requires the ability to measure photoreaction yields from optical perturbations, which is currently impossible in single cells. We therefore assessed whether all model parameters can still be identified if only a single photo-stimulation magnitude is used and photoreaction yields in individual cells are not known. Specifically, we assume the model parameters k_in_, k_out_, k_met_ and K_d_ to be identical in all cells, whereas the initial DAG- and C1-EGFP-NES concentrations are allowed to vary. We hypothesized that the additional information content of single cell traces may be sufficient to carry out such an analysis. Using the DAG uncaging system enabled direct comparison of parameter estimates obtained from analyses with and without knowledge of photoreaction yields. We initially used simulated timelapse data from a single cell without measurement noise at different levels of uncaged DAG to determine the parameter identifiability based on profile likelihood analysis [15,16]. A profile likelihood shows the maximal likelihood (or minimized negative log-likelihood) of a model for a given observed dataset while sweeping one single parameter across different values. If a parameter is identifiable, its profile likelihood should exhibit an optimum in the considered parameter space. This is used to determine identifiability and likelihood-based confidence intervals of a parameter (*see Section 4 of the Supplementary Materials*). In the model versions with and without k_out_, all model parameters were found to be identifiable using a single trace without measurement noise (Fig. 2B & D, *and Fig. S1B & D in the Supplementary Materials*). Adding increasing levels of Gaussian measurement noise resulted in impaired identifiability indicated by the gradual flattening of the profile likelihoods (Fig. 2C & E, *and Fig. S1C & E in the Supplementary Materials*). The effect of increasing measurement noise was much stronger for the more complex model with inside-out trans-bilayer movement as k_out_, k_met_, and K_d_ were already weakly-identifiable with [C1] measurement noise of 0.001μM (standard deviation) which is much lower than typical experimental measurement noise levels of around 0.1μM. We then tested whether practical nonidentifiability stemming from measurement noise could be remedied by simultaneously fitting 10-200 single cell traces with 0.1μM measurement noise. When using multiple cell traces, we consider all rate parameters to be fixed over the population of cells, while each individual cell can have different initial values of [DAG] and [C1]. In the model without k_out_, all parameter non-identifiabilities were recovered at 50-200 traces (Fig. 2F, *and Figs. S2-3 and Table S1 in the Supplementary Materials*), while k_out_, k_met_, and K_d_ remained non-identifiable for the model with k_out_ with 100 traces (*Fig. S4 in the Supplementary Materials*). Overall, using simulated data, we found that our models of DAG lipid signaling dynamics are structurally identifiable, in fact, a single cell trace with no measurement noise is sufficient to identify all model parameters and the initial DAG concentration. However, measurement noise and additional model parameters (k_out_) can significantly affect practical identifiability. Analyzing multiple cell traces can help alleviate this problem to some extent, but improvements become quickly marginal as the number of cells increases.

**Figure 2.**
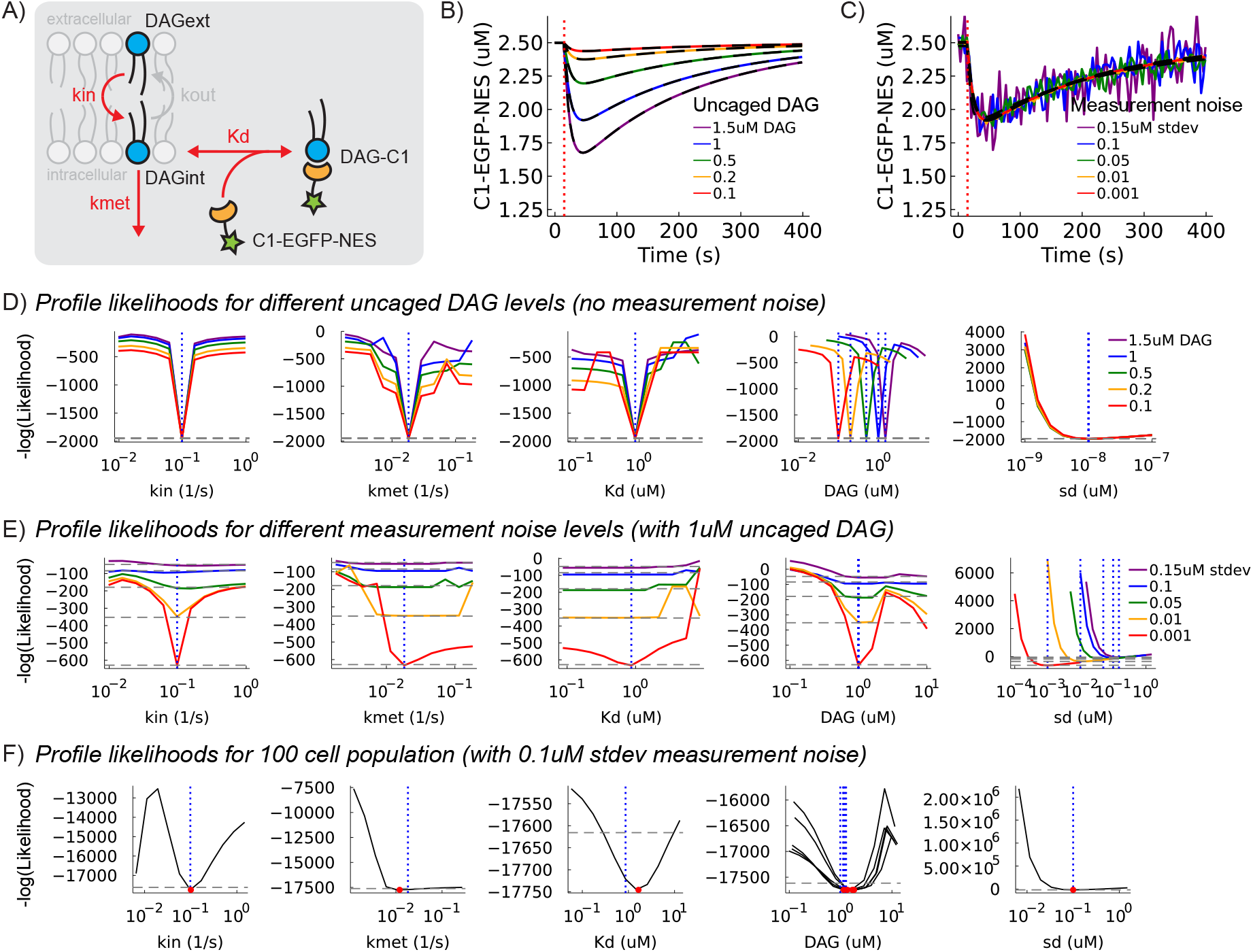
Approach and testing of parameter identifiability in a model of signaling lipid dynamics using simulated data. (A) Schematic of a model for signaling lipid dynamics. DAG at the outer leaflet of the cell membrane (DAGext) can flip into the inner leaflet (DAG_int_) where it can either be metabolized or recruit C1-EGFP-NES proteins in the cytosol to for the membrane-associated complex DAG-C1. Rate parameters (k_in_, k_met_, K_d_) of the model are shown in red. Simulated single cell traces of C1-EGFP-NES at (B) different uncaged DAG concentrations (0.1, 0.25, 0.5, 1.0, 2.0 μM) and no measurement noise and (C) different levels of measurement noise (0.001, 0.01, 0.05, 0.1, 0.15 μM) and fixed uncaged DAG at 1.5 μM. Fits of each individual cell are shown in black dashed lines. Red vertical dotted line indicates the time of UV exposure that results in DAG uncaging. Profile likelihoods of the model parameters from each individual cell at different uncaged DAG concentrations and levels of measurement noise are shown in (D) and (E), respectively. (F) Profile likelihoods of the model parameters using data from traces of 100 cells with a fixed measurement noise of 0.1 μM. Profile likelihoods of uncaged DAG are shown for five representative cells only. For all profile likelihood plots, grey horizontal dashed lines indicate the 95% likelihood-based confidence interval threshold, blue vertical dotted lines indicate the true value of the parameter, and red circles indicate the maximum likelihood estimation (MLE) of the parameter. True parameter values for the model are k_in_ = 0.098 s^-1^, k_met_ = 0.01823 s^-1^, and K_d_ = 0.866 μM. The parameter sd is an additional fitting parameter that represents the standard deviation of the measurement noise.

### Quantifying C1-EGFP-NES recruitment dynamics in single-cell DAG uncaging experiments

Having established that model parameters can be identified in principle without knowing the photoreaction yield, we next tested whether this holds true for experimental time trace data. In order to sample signaling dynamics across the chemical diversity of cellular DAG species, we expanded the repertoire of caged DAGs by synthesizing various new probes in addition to the previously reported species [14] (Fig. 3A). DAG structures were chosen to reflect the physiologically available chemical space featuring fatty acid chain lengths between 14 and 20 carbon atoms containing between 0 and 8 double bonds per DAG. Among the generated probes, dimyristoylglycerol (DMG), which contains two saturated (14:0) chains is the DAG with the shortest acyl chains commonly found in mammalian cells. Diarachidonylglycerol (DArG), contains two (20:4) chains and represents a species with long side chains and high unsaturation degree. SOG, the most common DAG in mammalian cells, bears an oleoyl (18:1) residue at the S*n*2 position and a stearoyl (18:0) residue at the S*n*1 position. Oleoyl-stearoylglycerol (OSG), featuring the same residues with switched attachment sites, was included to provide insight into the influence of fatty acid positioning. SAG, which is the major product of PLC mediated PI(4,5)P2 cleavage and widely considered to be the archetypical signaling DAG, bears an arachidonyl (20:4) residue at the S*n*2 position and a stearoyl (18:0) residue at the S*n*1 position. Finally, the biologically inactive regioisomer 1,3-dioleoylglycerol (1,3-DOG) features two oleoyl (18:1) chains at the S*n*1 and S*n*3 positions and was used as a negative control in this study. Caged DAGs were synthesized according to previously published procedures [14]. All new compounds were characterized by NMR, HRMS and coumarin-containing compounds were photochemically characterized (*see Section 9 of the Supplementary Materials for details on the synthesis of cgDAGs*). Quantification of cgDAG loading was carried out as described before [14]. Briefly, cgDAGs were loaded into the plasma membrane by exposing cells to a brief pulse (4 mins) of the respective probe in imaging buffer. The cellular localization was analyzed for all compounds by monitoring the intrinsic fluorescence of the coumarin group by confocal fluorescence microscopy. All compounds were found to localize to the plasma membrane (Fig. 3B). Loading concentrations were adjusted to achieve comparable incorporation levels for all cgDAGs (*Fig. S19 in the Supplementary Materials*).

**Figure 3.**
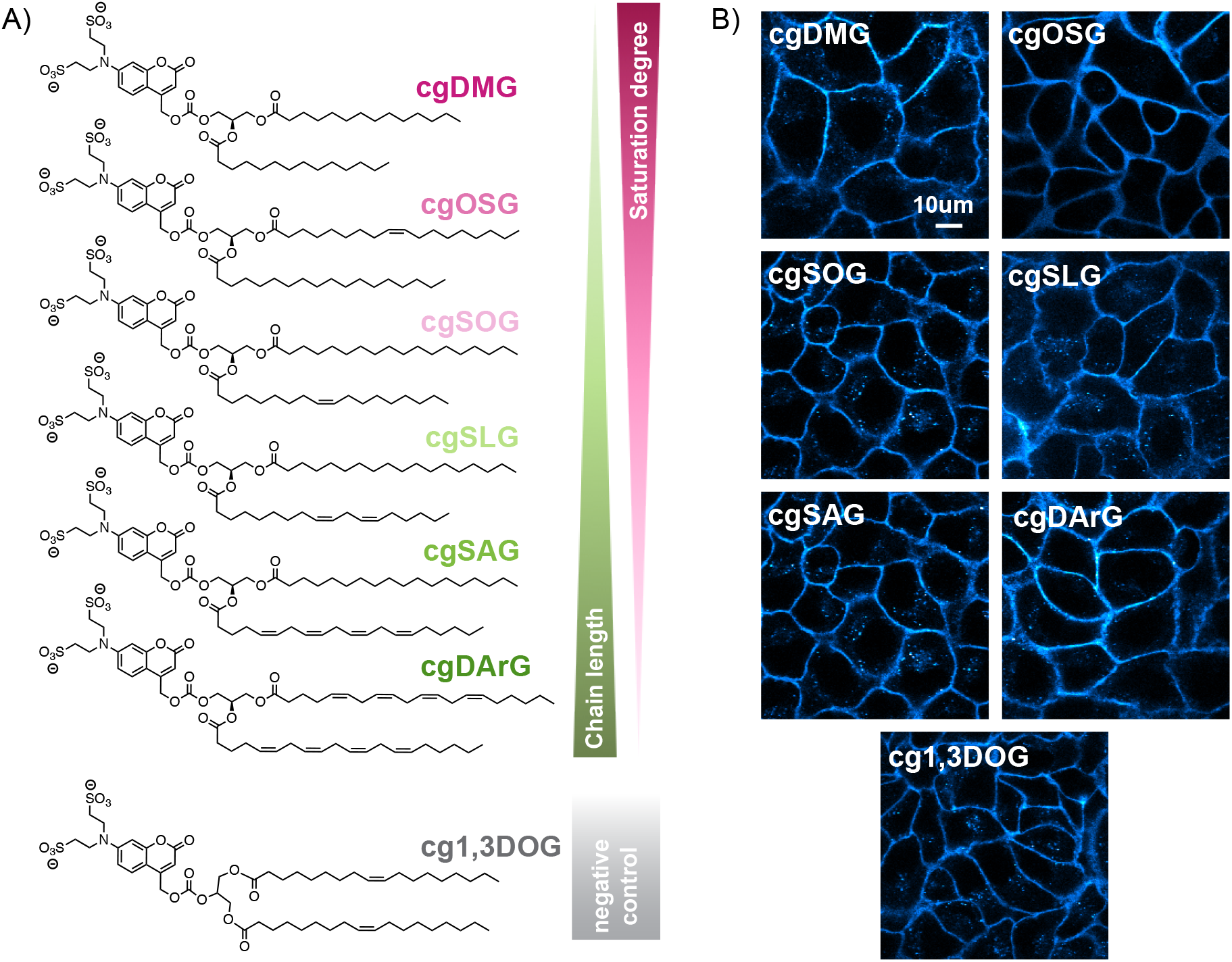
Chemistry of diacylglycerols (DAG). (A) Chemical structures of the different DAGs used in this study sorted by increasing saturation degree (top to bottom). 1,3DOG is does not recruit the C1 containing effector protein and is used as a negative control. (B) Confocal fluorescence microscopy of HeLa Kyoto cells with caged DAGs (cgDAG) localizing in the plasma membrane.

In an uncaging experiment, cells transiently expressing the C1-EGFP-NES sensor were loaded with the respective cgDAG. Uncaging was carried out by scanning the entire field of view using a 405nm laser with identical settings between experiments over a period of approximately 5s after acquiring a short baseline of five frames. We monitored the concentration of C1-EGFP-NES by comparing the time-dependent fluorescence intensity with a calibration curve obtained by measuring the fluorescence intensity of purified protein samples of defined concentration (*see Section 8 of the Supplementary Materials for fluorescence calibration details*). Single cell time traces were obtained by semi-automated image analysis (Fig. 4A, C). Briefly, cells were automatically segmented and cytosolic regions determined by eroding the detected cell shapes. The data was manually curated by removing traces from apoptotic cells and cells that moved too much during the acquisition (*see Sections 7-8 of the Supplementary Materials for details on uncaging experiments and image analysis*). Quantitative single cell time traces were obtained by converting fluorescence intensities into absolute concentrations with the calibration curve (Fig. 4B). Using this method, we obtained quantified single cell traces (80-280 cells for each DAG species) of cytosolic C1-EGFP-NES protein recruitment dynamics to the plasma membrane for the different DAG species. We also measured average photoreaction yields (*Section 7 of Supplementary Materials*), which is required for performing a population-level analysis using our previously reported methodology [14], and compare these results to our new single-cell approach (*Table S5 in the Supplementary Materials*).

**Fig.4.**
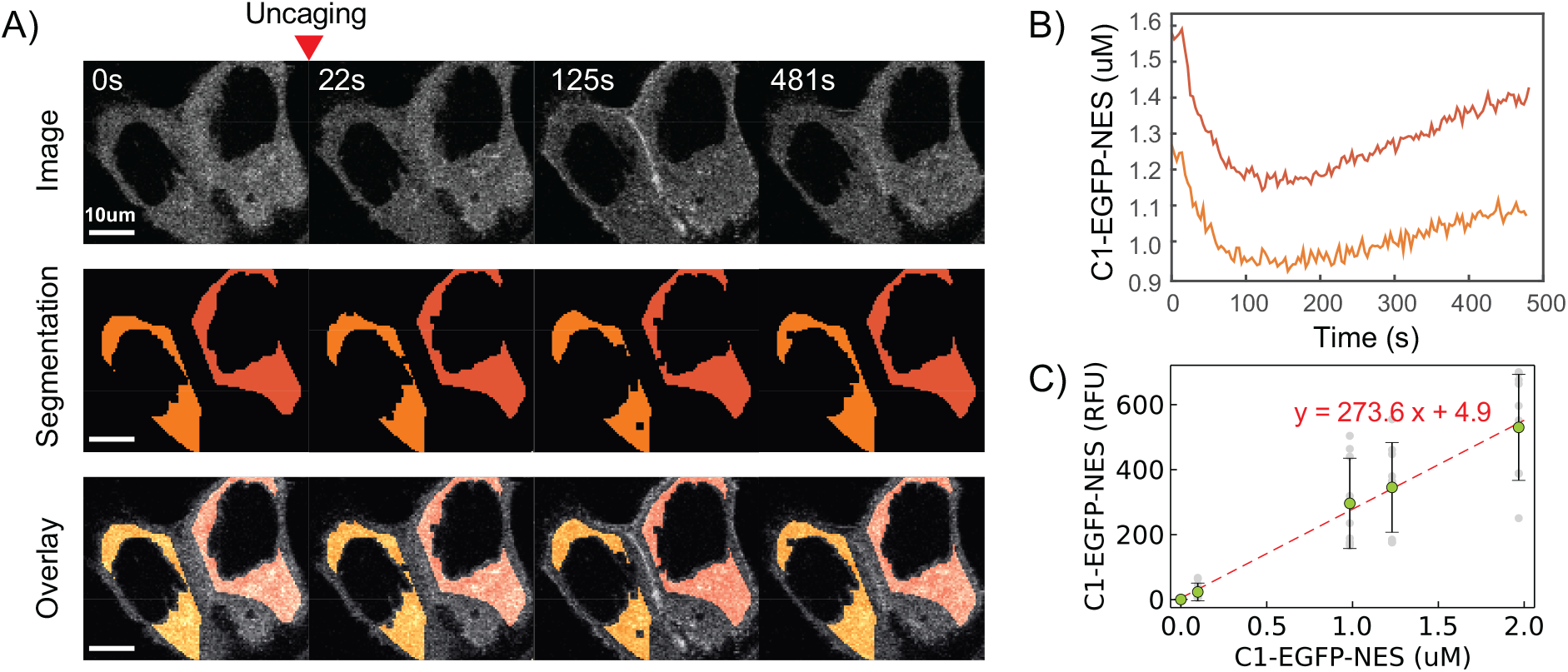
Timelapse microscopy and image analysis of DAG uncaging experiments. (A) Representative timelapse images of two cells during a DAG uncaging experiment. From the raw images, the segmented cytosolic regions are used to obtain single cell fluorescence values in the cytosol that are calibrated to protein concentration values. (B) Extracted data of C1-EGFP-NES concentration vs. time of the two representative cells in the DAG uncaging experiment. (C) Calibration curve of concentration (μM) vs. fluorescence signal (RFU) of C1-EGFP-NES in solution using the same confocal microscopy imaging settings. Grey dots are raw data points, green dots with error bars are mean and standard deviations. Red dashed line is the linear fit of the calibration curve (y = 273.6x + 4.9).

### Inference of rate parameters and photoreaction yields from single-cell uncaging experiments

We used the obtained single cell traces from uncaging experiments of each DAG species to parameterize the model of lipid signaling dynamics in Eqns. 1-2. We decided to use the simplified model featuring solely the inward trans-bilayer movement of the DAG (without k_out_) as most parameters of the model with k_out_ remained practically non-identifiable even using data from 100 cell traces in our analysis with simulated data (*Fig. S4 in the Supplementary Materials*). Utilizing the model without k_out_ is justifiable for many DAG species. For example, our previous work showed small differences in Akaike’s information criterion (AIC) and differences less than 0.5% in residual sum of squares (RSS) between both models for SAG and SOG. In addition, SOG had negligible k_out_ rates in the more complex model [14]. For the sake of simplicity, we therefore decided to use the simpler model where k_out_ is considered to be 0. We remark, however, that inside-out transbilayer movement may play a significant role for certain DAG species and/or membrane compositions in which case the more complex model would have to be considered. From results of the simulated dataset using the model without k_out_, we expected that the available number of single cell traces would be sufficient to avoid practical non-identifiability. Representative inference results for SAG are shown in Fig. 4A-C (*see Section 6 of the Supplementary Materials for the remaining DAG species*). For each DAG species, the model could fit all single cell traces from the heterogeneous population despite allowing only two parameters to vary across the population (the measured initial C1-EGFP-NES and inferred initial DAG concentrations). Initial DAG concentrations were bounded between 0-10 μM during parameter estimation. This boundary was based on a number of practical considerations: If we assume an average cell volume of 3000 μm^3^ [14], an estimated outer leaflet cell membrane surface area of 2600 μm^2^ (*see Section 8 in the Supplementary Materials*), a phospholipid area on the cell membrane of 0.65 nm^2^ [17], cgDAGs loading at 0.2% of the total lipid content of the outer leaflet cell membrane, and not more than 50% of cgDAGs uncaging with our methodology [14], this corresponds to approximately 1.77 μM of uncaged DAG per cell, which is well below the set upper bound of 10 μM. The mean values of inferred initial DAG concentrations were all within 1-4 μM and comparable to our measured average uncaged DAG concentrations (Fig. 5B, *and Fig. S12 and Table S5 in the Supplementary Materials*), which indicates that our fitting results are within reasonable and relevant ranges. Rate parameters and their respective 95% likelihood-based confidence intervals of each DAG are shown in Fig. 4D for comparison. Most parameters are identifiable, except for k_in_ for DArG. We observed a trend of higher k_in_ rates with increasing acyl chain unsaturation. In case of the highly unsaturated DArG, where k_in_ is non-identifiable, the profile likelihood still indicates a lower limit of k_in_ that is larger than the upper confidence bounds of all other species. The inferred K_d_ values exhibited a similar behavior, indicating that C1 effector protein affinities to DAGs are species-specific. For example, different engineered isoforms of the C1 domain are known to have different conformational dynamics of the DAG binding site, which will affect DAG affinity and specificity [18,19].

**Figure 5.**
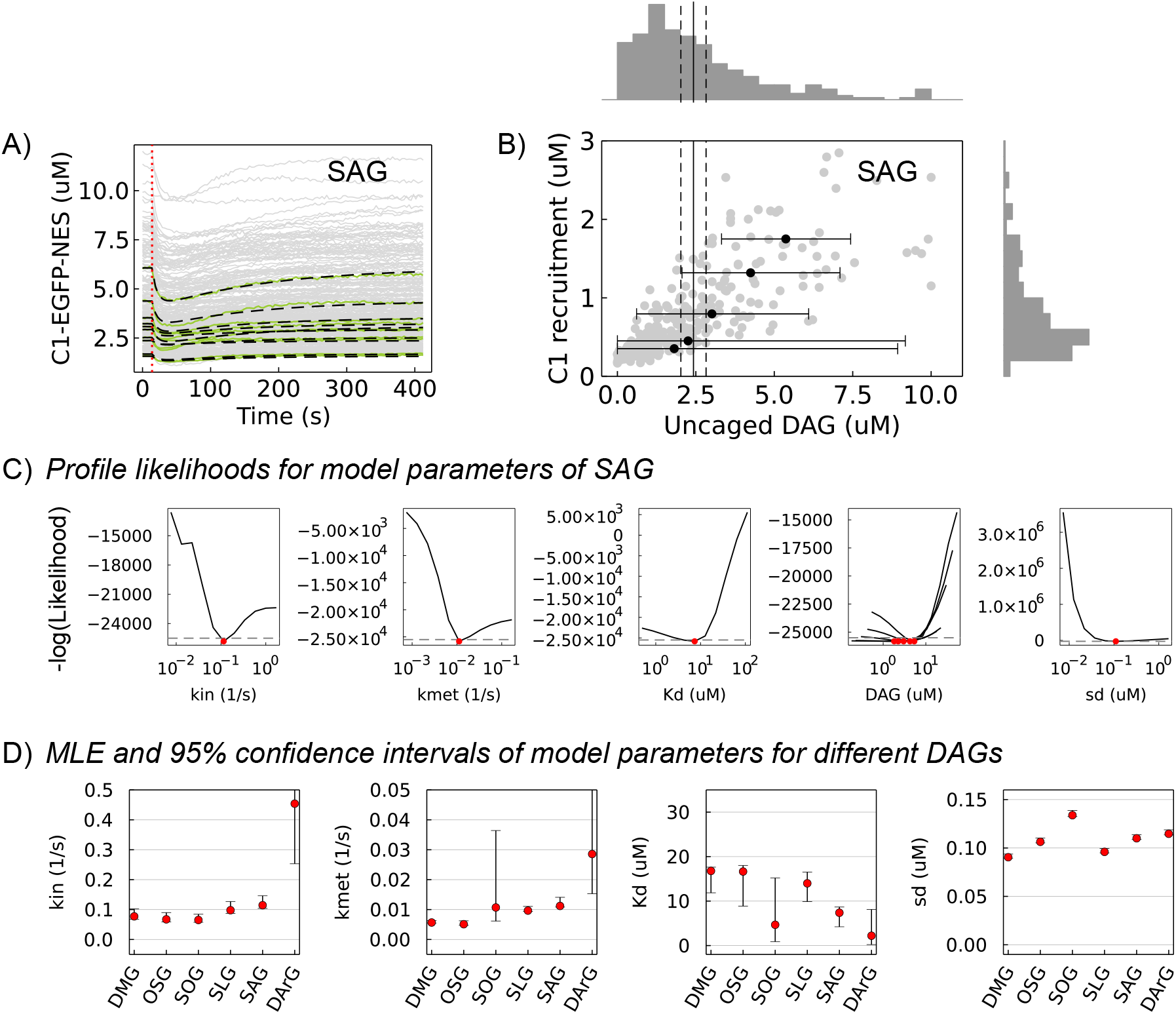
Fitting and parameter inference on single cell experimental data. (A) Single cell traces of C1-EGFP-NES dynamics (grey lines) after uncaging of SAG. Ten representative cell traces (green lines) and their respective fits (dashed black lines) are shown. Red vertical dotted line indicates the time of UV exposure that results in SAG uncaging. (B) Correlation between inferred uncaged SAG and measured C1-EGFP-NES recruitment in the cell population. Each grey dot refers to a single cell. Black dots with error bars show five representative cells and their 95% likelihood-based confidence intervals. Vertical solid and dashed lines indicate the experimentally measured mean and standard deviation of uncaged SAG. Marginal histograms of uncaged SAG and C1-EGFP-NES recruitment are shown on each axis. (C) Profile likelihoods of the model parameters. Grey horizontal dashed lines indicate the 95% likelihood-based confidence interval threshold and red circles indicate the fit MLE of the parameter. Profile likelihoods of uncaged SAG are shown for the same five representative cells as in B. (D) Inferred model parameters and 95% likelihood-based confidence intervals of the different DAG lipid species.

We further compared the inferred parameters with the results from our previous study, where we analyzed SOG and SAG using population-averaged time-traces. For SAG, we found all parameters to be in relatively good agreement with our previous study and the obtained differences can be explained by differences in the experimental design and statistical analysis. For SOG, the rate constants k_in_ and k_met_ were in good agreement with previous values, while notable differences were observed for the inferred K_d_ value (4.67 μM instead of 0.017 μM). We believe that this discrepancy mainly originates from additional nonlinearities in SOG-driven protein recruitment which are not captured by our simple model. As an example, our model considers protein binding affinities to be constant over the entire concentration range, which neglects *e.g*., coincidence detection of multiple lipids by a single protein or nano-domain formation due to changes in lipid composition. In line with this, we have previously observed in DAG titration experiments that SOG-driven protein recruitment saturated even before all cytosolic protein was bound at the membrane. This indicated that only a certain fraction of the theoretically possible lipid-protein complexes could be formed. In our previous work, we accounted for this phenomenologically by introducing a nonlinear correction for available average C1-EGFP-NES concentration into the model, which was necessary to explain the data. Using our single-cell approach, a similar correction is potentially problematic since individual traces show substantial heterogeneity in terms of initial protein concentration and recruitment. However, since initial DAG concentration is a free parameter in the single-cell model, these nonlinear effects are still captured. Specifically, it can compensate for saturation behavior in protein recruitment by lowering the amount of liberated DAG. For species, where such nonlinear effects are significant, the inferred initial DAG concentration has to be considered as an effective quantity capturing the fraction of liberated DAG, which is available for the formation of lipid-protein complexes. The distribution of inferred DAG values for SOG was indeed shifted towards lower values when compared to the other DAGs (*Fig. S12 in the Supplementary Materials*) and the inferred average liberated DAG was lower than experimental measurements (*Table S5 in Supplementary Materials*). Intriguingly, the same trend was observed for the highly unsaturated DArG but not for the regioisomer OSG. This suggests that species-specific effects may play an important role in DAG signaling.

## Discussion

In this study, we quantified kinetic parameters and lipid-protein affinities describing DAG signaling events from single cell time traces. We triggered perturbations of cellular DAG levels by DAG uncaging at the plasma membrane and monitored the formation of lipid-protein complexes by observing the recruitment of a C1-EGFP-NES reporter protein to the plasma membrane. We used a series of known and newly generated caged DAG probes in live cell photoactivation experiments to characterize the structure-activity relationships in DAG signaling processes with a particular focus on the influence of side chain unsaturation degree. We found that a higher unsaturation degree of DAGs resulted in faster outside-in rates (k_in_) and turnover rates (k_met_). Interestingly, SOG and OSG show significant differences (*i.e*. non-overlapping confidence intervals) in turnover rates (k_met_) despite being DAG isomers. DArG, having the highest degree of unsaturation in both acyl chains, showed the highest outside-in and turnover rate. A similar trend was also observed for lipid-protein affinities (with the exception of SOG), pointing to specific recognition and interaction sites between the C1 domain and the respective DAG. This is in line with recently observed structures between the C1 domain and DAG complexes that show stereospecific binding in both S*n*1 and S*n*2 positions [20].

Directly measuring single cell uncaging photoreaction yields is not possible using our previously published methodology, which requires separate experiments for photoreaction yield determination and acquisition of time trace data. To address this, we treated the initial photoreaction yield as an additional parameter that is inferred simultaneously with other parameters such as lipid-protein affinities and rates for trans-bilayer movement and turnover. We find that the resulting model can still be fully parameterized from time-trace data if the data quality is sufficient. This is remarkable because our previous population level analysis required the acquisition of dose response curves via uncaging light titrations to uniquely identify all model parameters. This seems to be the case as our current approach exploits natural protein level variations to sample the concentration space. The main determinant of parameter identifiability was found to be measurement noise. The number of cell traces used can also improve parameter identifiability, but results in more computational effort and improvements begin to diminish after a certain number of cells (N~100). This suggests that improvements in imaging technology and image analysis pipelines will directly result in improved capability for extracting quantitative parameters from live-cell time trace data.

Our analytical pipeline is based on a relatively simple model, which may not capture all relevant dynamic properties of the investigated signaling events. An example of this may be SOG-driven C1-domain recruitment, where protein recruitment appears to involve an unknown rate-limiting step [14] which is not included in the model. Similarly, we have assumed that cell-to-cell variability is predominantly due to variable photoreaction yields and C1-EGFP-NES concentrations, whereas other parameters may be subject to heterogeneity as well. However, including additional heterogeneous parameters would increase model complexity and likely impair parameter identifiability when using the current experimental data. In the future these limitations may be addressed by extending our approach to include more information-rich datasets, such as laser power titration series or more complex temporal perturbation patterns [21]. In summary, our study demonstrates how kinetic parameters of DAG-uncaging induced lipid signaling events can be quantified from single cell traces without the need to experimentally measure photoreaction yields. Such approaches will help understanding cellular information processing during lipid signaling on the single cell level.

## Supplementary materials

Materials and methods and additional results are described in the supplementary materials. Data and code used for analysis are available in the Edmond repository https://doi.org/10.17617/3.3JM5WX.

## Acknowledgments

A.N. gratefully acknowledges financial support by the European Research Council (ERC) under the European Union’s Horizon 2020 research and innovation program (grant agreement no GA 758334 ASYMMEM), by the Deutsche Forschungsgemeinschaft (DFG) via the TRR83 consortium. C.Z. and A.N. gratefully acknowledges core funding from MPI-CBG. We thank the following services and facilities at MPI-CBG Dresden for their support: Protein Expression Facility, Mass Spectrometry Facility, and the Light Microscopy Facility. We thank Jan Peychl, Britta Schroth-Diez, and Sebastian Bundschuh for the outstanding support and expert advice.

