## Supplementary Information for "Quantifying single cell lipid signaling kinetics after photo-stimulation"

**Contents**

S1. Model of DAG lipid dynamics
S2. Maximum likelihood estimation
S3. Profile likelihood analysis
S4. MLE and profile likelihoods of simulated data
S5. MLE and profile likelihoods of experimental data
S6. Loading and uncaging experiments*
S7. Image analysis and data processing*
S8. Synthesis of caged DAGs

*Sections 6 and 7 are based on methods in [1] and are partially reproduced here to improve clarity.

All data and code used for analysis are available in the Edmond repository under https://doi.org/10.17617/3.3JM5WX.

### S1. Model of DAG lipid dynamics

The kinetic model of diacylglycerol (DAG) lipid dynamics and C1-EGFP-NES protein recruitment is based on experiments and the model in [1]. Briefly, caged DAG is first loaded on the outer leaflet of the cell membrane. The caged DAG is then exposed to UV light for uncaging, freeing it to flip-flop between inner ($DAG_{int}$) and outer ($DAG_{ext}$) leaflets of the lipid bilayer. In the inner leaflet, $DAG_{int}$ can be metabolized by the cell or recruit C1-EGFP-NES (denoted as C1 in the equations) to form the DAG-C1 complex.

The rate equations of the $DAG_{ext}$ and $DAG_{int}$ species are

$$\frac{d}{dt}DAG_{ext} = -k_{in}DAG_{ext} + k_{out}DAG_{int} \quad \text{(Eqn. S1)}$$

$$\frac{d}{dt}DAG_{int} = k_{in}DAG_{ext} - k_{out}DAG_{int} - k_{met}DAG_{int} - \frac{d}{dt}DAGC1 \quad \text{(Eqn. S2)}$$

where $DAG_{int}$, $DAG_{ext}$, and DAG-C1 are in units of μmole, which is the total μmoles of a species on the membrane of a cell. The rate parameters $k_{in}$ and $k_{out}$ are the rate constants of DAG flip-flop between the inner and outer leaflet, and $k_{met}$ is the rate constant of DAG turnover in the inner leaflet. The recruitment of C1-EGFP-NES to $DAG_{int}$ is assumed to be fast and in quasi-equilibrium with equilibrium constant $K_d$

$$K_d = \frac{[C1]DAG_{int}}{DAGC1} \quad \text{(Eqn. S3)}$$

where [C1] is in μM concentration units (indicated by brackets) in the cytoplasm of the cell. Rearranging Eqn. S3 to get $DAG_{int}$,

$$DAG_{int} = \frac{K_d DAGC1}{[C1]}. \quad \text{(Eqn. S4)}$$

DAGC1 in Eqn.S4 can be replaced in terms of the difference of C1 concentration from its initial concentration

$$DAGC1 = ([C1]_0 - [C1])V_{cell} \quad \text{(Eqn. S5)}$$

where $V_{cell}$ is the volume of the cell in liters. Eqn. S4 can be rewritten as

$$DAG_{int} = K_d V_{cell}\frac{([C1]_0 - [C1])}{[C1]}. \quad \text{(Eqn. S6)}$$

Taking the derivative of Eqn. S6

$$\frac{d}{dt}DAG_{int} = \frac{d}{dt}\left(K_d V_{cell}\frac{([C1]_0 - [C1])}{[C1]}\right) \quad \text{(Eqn. S7)}$$

$$\frac{d}{dt}DAG_{int} = K_d V_{cell}\frac{d}{dt}\left(\frac{[C1]_0}{[C1]} - \frac{[C1]}{[C1]}\right) \quad \text{(Eqn. S8)}$$

$$\frac{d}{dt}DAG_{int} = K_d V_{cell}\frac{d}{dt}\left(\frac{[C1]_0}{[C1]}\right) \quad \text{(Eqn. S9)}$$

$$\frac{d}{dt}DAG_{int} = K_d V_{cell}[C1]_0\frac{d}{dt}\left(\frac{1}{[C1]}\right) \quad \text{(Eqn. S10)}$$

$$\frac{d}{dt}DAG_{int} = -\frac{K_d V_{cell}[C1]_0}{[C1]^2}\frac{d}{dt}[C1]. \quad \text{(Eqn. S11)}$$

Next, we combine the LHS of Eqn. S11 with the RHS of Eqn. S2

$$-\frac{K_d V_{cell}[C1]_0}{[C1]^2}\frac{d}{dt}[C1] = k_{in}DAG_{ext} - k_{out}DAG_{int} - k_{met}DAG_{int} - \frac{d}{dt}DAGC1 \quad \text{(Eqn. S12)}$$

$$\frac{d}{dt}[C1] = \left(k_{in}DAG_{ext} - k_{out}DAG_{int} - k_{met}DAG_{int} - \frac{d}{dt}DAGC1\right)\left(-\frac{[C1]^2}{K_d V_{cell}[C1]_0}\right) \quad \text{(Eqn. S13)}$$

In the RHS, DAGC1 can replaced using Eqn. S5 and $DAG_{int}$ can be replaced using Eqn. S6.

$$\frac{d}{dt}[C1] = \left(k_{in}DAG_{ext} - (k_{out} + k_{met})K_d V_{cell}\left(\frac{[C1]_0 - [C1]}{[C1]}\right) - \frac{d}{dt}([C1]_0 - [C1])V_{cell}\right)\left(-\frac{[C1]^2}{K_d V_{cell}[C1]_0}\right) \quad \text{(Eqn. S14)}$$

This is then rearranged and simplified accordingly

$$\frac{d}{dt}[C1] = \left(k_{in}DAG_{ext} - (k_{out} + k_{met})K_d V_{cell}\left(\frac{[C1]_0 - [C1]}{[C1]}\right) - V_{cell}\frac{d}{dt}[C1]\right)\left(-\frac{[C1]^2}{K_d V_{cell}[C1]_0}\right) \quad \text{(Eqn. S15)}$$

$$\frac{d}{dt}[C1] + \left(\frac{[C1]^2}{K_d[C1]_0}\right)\frac{d}{dt}[C1] = \left(k_{in}DAG_{ext} - (k_{out} + k_{met})K_d V_{cell}\left(\frac{[C1]_0 - [C1]}{[C1]}\right)\right)\left(-\frac{[C1]^2}{K_d V_{cell}[C1]_0}\right) \quad \text{(Eqn. S16)}$$

$$\left(1 + \frac{[C1]^2}{K_d[C1]_0}\right)\frac{d}{dt}[C1] = \left(k_{in}DAG_{ext} - (k_{out} + k_{met})K_d V_{cell}\left(\frac{[C1]_0 - [C1]}{[C1]}\right)\right)\left(-\frac{[C1]^2}{K_d V_{cell}[C1]_0}\right) \quad \text{(Eqn. S17)}$$

$$\frac{d}{dt}[C1] = \frac{\left(k_{in}DAG_{ext} - (k_{out} + k_{met})K_d V_{cell}\left(\frac{[C1]_0 - [C1]}{[C1]}\right)\right)\left(-\frac{[C1]^2}{K_d V_{cell}[C1]_0}\right)}{\left(1 + \frac{[C1]^2}{K_d[C1]_0}\right)} \quad \text{(Eqn. S18)}$$

$$\frac{d}{dt}[C1] = \frac{\left(-\frac{k_{in}DAG_{ext}[C1]^2}{K_d V_{cell}[C1]_0} + \frac{(k_{out} + k_{met})[C1]^2}{[C1]_0}\left(\frac{[C1]_0 - [C1]}{[C1]}\right)\right)}{\left(1 + \frac{[C1]^2}{K_d[C1]_0}\right)} \quad \text{(Eqn. S19)}$$

$$\frac{d}{dt}[C1] = \frac{\left(-\frac{k_{in}DAG_{ext}[C1]^2}{K_d V_{cell}[C1]_0} + \frac{(k_{out} + k_{met})[C1]^2}{[C1]_0}\left(\frac{[C1]_0 - [C1]}{[C1]}\right)\right)}{\left(1 + \frac{[C1]^2}{K_d[C1]_0}\right)}\left(\frac{\frac{K_d[C1]_0}{[C1]^2}}{\frac{K_d[C1]_0}{[C1]^2}}\right) \quad \text{(Eqn. S20)}$$

$$\frac{d}{dt}[C1] = \frac{\left(-\frac{k_{in}DAG_{ext}}{V_{cell}} + (k_{out} + k_{met})K_d\left(\frac{[C1]_0 - [C1]}{[C1]}\right)\right)}{\left(1 + \frac{K_d[C1]_0}{[C1]^2}\right)} \quad \text{(Eqn. S21)}$$

$$\frac{d}{dt}[C1] = \frac{\left(-\frac{k_{in}DAG_{ext}}{V_{cell}} + (k_{out} + k_{met})K_d\left(\frac{[C1]_0 - [C1]}{[C1]}\right)\right)}{\left(1 + \frac{K_d[C1]_0}{[C1]^2}\right)}. \quad \text{(Eqn. S22)}$$

Eqn. S22 describes the dynamics of C1-EGFP-NES as a function of the species C1-EGFP-NES and $DAG_{ext}$ only and not $DAG_{int}$. The $DAG_{int}$ term in Eqn. S1 can be replaced using Eqn. S6

$$\frac{d}{dt}DAG_{ext} = -k_{in}DAG_{ext} + k_{out}K_d V_{cell}\frac{([C1]_0 - [C1])}{[C1]}. \quad \text{(Eqn. S23)}$$

Lastly, for both Eqns. S22-23, $DAG_{ext}$ is converted into units of concentration by $[DAG_{ext}] = DAG_{ext}/V_{cell}$. This has the benefit of reducing the number of parameters by removing $V_{cell}$ in the model. The final model is then composed of

$$\frac{d}{dt}[DAG_{ext}] = -k_{in}[DAG_{ext}] + k_{out}K_d\frac{([C1]_0 - [C1])}{[C1]} \quad \text{(Eqn. S24)}$$

$$\frac{d}{dt}[C1] = \frac{\left(-k_{in}[DAG_{ext}] + (k_{out} + k_{met})K_d\left(\frac{[C1]_0 - [C1]}{[C1]}\right)\right)}{\left(\frac{K_d[C1]_0}{[C1]^2} + 1\right)}. \quad \text{(Eqn. S25)}$$

For a model with $k_{out} \rightarrow 0$, Eqns. S24-25 are reduced to

$$\frac{d}{dt}[DAG_{ext}] = -k_{in}[DAG_{ext}] \quad \text{(Eqn. S26)}$$

$$\frac{d}{dt}[C1] = \frac{\left(-k_{in}[DAG_{ext}] + k_{met}K_d\left(\frac{[C1]_0 - [C1]}{[C1]}\right)\right)}{\left(\frac{K_d[C1]_0}{[C1]^2} + 1\right)}. \quad \text{(Eqn. S27)}$$

**S2. Maximum likelihood estimation**

The agreement between the experimental data and model is measured by the natural logarithm of the probability of the experimental data $\boldsymbol{Y}$ given the model parameters $\boldsymbol{\theta}, \boldsymbol{\phi}, \sigma$ or equivalently the loglikelihood $\ell(\cdots)$ of the model parameters given the experimental data

$$\ln P(\boldsymbol{Y}|\boldsymbol{\theta}, \boldsymbol{\phi}, \sigma) = \ell(\boldsymbol{\theta}, \boldsymbol{\phi}, \sigma|\boldsymbol{Y}), \quad \text{(Eqn. S28)}$$

where $\boldsymbol{\theta}$ is a vector of the population-wide rate parameters, $\boldsymbol{\phi}$ is a vector containing the individual cell parameters that vary across the population, $\sigma$ is the standard deviation of measurement noise (denoted as sd in the tables and figures), and $\boldsymbol{Y}$ is the experimental data of single cell traces. More specifically, the population-wide parameters are the rate parameters of the model $\boldsymbol{\theta} = \{k_{in}, k_{out}, k_{met}, K_D\}$ and the individual cell parameters are the uncaged DAG concentrations of each cell $\boldsymbol{\phi} = \{[DAG]_{0,1}, [DAG]_{0,2}, \cdots, [DAG]_{0,N}\}$, where $N$ is the total number of cells. The best fit parameters of the model of lipid signaling dynamics were determined by maximum likelihood estimation (MLE). Practically, we minimized the negative joint loglikelihood of the model parameters given the experimental data

$$(\hat{\boldsymbol{\theta}}, \hat{\boldsymbol{\phi}}, \hat{\sigma}) = \underset{\boldsymbol{\theta},\boldsymbol{\phi},\sigma}{\operatorname{argmin}}\{-\ell(\boldsymbol{\theta}, \boldsymbol{\phi}, \sigma|\boldsymbol{Y})\}. \quad \text{(Eqn. S29)}$$

The total number of parameters scale with the cell population size, which results in a high dimensional optimization problem. Since optimizing all parameters at once is computationally very demanding, we instead used a heuristic approach that iterates between optimizing the population-wide parameters $\boldsymbol{\theta}$ and $\sigma$, and individual cell parameters $\boldsymbol{\phi}$. However, this approach tends to get stuck in local minima close to the initial values of population-wide parameters or individual cell parameters, depending on which was initialized first during the iteration. To address this problem, we additionally explore different initial conditions that results in the lowest negative loglikelihood. To do this, we took advantage of the observation that C1-EGFP-NES recruitment is positively correlated to uncaged DAG concentrations. This means that the uncaged DAG of individual cells can be roughly estimated as a linear function of their recruited C1-EGFP-NES

$$[\boldsymbol{DAG}]_{\boldsymbol{0}} \cong r\Delta[\boldsymbol{C1}]_{\boldsymbol{max}}, \quad \text{(Eqn. S30)}$$

where $[\boldsymbol{DAG}]_{\boldsymbol{0}}$ is the uncaged DAG, $\Delta[\boldsymbol{C1}]_{\boldsymbol{max}}$ is the maximum measured C1-EGFP-NES recruited from initial concentrations, and $r$ is typically between 0 to 1. This allows us to initialize uncaged DAG concentrations for each cell in the population by setting a single value $r$. Initializing $r$ instead of all individual cell parameters allows us to more easily explore the parameter space at different initial conditions. To efficiently scan different initial conditions and optimize the parameters, we used a bilevel optimization approach [2] consisting of two nested upper and lower optimization steps. The upper level is an optimization of the value of $r$ to find the best initial conditions of individual cell parameters $\boldsymbol{\phi}$ that minimize the negative loglikelihood. The lower level optimization involves minimizing the negative loglikelihood iteratively between population-wide parameters $\boldsymbol{\theta}$ and $\sigma$ and individual cell parameters $\boldsymbol{\phi}$. More specifically, the bilevel optimization problem reads as minimizing the negative loglikelihood with respect to $r$

$$\min_{r,\boldsymbol{\phi},\boldsymbol{\theta},\sigma}\{-\ell(r, \boldsymbol{\theta}, \boldsymbol{\phi}, \sigma|\boldsymbol{Y})\}$$

subject to

$$0 \le r \le 1,$$
$$(\boldsymbol{\phi}, \boldsymbol{\theta}, \sigma) \in S(r),$$

where the lower-level optimization solution $S(r)$ is given by the $r$-parameterized problem

$$\min_{\boldsymbol{\phi},\boldsymbol{\theta},\sigma}\{-\ell(r,\boldsymbol{\theta},\boldsymbol{\phi},\sigma|\boldsymbol{Y})\}.$$

The lower level optimization is further described in Algorithm S1. The values of $r$ are limited between 0-1, uncaged DAG concentrations are limited between 0-10 μM, and the rate parameters $k_{in}, k_{out}, k_{met}, K_D$ are optimized in log-space without bounds to avoid negative values. In the cases where only one individual cell trace is analyzed, parameter optimization is reduced to an MLE with only fixed parameters.

**Algorithm S1.** Lower level optimization algorithm.

Set number of iterations $p_{iter}$ and $c_{iter}$<br>
Given an $r$ value, initialize all individual cell parameters $\boldsymbol{\phi}$ by $[\boldsymbol{DAG}]_{\mathbf{0}} = \Delta[\boldsymbol{C1}]_{max}/r$<br>
**for** $i$ in $1{:}\,p_{iter}$ **do**<br>
    Get $\widehat{\boldsymbol{\theta}}, \hat{\sigma} = argmin_{\boldsymbol{\theta},\sigma}\{-\ell(\boldsymbol{\theta},\sigma|\boldsymbol{Y},\boldsymbol{\phi})\}$<br>
    Update $\boldsymbol{\theta}, \sigma$ with $\widehat{\boldsymbol{\theta}}, \hat{\sigma}$<br>
    **for** $j$ in $1{:}\,c_{iter}$ **do**<br>
        **for** $k$ in $1{:}\,N_{cells}$ **do**<br>
            Get $\hat{\phi}_k = argmin_{\boldsymbol{\phi_k}}\{-\ell(\phi_k|\boldsymbol{Y},\boldsymbol{\theta},\sigma)\}$<br>
            Update $\phi_k$ with $\hat{\phi}_k$<br>
        **end for**<br>
    **end for**<br>
**end for**

### S3. Profile likelihood analysis

Profile likelihoods are used to determine identifiability and confidence intervals of the parameter estimates [3]. The profile likelihood of a parameter $\theta_m$ is calculated by taking the minimum negative loglikelihood with respect to all parameters $\boldsymbol{\theta_{l\neq m}}$ while holding $\theta_m$ fixed

$$PL(\theta_m) = \min_{\boldsymbol{\theta_{l\neq m}}}\{-\ell(\boldsymbol{\phi},\boldsymbol{\theta},\sigma|\boldsymbol{Y})\}. \qquad \text{(Eqn. S31)}$$

The 95% likelihood-based confidence intervals are calculated by the regions

$$\left\{\theta_m \middle| PL(\theta_m) < \left(-\ell\big(\widehat{\boldsymbol{\phi}},\widehat{\boldsymbol{\theta}},\hat{\sigma}\big|\boldsymbol{Y}\big) + \chi^2(\alpha, df)\right)\right\}, \qquad \text{(Eqn. S32)}$$

where $\chi^2(\alpha, df)$ is the chi-squared distribution with $\alpha = 0.95$ confidence levels and $df$ is the degrees of freedom or number of parameters of the model [3]. Profile likelihoods of each parameter were calculated between a range greater and less than the MLE parameter value by a factor of 10 at eleven equally spaced intervals. The 95% confidence intervals were estimated by linearly interpolating the intersections of profile likelihood and the likelihood-based confidence interval threshold $-\ell\big(\widehat{\boldsymbol{\phi}},\widehat{\boldsymbol{\theta}},\hat{\sigma}\big|\boldsymbol{Y}\big) + \chi^2(\alpha, df)$ in Eqn. S32.

### S4. MLE and profile likelihoods of simulated data

Parameter identifiability of the model of DAG signalling dynamics was evaluated by looking at the profile likelihoods of each parameter fit on the simulated data of single cell trajectories. Simulated data of different levels of uncaged DAG without measurement noise (Fig. 2B and Fig. S1B) and different levels of measurement noise were tested to see if the amount of C1 recruitment or measurement noise affected parameter identifiability (Fig. 2C and Fig. S1C). Tests were performed on models of DAG signalling dynamics with $k_{out}$ (Eqns. S24-25) and without $k_{out}$ (Eqns. S26-27). For both models, increasing the amount of uncaged DAG (and consequently the recruited C1 signal) can improve identifiability as indicated by steeper profile likelihoods in the optimum values. All parameters remain identifiable even at low levels of uncaged DAG of 0.1 μM (Fig. 2D and Fig. S1D). In contrast, increasing measurement noise significantly affects parameter identifiability. Profile likelihoods quickly flatten and show non-identifiability as the standard deviation of the simulated Gaussian measurement noise is increased from 0.001 to 0.15 μM (Fig. 2E and Fig. S1E). In addition, the profile likelihoods of parameters $k_{out}$, $k_{met}$, and $K_d$ in the model with $k_{out}$ show more non-identifiabilities as compared to the simpler model without $k_{out}$. We next tested if using traces from multiple cells would help improve parameter identifiability. We simulated data from 10, 50, 100, and 200 cells and used them for parameter estimation and profile likelihood analysis as described in Section 3. Increasing the number of cells used for parameter estimation improved parameter identifiability as shown by the profile likelihoods and confidence intervals of the fit parameters (Fig. S2 & S3, Table S1). However, in the model without $k_{out}$, non-identifiable parameters $k_{out}$, $k_{met}$, and $K_d$ persisted when 100 cells were used for parameter estimation (Fig. S4). The true values were used to initialize the parameter values for MLE and profile likelihood analysis.

### S5. MLE and profile likelihoods of experimental data

Model parameters and parameter identifiability were estimated by MLE and profile likelihoods for each DAG species using all cells in the population individually as described in S6 using the model without $k_{out}$. Fitting of single cell traces, profile likelihoods, and inferred uncaged DAG concentrations for DMG, OSG, SOG, SLG, SAG, and DArG lipids are shown in Figs. S5-S12. MLE values of model parameters and their 95% likelihood-based confidence intervals are shown in Table S2. The MLE and 95% likelihood-based confidence intervals for the same datasets using population averaged or mean traces for each DAG and the measured average uncaged DAG are shown in Table S3 and Fig. S13. In [1], the effective C1 concentrations were approximately 25% and 75% of the initial measured C1 concentrations for SOG and SAG, respectively. We reanalyzed the population averaged data using this C1 correction for our SOG and SAG data as well (Table S4 and Fig. S14). Table S5 shows the comparison of MLE values from fitting using individual cells, population averaged data with average uncaged DAG measurements, population averaged data with average uncaged DAG measurements and corrected C1 concentrations for SOG and SAG, and parameter values from our previous work using population averaged data with average uncaged DAG measurements and corrected C1 concentrations [1]. To ensure that the optimization procedure described in Section 2 resulted in the minimum negative loglikelihood, 100 random parameter sets between the ranges 0-1 $s^{-1}$, 0-1 $s^{-1}$, 0-20 μM, and 0-0.5 μM for $k_{in}$, $k_{met}$, $K_d$, and $\sigma$, respectively, were used to initialize the optimization (Figs. S15-17). The parameters with the lowest resulting negative loglikelihood was subsequently used as initial conditions for MLE and profile likelihood analysis. Residuals of the fit for each DAG using single cell data are shown in Fig. S18.

### S6. Loading and uncaging experiments

*Loading of caged DAGs*

Cells were imaged in eight-well Lab-Tek microscope dishes (155411, Thermo Scientific) for laser-scanning confocal microscopy (Olympus Fluoview 1000) at 37 °C and 5% CO2 in imaging buffer containing (20 mM HEPES, 115 mM NaCl, 1.2 mM $CaCl_2$, 1.2 mM $MgCl_2$, 1.2 mM $K_2HPO_4$, 10 mM glucose). Cells were placed into imaging buffer 30 min prior to imaging. To load the caged DAGs, DMSO stocks were diluted respectively to reach final loading concentrations (cgSAG 602 μM, cgSOG 563 μM, cg1,3DOG 348 μM, cgOSG 585 μM, cgSLG 530 μM, cgDMG 256 μM and cgDArG 292 μM) with imaging buffer containing 0.05% pluronic and added to the cells. The loading solution was removed after 5 min incubation at room temperature and the cells subsequently washed three times with imaging buffer and then placed into a climate chamber at the microscope to equilibrate for 5 min. Loading concentrations were optimized for each caged DAG to give comparable plasma membrane fluorescence intensity of the coumarin cage (Fig. 3 and Fig. S19).

*Laser scanning confocal microscopy*

To monitor the translocation of the C1-EGFP-NES, cells were transfected and cgDAGs were loaded as described above. Imaging was performed on a dual scanner confocal microscope Olympus Fluoview 1000, with an Olympus UPlanSApochromat 60x 1.35 Oil objective. Microscope settings were adjusted to generate images displaying background fluorescence values slightly larger than zero in order to capture the complete signal stemming from the respective fluorescent dyes or proteins. Coumarin dyes were excited with the 405 nm laser and emitted light was collected between 425 and 475 nm. C1-EGFP-NES was excited with a 488 nm laser and emitted light was collected at 500–550 nm. Images were acquired with the software FV10-ASW 1.7. For quantitative datasets used for modeling, the following software settings were used: 2.5% laser power 488 nm, 580 V HV, 4x gain and 9% offset). For quantitative datasets used for modeling including cell volume and surface, the following software settings were used: 3.5% laser power 488 nm, 680 V HV, 4x gain and 10% offset).

Uncaging experiments always comprised the imaging of the protein for a defined number of frames prior to uncaging to establish a baseline, then the uncaging step and finally monitoring of the effect of the uncaging on the protein localization over time. Uncaging was triggered by scanning one frame with the 405 nm laser using 10% laser power for all experiment and a pixel dwell time of 4 µs. Uncaging experiments were carried out for no longer than an hour after loading. All life cell uncaging experiments were performed at least in biological triplicates, each comprising 4-6 technical replicates to ensure highly significant measurements. This typically resulted in datasets of 50-200 single cell traces per condition.

### S7. Image analysis and data processing

*Calibration curve with purified DAG sensor C1-EGFP-NES*

Solutions of purified DAG sensor (C1-EGFP-NES) [1] at different concentration were imaged using the same settings as for live cell microscopy on the dual scanner confocal microscope Olympus Fluoview 1000, with an

Olympus UPlanSApochromat 60x 1.35 Oil objective. Care was taken to image in homogenous solution and several images were taken per condition. The calibration curve was generated by plotting the mean fluorescence intensity of the whole field of view fluorescence intensity against the protein concentration. A linear fit resulted in the following equation y=273.6µM*x+4.9.

*Image quantification*
Time course experiments were analyzed using a custom-made Python script provided in the Supplementary Materials. To segment the cells, the sum of all the frames was used. This resulting image was then segmented using Cellpose v0.0.2.5 (www.cellpose.org) [4]. After segmentation, the nucleus of each cell was removed by using a Yen threshold on each segmented cell. Finally, cell segmentation was adjusted for cell movements during the course of the experiment by performing a Li threshold on every frame and using the regions that overlapped with the mask obtained previously.

*Quantification of the efficiency of the photo-reaction in living cells*
DAG photorelease was quantified using the methodology in [1] and the experimental details are reproduced here to provide the reader with a clear description of the utilized approaches. To quantify the efficiency of the DAG photo-release from cgDAGs at the plasma membrane, the compounds were loaded as described above and kept at 37°C for 40 min to allow for endocytosis. Whole field of view uncaging experiments were carried out while monitoring coumarin fluorescence at the lowest laser intensity (0.1% 405 nm laser power). Uncaging took place after 3 frames with a 405 nm laser intensity at 10% laser power using a frame rate of 0.14 Hz. The resulting photo-reaction released the cleaved coumarin alcohol into the extracellular space from the outer leaflet of the plasma membrane and into the endosomal lumen, respectively. Uncaging experiments were performed at least in biological triplicates, each comprising multiple technical replicates to ensure highly accurate measurements. The acquired datasets were then analyzed using a Fiji tool as described in [1] for measuring fluorescence intensity (FI) at the plasma membrane. Normalized plasma membrane FI values before ($FI_{before}$) and after uncaging ($FI_{after}$) were determined for every cell and the uncaging efficiency E was calculated according to the following equation.

$$E = 1 - \frac{FI_{after}}{FI_{before}} \qquad \text{(Eqn. S33)}$$

In order to correct for stable background fluorescence intensity unaffected by the UV irradiation at the plasma membrane, we also performed uncaging experiments featuring multiple uncaging steps (1 to 15 times) with 1% laser power for each caged DAG. The resulting data for E exhibited an asymptotic behavior. We determined the asymptotic value ($FI_{base}$) indicative of the background fluorescence by fitting a monoexponential decay function to the data (Fig. S20G). The baseline corrected uncaging efficiency $E_{corr}$ is given by the following equation.

$$E_{corr} = 1 - \frac{norm.FI_{after} - FI_{base}}{1 - FI_{base}} \qquad \text{(Eqn. S34)}$$

The error of $E_{corr}$ ($\Delta E_{corr}$) was calculated according to the general equation for error propagation. The errors for $FI_{after}$ and $FI_{base}$ were determined either as standard error ($FI_{after}$) or extracted from the monoexponential fit ($FI_{base}$).

$$\Delta E_{corr} = \left|\frac{\partial E_{corr}}{\partial FI_{after}}\right| \Delta FI_{after} + \left|\frac{\partial E_{corr}}{\partial FI_{base}}\right| \Delta FI_{base} \qquad \text{(Eqn. S35)}$$

In the current case, this results in the following equation:

$$\Delta E_{corr} = \left|\frac{1}{1 - FI_{base}}\right| \Delta norm.FI_{after} + \left|\frac{(FI_{base} - 1) - (FI_{base} - norm.FI_{after})}{(1 - FI_{base})^2}\right| \Delta FI_{base} \qquad \text{(Eqn. S36)}$$

The resulting values are summarized in Fig. S20D.

### S8. Synthesis of caged DAGs

*General synthetic procedures*

All chemicals were purchased from commercial sources (Roth, AppliChem, Arcos Organics, Biozol, Sigma, chem.impex, Otto Nordwald, Chemodex. Alfa Aesar or Merck) and were used without further purification. Solvents for flash chromatography were obtained from VWR and dry solvents were obtained from Sigma. Deuterated solvents were obtained from Deutero GmbH, Karlsruhe, Germany. All reactions were carried out using dry solvents under an inert atmosphere unless otherwise stated in the respective experimental procedure. TLC was performed on precoated silica plates (Merck, 60 $F_{254}$) using UV light (254 or 366 nm) or a solution of phosphomolybdic acid in EtOH (3% phosphomolybdic acid in 100 mL EtOH) for analysis. Preparative column chromatography was performed with 40-63 µm silica gel from VWR Chemicals with a pressure of 1-1.5 bar. HPLC purification was performed on a Knauer HPLC (pump: AZURA P2.1S, UV/VIS detector: AZURA UVD 2.1S, Evaporative Light-Scattering Detector: SEDEX LT-ELSD Model 85LT). $^{1}$H and $^{13}$C-NMR-spectra were obtained on a 400 MHz Bruker Ascend$^{TM}$ Nanobay spectrometer. J values are given in Hz and chemical shifts in ppm. Splitting patterns are designated as follows: s, singlet; d, doublet; t, triplet; q, quartett; m, multiplet; b, broad. $^{13}$C-NMR-spectra were broadband hydrogen decoupled. Mass spectra (ESI) were recorded using a QExactive instrument (Thermo Fisher Scientific) equipped with a robotic nanoflow ion source. FTMS spectra were acquired within the range of m/z 200-1200 with the target mass resolution of m/z 200=140000. The spectra were evaluated using the Xcalibur Qual Browser software.

We describe the synthesis of the different DAGs and caged DAGs below. Intermediates and final DAG products are labelled with numbers in parentheses in bold font. An overview of procedure is illustrated in Fig. S5. The sulfonated caging group, caged SOG, SAG, and 1,3-DOG were synthesized as previously described in [1], SLG **(20)** was prepared as described in [5] and the preparation of 3-O-p-Methoxybenzyl-sn-glycerol **(1)** according to [6]. **(7)** was synthesized according to [7]. An overview of the synthesis of DAGs is illustrated in Fig. S21. NMR spectra of each product is shown in Figs. S22-31.

*Synthesis of DMG and DArG*

General procedure for the synthesis of **(2)** and **(3)**

The respective fatty acid (1.19-1.58 g, 5.19 mmol, 2.2 eq.), EDC (1.086 g, 5.66 mmol, 2.4 eq.) and DMAP (29.3 mg, 0.24 mmol, 0.1 eq.) were dissolved in 20 ml dry DCE and stirred for 15 min before the addition of 500mg of 1. The turbid solution turned clear immediately and was stirred at room temperature overnight. The reaction mixture was subsequently poured into a separation funnel charged with 400 ml of $H_2O$/EtOAc (1:1). the layers were separated, the organic layer washed with $H_2O$ (1 x 100 mL) and sat. NaCl solution (1x100 mL) and dried over $Na_2SO_4$. The solvent was removed under reduced pressure and the crude product mixture was purified using FC (eluent: EtOAc/CH: 1/9). The product was obtained as a colorless oil.

**(2)**

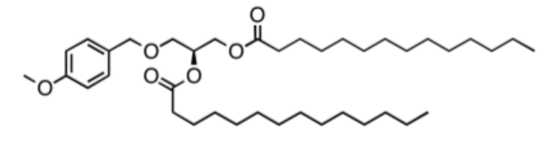

Yield: 1.29 g, 2.00 mmol (86%)

$^{1}$H NMR (400 MHz, Chloroform-d) δ 7.25 – 7.19 (m, 2H), 6.91 – 6.84 (m, 2H), 5.27 – 5.17 (mc, 1H), 4.53 – 4.40 (mc, 2H), 4.32 (dd, J = 11.9, 3.8 Hz, 1H), 4.17 (dd, J = 11.9, 6.5 Hz, 1H), 3.80 (s, 3H), 3.59 – 3.52 (m, 2H), 2.35 – 2.23 (m, 4H), 1.68 – 1.57 (m, 4H) superimposed with water, 1.32 – 1.23 (m, 40H), 0.88 (t, J = 6.7Hz, 6H).

$^{13}$C NMR (101 MHz, CDCl3) δ 129.44, 113.98, 77.48, 77.16, 76.84, 73.12, 70.19, 68.08, 55.42, 34.29, 32.08, 29.85, 29.82, 29.66, 29.52, 29.45, 29.30, 27.08, 25.13, 22.85, 14.27. Signals around 29.8-29.1 are partially not resolved due to very similar $^{13}$C chemical shifts of $CH_2$ groups.

HRMS calculated for $[M+NH_4]^+$: 650.54 Da, found: $[M+NH_4]^+$: 650.532 Da.

**(3)**

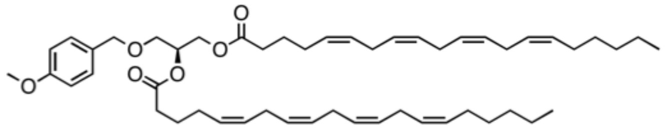

Yield: 0.87 g, 1.10 mmol (47%)

$^{1}$H NMR (400 MHz, Chloroform-d) δ 7.26 – 7.19 (m, 2H), 6.90 – 6.83 (m, 2H), 5.46 – 5.27 (m, 16H), 5.22 (mc, 1H), 4.52 – 4.40 (mc, 2H), 4.33 (dd, J = 11.9, 3.8 Hz, 1H), 4.22 – 4.04 (m, 2H), 3.80 (s, 3H), 3.59 – 3.50 (m, 2H),

2.90 – 2.76 (m, 12H), 2.38 – 2.25 (m, 4H), 2.16 – 2.00 (m, 8H), 1.76 – 1.61 (m, 4H), 1.41 – 1.22 (m, 12H), 0.93 – 0.84 (m, 6H).
$^{13}$C NMR (101 MHz, CDCl3) δ 173.27, 172.99, 159.48, 130.65, 129.89, 129.45, 129.05, 129.02, 129.01, 128.75, 128.41, 128.28, 128.00, 127.69, 113.98, 77.48, 77.16, 76.84, 73.12, 70.27, 68.02, 62.92, 60.54, 55.42, 33.86, 33.63, 31.67, 29.48, 27.38, 27.07, 26.67, 25.80, 25.78, 24.97, 24.87, 22.73, 14.22. Signals around 29.8-29.1 are partially not resolved due to very similar $^{13}$C chemical shifts of $CH_2$ groups.
HRMS calculated for $[M+NH_4]^+$: 802.60 Da, found: $[M+ NH_4]^+$: 802.595 Da.

For the synthesis of **(4)** and **(5)**, **(2)** or **(3)**, respectively (300 mg, 382-473 µmol) were dissolved in 3 ml DCM and treated with a solution of 20% TFA in DCM at 0°C (10 mL) under normal atmospheric conditions and stirred at rt. for 4 min (the clear solution turns purple). The reaction was quenched by addition of 250 ml saturated $Na_2CO_3$ solution (reaction loses the color) and transferred into a separation funnel charged with EtOAc. The layers were separated, the organic layer washed with $H_2O$ (1 x 50 mL) and sat. NaCl solution (1x50 mL) and dried over $Na_2SO_4$. The solvent was removed under reduced pressure and the residue purified by FC (eluent: cyclohexane/EtOAc 4:1). Fractions containing the product were pooled and the solvent removed under reduced pressure. The products were obtained as a white solid **(2)** or a colorless oil **(3)**.

**(4)**

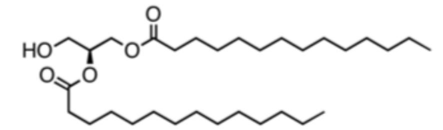


Yield: 204.1 mg, 397.9 µmol, 84%
$^{1}$H NMR (400 MHz, Chloroform-d) δ 5.12 – 5.05 (mc, 1H), 4.36 – 4.18 (m, 2H), 3.75 – 3.69 (m, 2H), 2.38 – 2.27 (m, 4H), 1.68 – 1.55 (m, 4H), 1.34 – 1.23 (m, 40H), 0.92 – 0.83 (m, 6H).
$^{13}$C NMR (101 MHz, CDCl3) δ 173.92, 173.56, 77.47, 77.15, 76.83, 72.25, 62.15, 61.70, 34.44, 34.26, 32.07, 29.83, 29.80, 29.76, 29.62, 29.50, 29.42, 29.27, 29.24, 25.09, 25.04, 22.83, 14.26. Signals around 29.8-29.1 are partially not resolved due to very similar $^{13}$C chemical shifts of $CH_2$ groups
HRMS calculated for $[M+NH_4]^+$: 530.48 Da, found: $[M+NH_4]^+$: 530.477 Da.

**(5)**

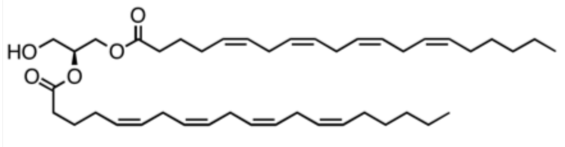


Yield: 88.7 mg, 133.3 µmol, 35%
$^{1}$H NMR (400 MHz, Chloroform-d) δ 5.46 – 5.28 (m, 16H), 5.13 – 5.03 (m, 1H), 4.37 – 4.19 (m, 2H), 3.76 – 3.69 (m, 2H), 2.87 – 2.77 (m, 12H), 2.41 – 2.30 (m, 4H), 2.17 – 2.00 (m, 8H), 1.76 – 1.64 (m, 4H), 1.42 – 1.23 (m, 12H), 0.89 (t, J = 6.6 Hz, 6H).
$^{13}$C NMR (101 MHz, CDCl3) δ 173.62, 173.26, 130.64, 129.16, 129.14, 128.89, 128.75, 128.44, 128.42, 128.22, 127.97, 127.66, 77.47, 77.15, 76.83, 72.31, 62.18, 61.65, 33.77, 33.58, 31.66, 29.46, 27.36, 26.65, 26.63, 25.78, 25.77, 25.76, 24.89, 24.85, 22.71, 14.21. Signals around 29.8-29.1 are partially not resolved due to very similar $^{13}$C chemical shifts of $CH_2$ groups.
HRMS calculated for $[M+NH_4]^+$: 682.54 Da, found: $[M+NH_4]^+$: 682.539 Da.

*Synthesis of OSG*
<u>Synthesis of **(8)**</u>
A solution of **(7)** (1.19 g, 3.00 mmol, 1 eq.) in DCE (20 mL) was subsequently treated with DIEA (2.87 mL, 16.5 mmol, 5.5 eq.) and TESOTf (0.68 mL, 3.30 mmol, 1.1). The reaction mixture was stirred at 100 °C for 1.5 h at which time an additional portion of TESOTf (400 µL) was added. After 3h once more 400 uL TESOTf was added. Stirring at 100 °C was continued for in total 3.5 and the reaction mixture allowed to reach rt. The reaction mixture was transferred onto a mixture of EtOAc and H2O (1:1, 200 mL), the layers were separated, the organic layer washed with $H_2O$ (1 x 50 mL) and sat. NaCl solution (1x50 mL) and dried over Na2SO4. The solvent was removed under reduced pressure and the residue dissolved in a 2:1 mixture of THF (50 mL) and $Na_2CO_3$ solution (10 % in $H_2O$, 25 mL). The resulting suspension was treated with iodine (1.5 g, 3.8 mmol) and stirred for 45 min. The reaction mixture was transferred onto a mixture of EtOAc and sat. $Na_2S_2O_3$ solution (1:1, 200 mL), the layers were separated, the organic layer washed with $H_2O$ (1 x 50 mL) and sat. NaCl solution (1 x 50 mL) and dried over $Na_2SO_4$. The solvent was removed under reduced pressure and the residue purified by FC (eluent: cyclohexane/EtOAc 9:1). The product was obtained as light yellow, viscous oil.

**(8)**

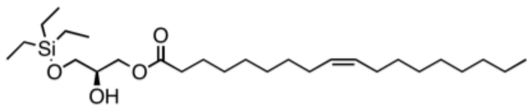

Yield: 0.80 g, 1.70 mmol, 57%

$^{1}$H NMR (400 MHz, Chloroform-d) δ 5.42 – 5.28 (mc, 2H), 4.21 – 4.04 (mc, 2H), 3.93 – 3.82 (mc, 1H), 3.67 (dd, J = 10.1, 4.6 Hz, 1H), 3.60 (dd, J = 10.1, 5.7 Hz, 1H), 2.33 (t, J = 7.6 Hz, 2H), 2.09 – 1.95 (m, 4H), 1.66 – 1.60 (m, 2H), 1.37 – 1.22 (m, 20H), 0.96 (t, 9H), 0.87 (t, 3H), 0.61 (q, J = 8.3 Hz, 6H).

$^{13}$C NMR (101 MHz, CDCl3) δ 174.07, 130.15, 129.87, 77.48, 77.16, 76.84, 70.19, 65.16, 63.52, 34.33, 32.05, 29.92, 29.85, 29.67, 29.47, 29.32, 29.27, 29.25, 27.37, 27.32, 27.06, 25.08, 22.83, 14.25, 6.81, 4.43. Signals around 29.8-29.1 are partially not resolved due to very similar $^{13}$C chemical shifts of $CH_2$ groups.

HRMS calculated for $[M+NH_4]^+$: 488.41 Da, found: $[M+NH_4]^+$: 488.411 Da.

Synthesis of **(9)**

A solution of the stearic acid (582mg, 2.04mmol, 2 eq.), EDC (558 mg, 3.06 mmol) and DMAP (12.5 mg, 10.0 µmol) in DCM (5-20mL) was stirred for 15 min at rt and subsequently treated with a solution of 8 (500 mg 1.02 mmol) in 25 mL DCM) and the reaction mixture was stirred at room temperature overnight. The reaction mixture was transferred onto a mixture of EtOAc and H2O (1:1, 100 mL), the layers were separated, the organic layer washed with $H_2O$ (1 x 50 mL) and sat. NaCl solution (1 x 50 mL) and dried over $Na_2SO_4$. The solvent was removed under reduced pressure and the residue purified by FC (eluent: cyclohexane/EtOAc 9:1). The title compound was obtained as a colourless oil.

**(9)**

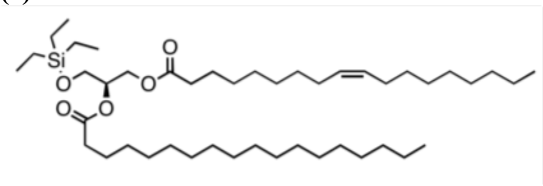

Yield: 622 mg, 0.84 mmol (82%)

$^{1}$H NMR (400 MHz, Chloroform-d) δ 5.42 – 5.29 (mc, 2H), 5.12 – 5.03 (mc, 1H), 4.35 (dd, J = 11.8, 3.7 Hz, 1H), 4.16 (dd, J = 11.8, 6.3 Hz, 1H), 3.76 – 3.67 (m, 2H), 2.36 – 2.25 (mc, 4H), 2.10 – 1.96 (m, 4H), 1.67 – 1.56 (m, 4H) superimposed with water, 1.37 – 1.23 (m, 48H), 0.95 (t, J = 7.9 Hz, 9H), 0.88 (t, 6H), 0.59 (q, J = 7.9 Hz, 6H).

$^{13}$C NMR (101 MHz, CDCl3) δ 173.60, 173.29, 130.15, 129.87, 77.48, 77.16, 76.84, 71.87, 62.64, 61.38, 34.51, 34.31, 32.09, 32.07, 29.93, 29.86, 29.82, 29.80, 29.69, 29.65, 29.52, 29.48, 29.46, 29.36, 29.29, 29.27, 27.38, 27.34, 25.11, 25.06, 22.85, 14.27, 6.81, 4.45. Signals around 29.8-29.1 are partially not resolved due to very similar $^{13}$C chemical shifts of $CH_2$ groups.

HRMS calculated for $[M+NH_4]^+$: 754.67 Da, found: $[M+NH_4]^+$: 754.670 Da.

Synthesis of **(10)**

**(9)** (200 mg, 217 µmol) was added to a 5 mM solution of $FeCl_3$ x 6 $H_2O$ in MeOH/DCM (3:1, 10 mL) under normal atmospheric conditions and stirred at rt. for 45 min. The reaction mixture was transferred onto a mixture of EtOAc and $H_2O$ (1:1, 100 mL), the layers were separated, the organic layer washed with $H_2O$ (1 x 50 mL) and sat. NaCl solution (1 x 50 mL). The solvent was removed under reduced pressure and the residue purified by FC (eluent: cyclohexane/EtOAc 3:1). Fractions containing product were pooled and the solvents removed under reduced pressure. The product was obtained as a colourless oil.

**(10)**

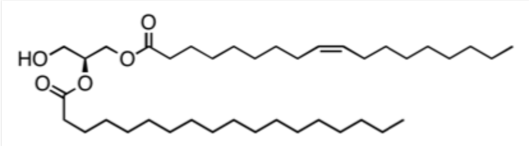

Yield: 154.9 mg, 249 µmol, 92%

$^{1}$H NMR (400 MHz, Chloroform-d) δ 5.41 – 5.29 (mc, 2H), 5.13 – 5.04 (mc, 1H), 4.36 – 4.18 (m, 2H), 3.76 – 3.69 (m, 2H), 2.38 – 2.27 (mc, 4H), 2.09 – 1.96 (m, 5H), 1.67 – 1.55 (m, 4H), 1.37 – 1.23 (m, 48H), 0.88 (t, J = 6.6 Hz, 6H).

$^{13}$C NMR (101 MHz, CDCl3) δ 173.88, 173.55, 130.36, 130.16, 130.13, 129.83, 77.47, 77.15, 76.83, 72.25, 62.14, 61.70, 34.43, 34.25, 34.23, 32.07, 32.05, 31.67, 29.91, 29.84, 29.80, 29.77, 29.67, 29.62, 29.51, 29.46, 29.42,

29.31, 29.25, 29.24, 27.37, 27.31, 25.09, 25.04, 25.02, 22.83, 14.25. Signals around 29.8-29.1 are partially not resolved due to very similar $^{13}C$ chemical shifts of $CH_2$ groups.
HRMS calculated for $[M+NH_4]^+$: 640.59 Da, found: $[M+NH_4]^+$: 640.585 Da.

*Synthesis of caged DAGs* ***(22a-d)***
The respective diacylglycerol (33.6- 46.7 mg, 65.6 µmol, 1 eq.), was dissolved in 1.5 mL of dry THF under an inert atmosphere and subsequently treated with DIPEA (20 µL, 196.8 µmol, 3 eq). The mixture was placed in an ice bath and stirred for 10 min before phosgene solution (15% in toluene) (85,72 µL, 65.6 µmol, 1 eq.) was added. The reaction mixture was stirred for 2 h in the ice bath and was then transferred onto a mixture of EtOAc and $H_2O$ (1:1, 100 mL), the layers were separated, the organic layer washed with H2O (1 x 50 mL) and sat. NaCl solution (1 x 50 mL) and dried over $Na_2SO_4$. The solvents were removed under reduced pressure.
The sulfonated caging group (40 mg, 65.6 µmol, 1 eq.) was dissolved in 1 mL of dry DCM and the resulting solution added to 100 mg of freshly activated molecular sieves (3Å) under an inert atmosphere. After addition of DIPEA (12.89 µL, 131.2 µmol, 2 eq.), the mixture was stirred for 5 min in an ice bath. Subsequently the dried reaction mixture from before (21a-d) was added in 1.5 mL dry DCM. The reaction mixture was then stirred overnight in a thawing ice bath. The molecular sieve was removed either by filtration or centrifugation and the solvents were removed under reduced pressure. The residue was purified by FC (eluent: 65% $CHCl_3$, 35% MeOH, 0.5% $H_2O$) to give a yellow solid, which was washed twice with EtOAc (50 mL) to remove residual free diacylglycerol. The products were obtained as a yellow solid.

**(22a)**

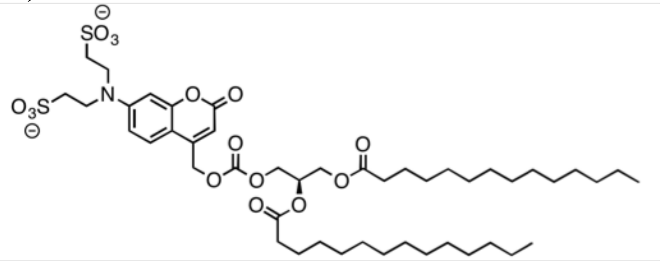

Yield: 27.8 mg, 29.4 µmol, 45%
$^1$H NMR (400 MHz, DMSO-d6) δ 7.51 (d, J = 9.0 Hz, 1H), 6.70 (dd, J = 9.0, 2.5 Hz, 1H), 6.54 (d, J = 2.5 Hz, 1H), 5.96 (s, 1H), 5.37 (s, 2H), 5.30 – 5.22 (mc, 1H), 4.45 – 4.25 (m, 3H), 4.20 – 4.10 (m, 1H), 3.69 – 3.60 (mc, 4H), 2.75 – 2.66 (mc, 4H), 2.32 – 2.21 (m, 4H), 1.54 – 1.43 (m, 4H), 1.33 – 1.13 (m, 40H), 0.88 – 0.79 (mc, 6H).
$^{13}$C NMR (101 MHz, DMSO) δ 172.46, 172.20, 160.31, 155.62, 153.80, 150.36, 149.76, 125.56, 108.74, 105.49, 97.11, 68.56, 66.18, 64.73, 61.61, 47.91, 47.42, 40.19, 40.15, 39.94, 39.73, 39.52, 39.31, 39.10, 38.89, 33.44, 33.33, 31.26, 29.03, 28.98, 28.86, 28.68, 28.36, 28.26, 24.38, 24.36, 22.06, 13.90. Signals around 29.8-29.1 are partially not resolved due to very similar $^{13}C$ chemical shifts of $CH_2$ groups.
HRMS calculated for $[M+2NH_4+H]^+$: 980.52 Da, found: $[M+2NH_4+H]^+$: 980.521 Da.

**(22b)**

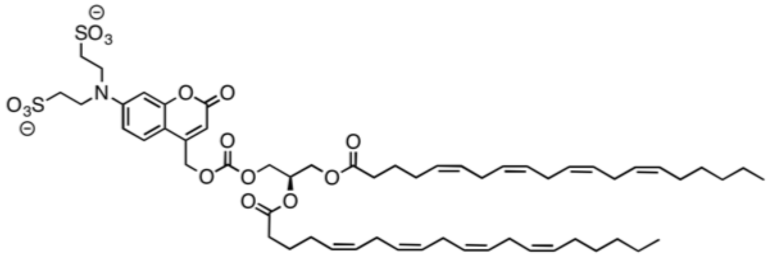

Yield: 29.8 mg, 27.1 µmol, 41%
$^1$H NMR (400 MHz, DMSO-d6) δ 7.50 (d, J = 9.1 Hz, 1H), 6.70 (dd, J = 9.0, 2.5 Hz, 1H), 6.54 (d, J = 2.5 Hz, 1H), 5.96 (s, 1H), 5.43 – 5.20 (m, 19H), 4.45 – 4.11 (m, 4H), 3.70 – 3.61 (mc, 4H), 2.83 – 2.68 (m, 16H), 2.33 – 2.22 (mc, 4H), 2.09 – 1.95 (m, 8H), 1.63 – 1.50 (mc, 4H), 1.36 – 1.17 (m, 12H), 0.88 – 0.79 (mc, 6H).
$^{13}$C NMR (101 MHz, DMSO) δ 172.33, 172.07, 160.32, 155.63, 153.81, 150.36, 149.72, 129.88, 128.84, 128.81, 128.39, 128.07, 128.05, 127.86, 127.78, 127.61, 127.46, 125.58, 108.75, 105.53, 97.13, 68.60, 66.15, 64.74, 61.69, 47.94, 47.40, 40.19, 40.15, 39.98, 39.94, 39.73, 39.52, 39.31, 39.10, 38.89, 32.83, 32.70, 30.85, 28.68, 26.58, 25.92, 25.85, 25.18, 25.16, 25.12, 24.30, 24.26, 21.94, 13.87. Signals around 29.8-29.1 are partially not resolved due to very similar $^{13}C$ chemical shifts of $CH_2$ groups.
HRMS calculated for $[M+3H]^+$: 1098.53 Da, found: $[M+3H]^+$: 1098.530 Da.

**(22c)**

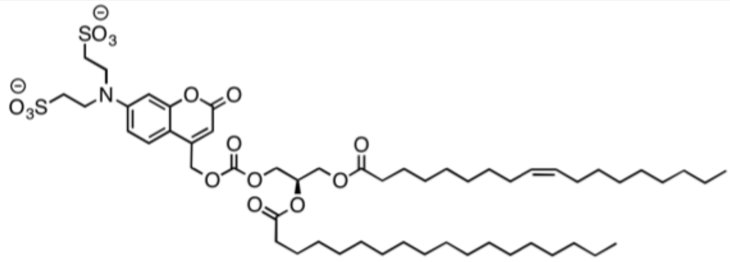

Yield: 26.6 mg, 25.2 μmol, 38%

$^1$H NMR (400 MHz, DMSO-d6) δ 7.51 (d, J = 9.0 Hz, 1H), 6.70 (dd, J = 9.0, 2.5 Hz, 1H), 6.54 (d, J = 2.4 Hz, 1H), 5.96 (s, 1H), 5.39 – 5.22 (m, 5H), 4.44 – 4.25 (m, 3H), 4.19 – 4.10 (m, 1H), 3.70 – 3.61 (mc, 4H), 2.76 – 2.66 (mc, 4H), 2.32 – 2.20 (m, 4H), 2.03 – 1.91 (m, 4H), 1.54 – 1.45 (m, 4H), 1.32 – 1.15 (m, 48H), 0.88 – 0.80 (m, 6H).

$^{13}$C NMR (101 MHz, DMSO) δ 172.41, 172.19, 160.31, 155.63, 153.81, 150.37, 149.76, 129.59, 129.51, 125.58, 108.75, 105.52, 97.13, 68.57, 66.18, 64.74, 61.64, 47.94, 47.40, 40.20, 40.15, 39.99, 39.94, 39.73, 39.52, 39.31, 39.10, 38.90, 33.44, 33.32, 31.27, 29.07, 28.99, 28.85, 28.81, 28.67, 28.57, 28.54, 28.46, 28.38, 28.26, 26.54, 24.39, 24.35, 22.07, 13.91, 13.89. Signals around 29.8-29.1 are partially not resolved due to very similar $^{13}$C chemical shifts of $CH_2$ groups.

HRMS calculated for $[M+2NH_4+H]^+$: 1090.63 Da, found: $[M+2NH_4+H]^+$: 1090.629 Da.

**(22d)**

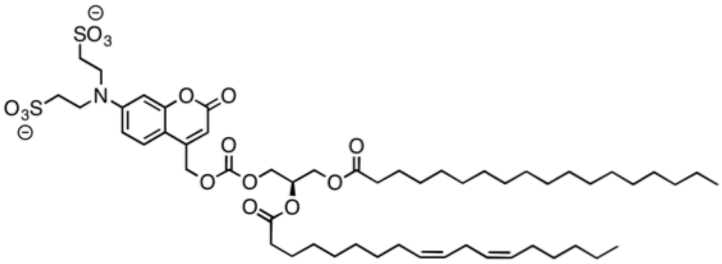

Yield: 32.5 mg, 30.8 μmol, 47 %

$^1$H NMR (400 MHz, DMSO-d6) δ 7.51 (d, J = 9.0 Hz, 1H), 6.70 (dd, J = 9.0, 2.4 Hz, 1H), 6.54 (d, J = 2.4 Hz, 1H), 5.96 (s, 1H), 5.39 – 5.21 (m, 7H), 4.44 – 4.25 (m, 3H), 4.19 – 4.10 (m, 1H), 3.70 – 3.61 (mc, 4H), 2.77 – 2.67 (m, 6H), 2.32 – 2.23 (mc, 4H), 2.04 – 1.94 (m, 4H), 1.54 – 1.45 (m, 4H), 1.34 – 1.13 (m, 42H), 0.84 (t, J = 6.6 Hz, 6H).

$^{13}$C NMR (101 MHz, DMSO) δ 172.45, 172.17, 160.31, 155.63, 153.81, 150.37, 149.76, 129.66, 129.60, 127.70, 127.68, 125.58, 108.75, 105.52, 97.14, 68.57, 66.18, 64.74, 61.63, 47.94, 47.39, 40.19, 40.15, 39.99, 39.94, 39.73, 39.52, 39.31, 39.10, 38.90, 33.43, 33.33, 31.27, 30.88, 28.99, 28.96, 28.85, 28.70, 28.67, 28.53, 28.49, 28.36, 28.28, 26.58, 25.17, 24.36, 22.07, 21.94, 13.92, 13.87. Signals around 29.8-29.1 are partially not resolved due to very similar $^{13}$C chemical shifts of $CH_2$ groups.

HRMS calculated for $[M+2NH_4+H]^+$: 1088.61 Da, found: $[M+2NH_4+H]^+$: 1088.614 Da.

**Table S1. MLE and 95% likelihood-based confidence intervals (CI) of fit parameters using different cell population sizes for model without $k_{out}$.** Plot of fit and profile likelihoods are shown in Fig. 2 in the main text and Fig. S2. The "-" denotes that the lower or upper bound threshold was not reached and the parameter is non-identifiable within the range tested.

| Parameter | 95% CI | 1 cell | 10 cells | 50 cells | 100 cells | 200 cells | True value |
|---|---|---|---|---|---|---|---|
| $k_{in}$ (1/s) | lower | 0.0208 | 0.088 | 0.091 | 0.092 | 0.092 | 0.098 |
| | upper | 0.4034 | 0.143 | 0.117 | 0.117 | 0.116 | |
| | MLE | 0.0674 | 0.106 | 0.099 | 0.100 | 0.100 | |
| $k_{met}$ (1/s) | lower | 0.0041 | 0.006 | 0.007 | 0.009 | 0.009 | 0.01823 |
| | upper | 0.0987 | 0.022 | 0.053 | 0.045 | 0.077 | |
| | MLE | 0.0207 | 0.008 | 0.012 | 0.012 | 0.015 | |
| $K_d$ (μM) | lower | - | 0.863 | 0.284 | 0.288 | 0.164 | 0.866 |
| | upper | 4.3161 | - | 11.572 | 9.807 | 5.114 | |
| | MLE | 0.8967 | 4.389 | 1.777 | 1.651 | 1.183 | |
| sd (μM) | lower | 0.0908 | 0.099 | 0.099 | 0.099 | 0.099 | 0.1 |
| | upper | 0.1454 | 0.104 | 0.102 | 0.102 | 0.102 | |
| | MLE | 0.1057 | 0.102 | 0.100 | 0.100 | 0.101 | |
| $DAG_1$ (μM) | lower | 0.5438 | 0.987 | 0.720 | 0.666 | 0.523 | 1.345 |
| | upper | - | 3.303 | 3.038 | 3.118 | 2.879 | |
| | MLE | 1.0053 | 2.310 | 1.394 | 1.348 | 1.175 | |
| $DAG_2$ (μM) | lower | | 1.186 | 0.878 | 0.790 | 0.560 | 1.147 |
| | upper | | 4.298 | 4.103 | 4.254 | 3.933 | |
| | MLE | | 3.018 | 1.756 | 1.693 | 1.450 | |
| $DAG_3$ (μM) | lower | | 0.905 | 0.686 | 0.638 | 0.483 | 1.072 |
| | upper | | 2.705 | 2.279 | 2.453 | 2.314 | |
| | MLE | | 1.917 | 1.191 | 1.155 | 1.018 | |
| $DAG_4$ (μM) | lower | | 1.267 | 1.100 | 1.022 | 0.783 | 1.267 |
| | upper | | 4.761 | 3.617 | 3.731 | 3.619 | |
| | MLE | | 3.060 | 1.846 | 1.785 | 1.556 | |
| $DAG_5$ (μM) | lower | | 1.394 | 1.136 | 1.084 | 0.893 | 1.144 |
| | upper | | 5.380 | 3.538 | 3.645 | 3.567 | |
| | MLE | | 3.114 | 1.891 | 1.830 | 1.599 | |

**Table S2. MLE and 95% likelihood-based confidence intervals (CI) of fit parameters and five representative cells for uncaged DAG from experimental data using the model without $k_{out}$.** The "-" denotes that the lower or upper bound threshold was not reached and the parameter is non-identifiable within the range tested. Number of cells analyzed are 97,105, 86, 108, 273, and 120 for DMG, OSG, SOG, SLG, SAG, and DArG, respectively.

| Parameter | 95% CI | DMG | OSG | SOG | SLG | SAG | DArG |
|---|---|---|---|---|---|---|---|
| $k_{in}$ (1/s) | lower | 0.0678 | 0.0590 | 0.0576 | 0.0872 | 0.1039 | 0.2532 |
| | upper | 0.1022 | 0.0899 | 0.0844 | 0.1270 | 0.1460 | - |
| | MLE | 0.0773 | 0.0674 | 0.0652 | 0.0979 | 0.1146 | 0.4540 |
| $k_{met}$ (1/s) | lower | 0.0055 | 0.0048 | 0.0062 | 0.0094 | 0.0108 | 0.0153 |
| | upper | 0.0063 | 0.0062 | 0.0364 | 0.0111 | 0.0141 | 0.2119 |
| | MLE | 0.0057 | 0.0051 | 0.0107 | 0.0097 | 0.0112 | 0.0286 |
| $K_d$ (μM) | lower | 11.8646 | 8.8763 | 0.8620 | 9.8740 | 4.2203 | 0.2207 |
| | upper | 17.6314 | 18.0123 | 15.2129 | 16.5275 | 8.6616 | 8.1153 |
| | MLE | 16.7766 | 16.6422 | 4.6735 | 13.9923 | 7.3750 | 2.2068 |
| sd (μM) | lower | 0.0895 | 0.1053 | 0.1325 | 0.0950 | 0.1090 | 0.1135 |
| | upper | 0.0937 | 0.1102 | 0.1388 | 0.0994 | 0.1136 | 0.1187 |
| | MLE | 0.0904 | 0.1064 | 0.1339 | 0.0960 | 0.1100 | 0.1147 |
| $DAG_1$ (μM) | lower | 5.7050 | 1.6773 | 1.0591 | 2.3132 | - | 0.8405 |
| | upper | 7.9971 | 7.2815 | 8.6928 | 15.9434 | 9.1786 | 6.4363 |
| | MLE | 7.3927 | 4.2756 | 2.9120 | 7.4308 | 2.2524 | 2.5137 |
| $DAG_2$ (μM) | lower | 6.3008 | - | - | 9.3966 | 0.6109 | - |
| | upper | 8.8988 | 4.7821 | 6.7200 | 20.0876 | 6.0962 | 4.3649 |
| | MLE | 8.2185 | 1.1508 | 1.5041 | 10.0000 | 3.0121 | 1.0500 |
| $DAG_3$ (μM) | lower | 1.5486 | 4.5711 | 0.8496 | 0.7148 | 3.3194 | 1.5078 |
| | upper | 8.0044 | 9.1145 | 6.9445 | 5.1374 | 7.4345 | 5.4352 |
| | MLE | 4.5602 | 7.6649 | 2.3438 | 2.5926 | 5.3681 | 2.6581 |
| $DAG_4$ (μM) | lower | 4.5772 | 1.3937 | 0.8702 | 1.2306 | 2.0601 | - |
| | upper | 8.3627 | 8.1292 | 5.5281 | 6.0166 | 7.0954 | 2.6007 |
| | MLE | 6.8343 | 4.4763 | 2.0268 | 3.2367 | 4.2450 | 0.6229 |
| $DAG_5$ (μM) | lower | - | 4.9975 | 0.8045 | 4.9498 | - | 1.6384 |
| | upper | 4.2721 | 8.6025 | 5.8743 | 12.7593 | 8.9335 | 5.3954 |
| | MLE | 1.1105 | 7.9302 | 2.0741 | 7.9618 | 1.8105 | 2.7137 |

**Table S3. MLE and 95% likelihood-based confidence intervals (CI) of fit parameters from experimental data using population averaged data and measured average initial uncaged DAG for the model without $k_{out}$.** The "-" denotes that the lower or upper bound threshold was not reached and the parameter is non-identifiable within the range tested.

| Parameter | 95% CI | DMG | OSG | SOG | SLG | SAG | DArG |
|---|---|---|---|---|---|---|---|
| $k_{in}$ (1/s) | lower | 0.06928 | 0.06650 | 0.06447 | 0.10406 | 0.11524 | 0.22577 |
| | upper | 0.09641 | 0.09389 | 0.08287 | 0.14715 | 0.17376 | - |
| | MLE | 0.07530 | 0.07220 | 0.06944 | 0.11172 | 0.12815 | 0.53641 |
| $k_{met}$ (1/s) | lower | 0.00786 | 0.00881 | 0.00786 | 0.01005 | 0.01059 | 0.00774 |
| | upper | 0.00882 | 0.01082 | 0.00891 | 0.01119 | 0.01197 | 0.01620 |
| | MLE | 0.00826 | 0.00972 | 0.00830 | 0.01059 | 0.01118 | 0.01359 |
| $K_d$ (μM) | lower | 5.49079 | 2.95478 | 7.73385 | 6.76285 | 6.82204 | 7.80739 |
| | upper | 5.98339 | 3.42875 | 8.48474 | 7.30779 | 7.51164 | 10.64500 |
| | MLE | 5.72548 | 3.21228 | 8.07346 | 7.00444 | 7.11717 | 8.81467 |
| sd (μM) | lower | 0.01836 | 0.01909 | 0.02292 | 0.01526 | 0.02298 | 0.04709 |
| | upper | 0.02621 | 0.02963 | 0.03386 | 0.02719 | 0.03704 | 0.08266 |
| | MLE | 0.02050 | 0.02045 | 0.02485 | 0.01607 | 0.02451 | 0.04967 |

**Table S4. MLE and 95% likelihood-based confidence intervals (CI) of fit parameters from experimental data using population averaged data, corrected C1 concentrations, and measured average initial uncaged DAG for the model without $k_{out}$.** Available C1-EGFP-NES concentrations for DAG binding are corrected to be 23.4% and 75% of initial measured C1-EGFP-NES for SOG and SAG, respectively. The "-" denotes that the lower or upper bound threshold was not reached and the parameter is non-identifiable within the range tested.

| Parameter | 95% CI | SOG | SAG |
|---|---|---|---|
| $k_{in}$ (1/s) | lower | 0.05332 | 0.11280 |
| | upper | 0.07258 | 0.17232 |
| | MLE | 0.05859 | 0.12586 |
| $k_{met}$ (1/s) | lower | 0.01068 | 0.01095 |
| | upper | 0.01270 | 0.01243 |
| | MLE | 0.01145 | 0.01159 |
| $K_d$ (μM) | lower | 1.00616 | 4.75538 |
| | upper | 1.17834 | 5.26574 |
| | MLE | 1.08757 | 4.97424 |
| sd (μM) | lower | 0.02486 | 0.02324 |
| | upper | 0.03653 | 0.03744 |
| | MLE | 0.02702 | 0.02478 |

**Table S5. Comparison of rate parameters using single cell and population averaged data using the model without $k_{out}$.** Rate parameter and initial DAG values inferred from (1) single cell data, (2) population averaged data with measured DAG concentrations, (3) population averaged data with measured DAG concentrations and corrected C1 concentrations, and (4) values obtained from [1] using the model without $k_{out}$. Inferred rate parameters in [1] used a dataset of population averaged traces from different laser power treatments with experimentally measured uncaged DAG and corrected C1 concentrations.

| DAG | Parameter | (1) Single cell data | (2) Population ave. | (3) Corrected C1 | (4) Values in [1] |
|---|---|---|---|---|---|
| DMG | $k_{in}$ (1/s) | 0.0773 | 0.0753 | | |
| | $k_{met}$ (1/s) | 0.0057 | 0.0083 | | |
| | $K_d$ (μM) | 16.7766 | 5.7255 | | |
| | sd (μM) | 0.0904 | 0.0205 | | |
| | $DAG_{ave.}$ (μM) | 4.0420 | (1.824 ± 0.302) | | |
| OSG | $k_{in}$ (1/s) | 0.0674 | 0.0722 | | |
| | $k_{met}$ (1/s) | 0.0051 | 0.0097 | | |
| | $K_d$ (μM) | 16.6422 | 3.2123 | | |
| | sd (μM) | 0.1064 | 0.0205 | | |
| | $DAG_{ave.}$ (μM) | 3.0960 | (1.054 ± 0.175) | | |
| SOG | $k_{in}$ (1/s) | 0.0652 | 0.0694 | 0.0585902 | 0.03625 |
| | $k_{met}$ (1/s) | 0.0107 | 0.0083 | 0.0114545 | 0.07077 |
| | $K_d$ (μM) | 4.6735 | 8.0735 | 1.08757 | 0.01698 |
| | sd (μM) | 0.1339 | 0.0249 | 0.0270158 | |
| | $DAG_{ave.}$ (μM) | 1.6699 | (2.208 ± 0.367) | (2.208 ± 0.367) | |
| SLG | $k_{in}$ (1/s) | 0.0979 | 0.1117 | | |
| | $k_{met}$ (1/s) | 0.0097 | 0.0106 | | |
| | $K_d$ (μM) | 13.9923 | 7.0044 | | |
| | sd (μM) | 0.0960 | 0.0161 | | |
| | $DAG_{ave.}$ (μM) | 3.3078 | (1.850 ± 0.307) | | |
| SAG | $k_{in}$ (1/s) | 0.1146 | 0.1282 | 0.125857 | 0.09802 |
| | $k_{met}$ (1/s) | 0.0112 | 0.0112 | 0.0115862 | 0.01823 |
| | $K_d$ (μM) | 7.3750 | 7.1172 | 4.97424 | 0.866497 |
| | sd (μM) | 0.1100 | 0.0245 | 0.0247837 | |
| | $DAG_{ave.}$ (μM) | 2.5377 | (2.424 ± 0.402) | (2.424 ± 0.402) | |
| DArG | $k_{in}$ (1/s) | 0.4540 | 0.5364 | | |
| | $k_{met}$ (1/s) | 0.0286 | 0.0136 | | |
| | $K_d$ (μM) | 2.2068 | 8.8147 | | |
| | sd (μM) | 0.1147 | 0.0497 | | |
| | $DAG_{ave.}$ (μM) | 1.1270 | (2.203 ± 0.365) | | |

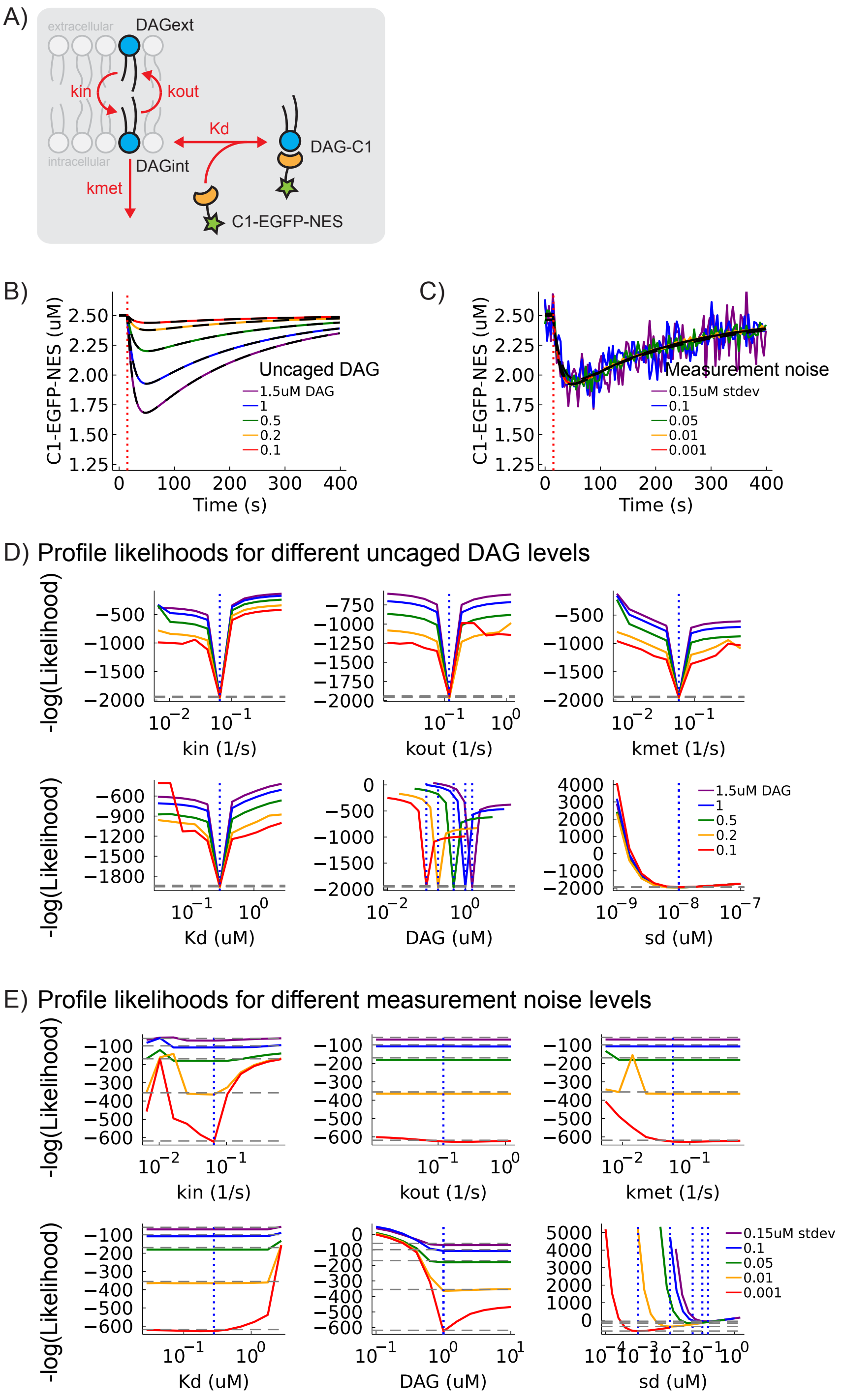


**Figure S1. Testing parameter identifiability in a model of signaling lipid dynamics with $k_{out}$ using simulated data.** (A) Schematic of a model for signaling lipid dynamics. DAG at the outer leaflet of the cell membrane (DAGext) can flip and flop between the inner and outer leaflet. In the inner leaflet, $DAG_{int}$ can either be metabolized or recruit C1-EGFP-NES proteins in the cytosol to for the membrane-associated complex DAG-C1. Rate parameters ($k_{in}$, $k_{out}$, $k_{met}$, $K_d$) of the model are shown in red. Simulated single cell traces of C1-EGFP-NES at (B) different uncaged DAG concentrations (DAG = 0.1, 0.25, 0.5, 1.0, 2.0 μM) and no measurement noise and (C) different levels of measurement noise ($\sigma$ = 0.001, 0.01, 0.05, 0.1, 0.15 μM) and fixed uncaged DAG at 1.0 μM. Fits of each individual cell are shown in black dashed lines. Red vertical dotted line indicates the time of UV exposure that results in DAG uncaging. Profile likelihoods of the model parameters from each individual cell at different uncaged DAG concentrations and levels of measurement noise are shown in (D) and (E), respectively. For all profile likelihood plots, grey horizontal dashed lines indicate the 95% likelihood-based confidence interval threshold, and blue vertical dotted lines indicate the true value of the parameter. True parameter values for the model are $k_{in} = 0.065\ s^{-1}$, $k_{out} = 0.12\ s^{-1}$, $k_{met} = 0.056\ s^{-1}$, and $K_d = 0.28$ μM. The measurement noise parameter ($\sigma$ in the equations in Section 2) is denoted as sd in all the plots.

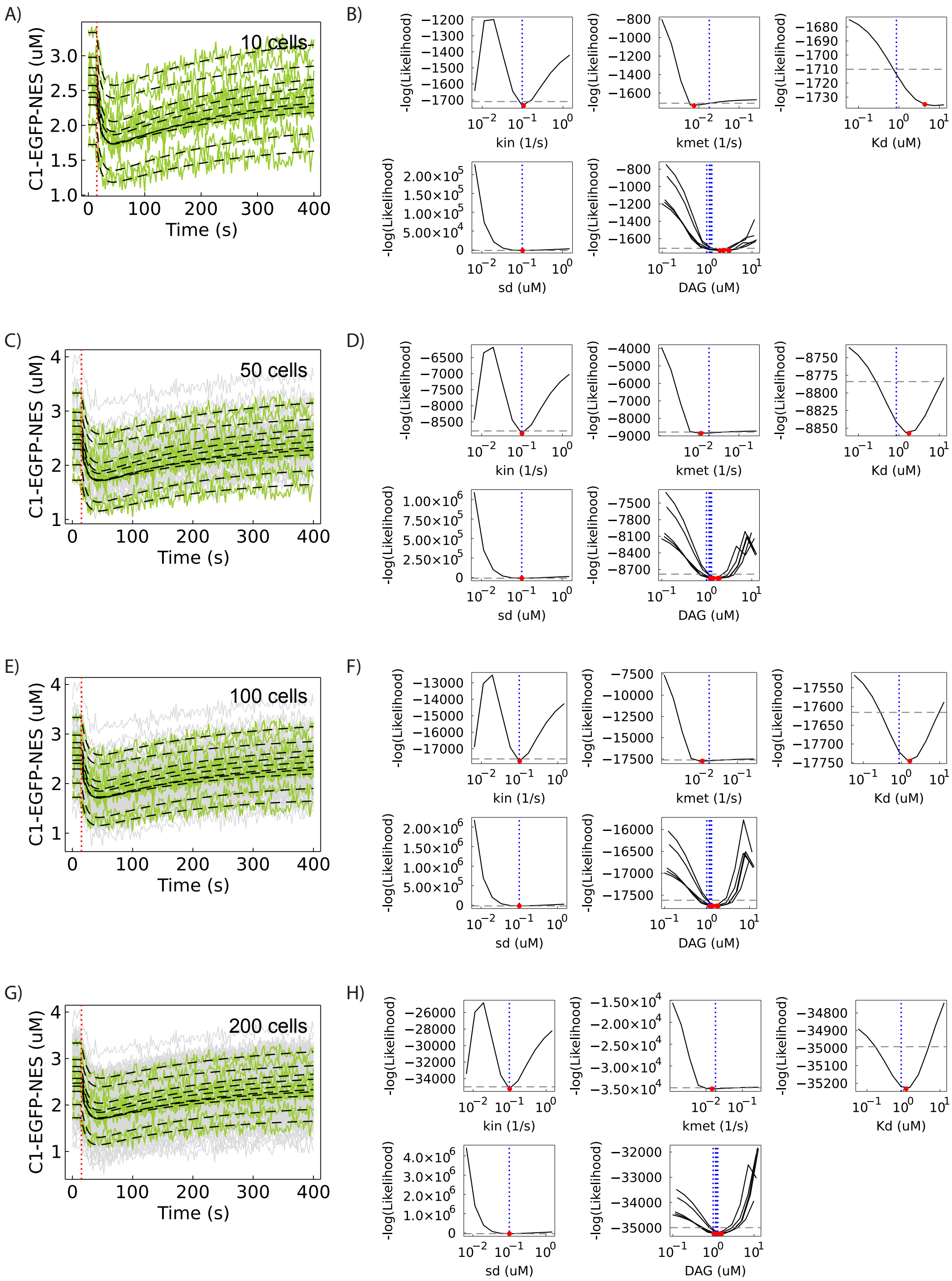


**Figure S2. Fitting and profile likelihoods using multiple cells for the model without $k_{out}$.** Single cell traces of C1-EGFP-NES dynamics (grey lines) after uncaging of DAG (left hand plots). Ten representative cell traces (green lines) and their respective fits (dashed black lines) are shown. Red vertical dotted line indicates the time of UV exposure that results in DAG uncaging. Profile likelihoods of the model parameters (right hand plots). Grey horizontal dashed lines indicate the 95% likelihood-based confidence interval threshold and red circles indicate the fit MLE of the parameter. Profile likelihoods of uncaged DAG are shown for the same five representative cells in the population. Fitting and profile likelihood analysis was performed on different cell population sizes at (A, B) 10, (C, D) 50, (E, F) 100, and (G, H) 200 cells. Data was simulated with random initial C1 and uncaged DAG concentrations from gaussian distributions N($\mu$ = 2.5 μM, $\sigma$ = 0.5 μM) and N(1.2 μM, 0.1 μM), respectively. True parameter values for the model are $k_{in}$ = 0.098 $s^{-1}$, $k_{met}$ = 0.01823 $s^{-1}$, and $K_d$ = 0.866 μM. Gaussian measurement noise N(0 μM, 0.1 μM) was added to the simulated data.

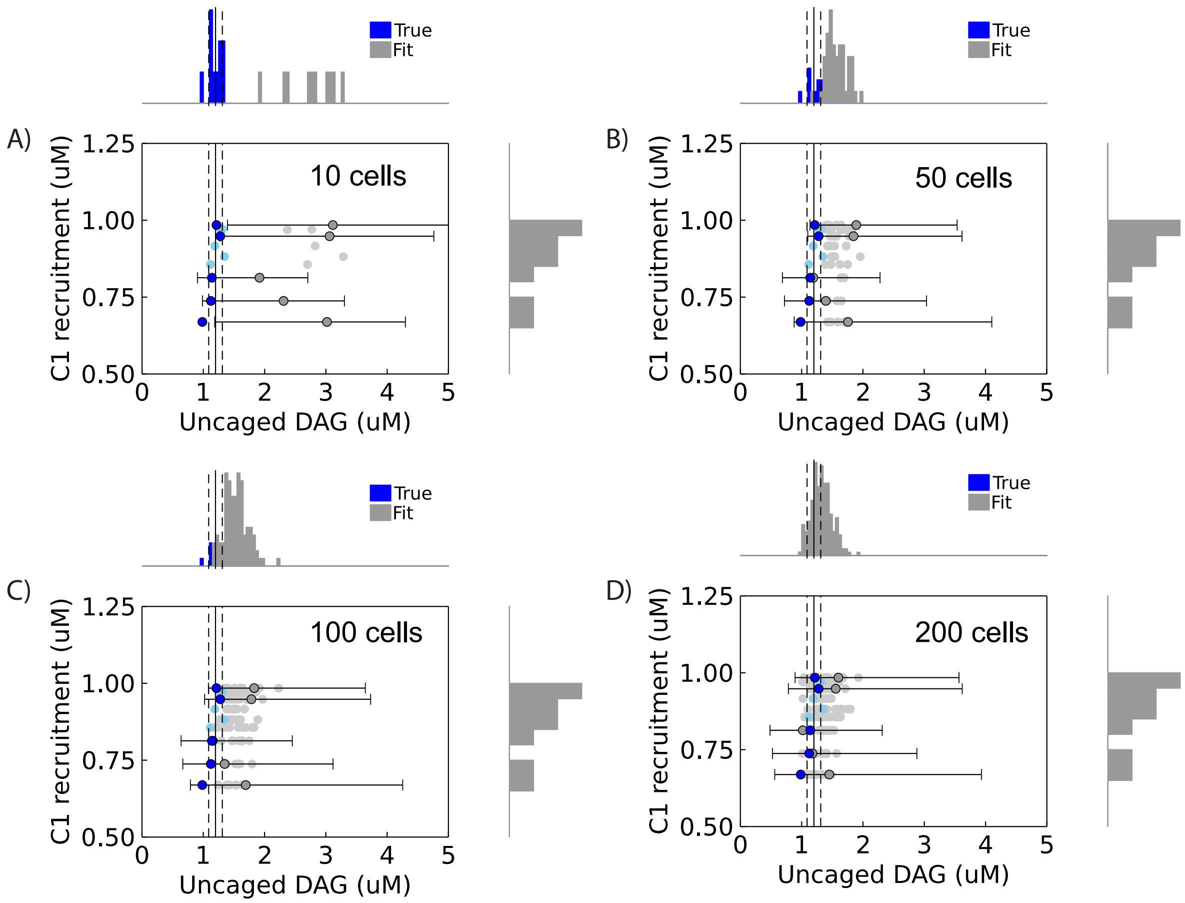


**Figure S3. C1 recruitment vs. inferred uncaged DAG concentrations.** Correlation between inferred uncaged DAG and measured C1 recruitment in the cell population using (A) 10, (B) 50, (C) 100, and (D) 200 cells as in Fig. S2. Each light blue dot refers to a single cell. Dark blue dots with error bars show five representative cells and their 95% likelihood-based confidence intervals. Vertical solid and dashed lines indicate the known mean and standard deviation of uncaged DAG. Marginal histograms of uncaged DAG and C1 recruitment are shown on each axis.

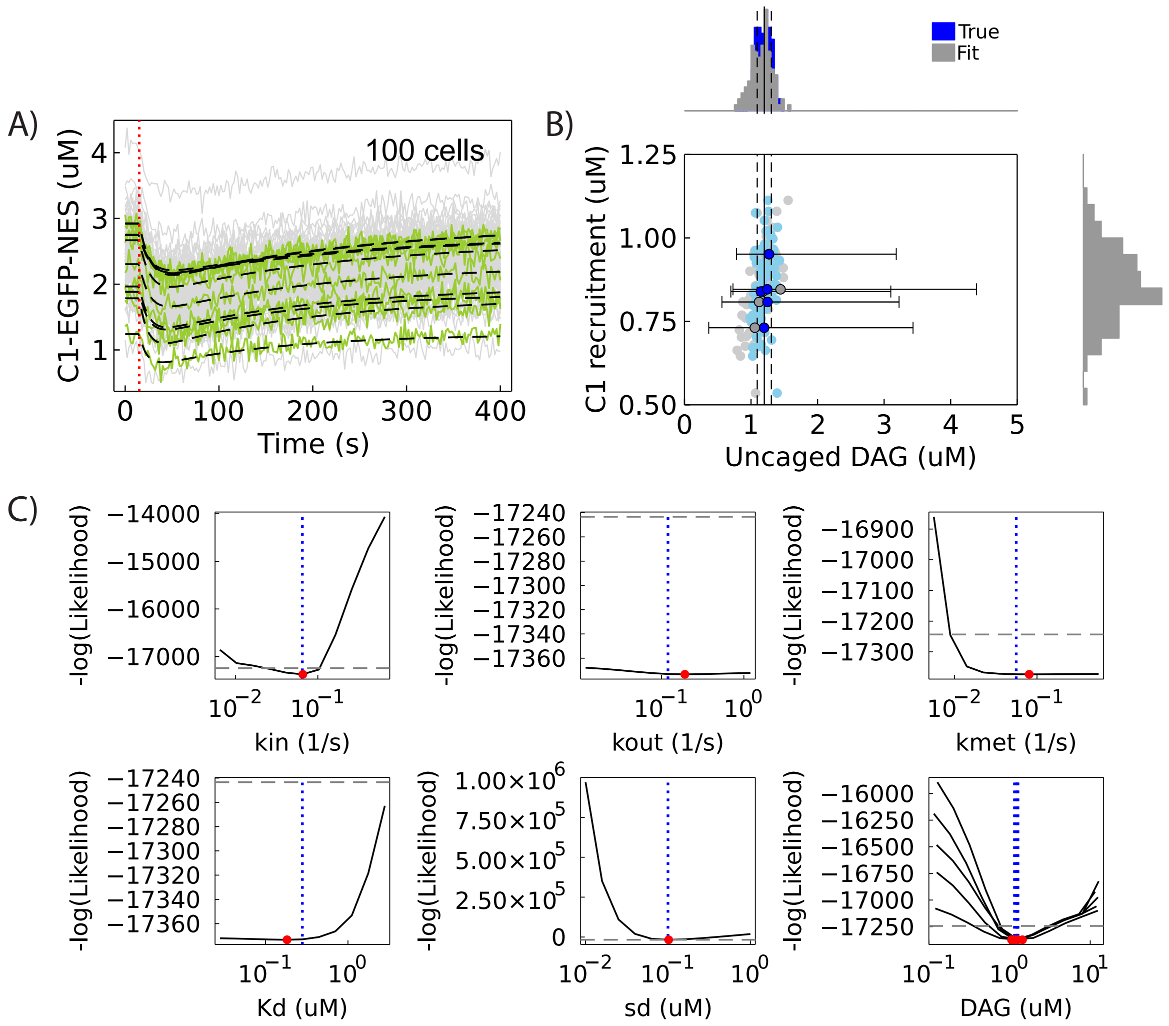


**Figure S4. Fitting and profile likelihoods on DAG signalling model with $k_{out}$ using 100 cells.** (A) Single cell traces of C1-EGFP-NES dynamics (grey lines) after uncaging of DAG. Ten representative cell traces (green lines) and their respective fits (dashed black lines) are shown. Red vertical dotted line indicates the time of UV exposure that results in DAG uncaging. (B) Correlation between inferred uncaged DAG and measured C1 recruitment in the cell population. Each light blue dot refers to a single cell. Dark blue dots with error bars show five representative cells and their 95% likelihood-based confidence intervals. Vertical solid and dashed lines indicate the known mean and standard deviation of uncaged DAG. Marginal histograms of uncaged DAG and C1 recruitment are shown on each axis. (C) Profile likelihoods of the model parameters. Grey horizontal dashed lines indicate the 95% likelihood-based confidence interval threshold and red circles indicate the fit MLE of the parameter. Profile likelihoods of uncaged DAG are shown for the same five representative cells in the population. Data was simulated with random initial C1 and uncaged DAG concentrations from gaussian distributions N($\mu$ = 2.5 $\mu$M, $\sigma$ = 0.5 $\mu$M) and N(1.2 $\mu$M, 0.1 $\mu$M), respectively. True parameter values for the model are $k_{in}$ = 0.065 $s^{-1}$, $k_{out}$ =0.12 $s^{-1}$, $k_{met}$ = 0.056 $s^{-1}$, and $K_d$ = 0.28 $\mu$M. Gaussian measurement noise N(0 $\mu$M, 0.1 $\mu$M) was added to the simulated data.

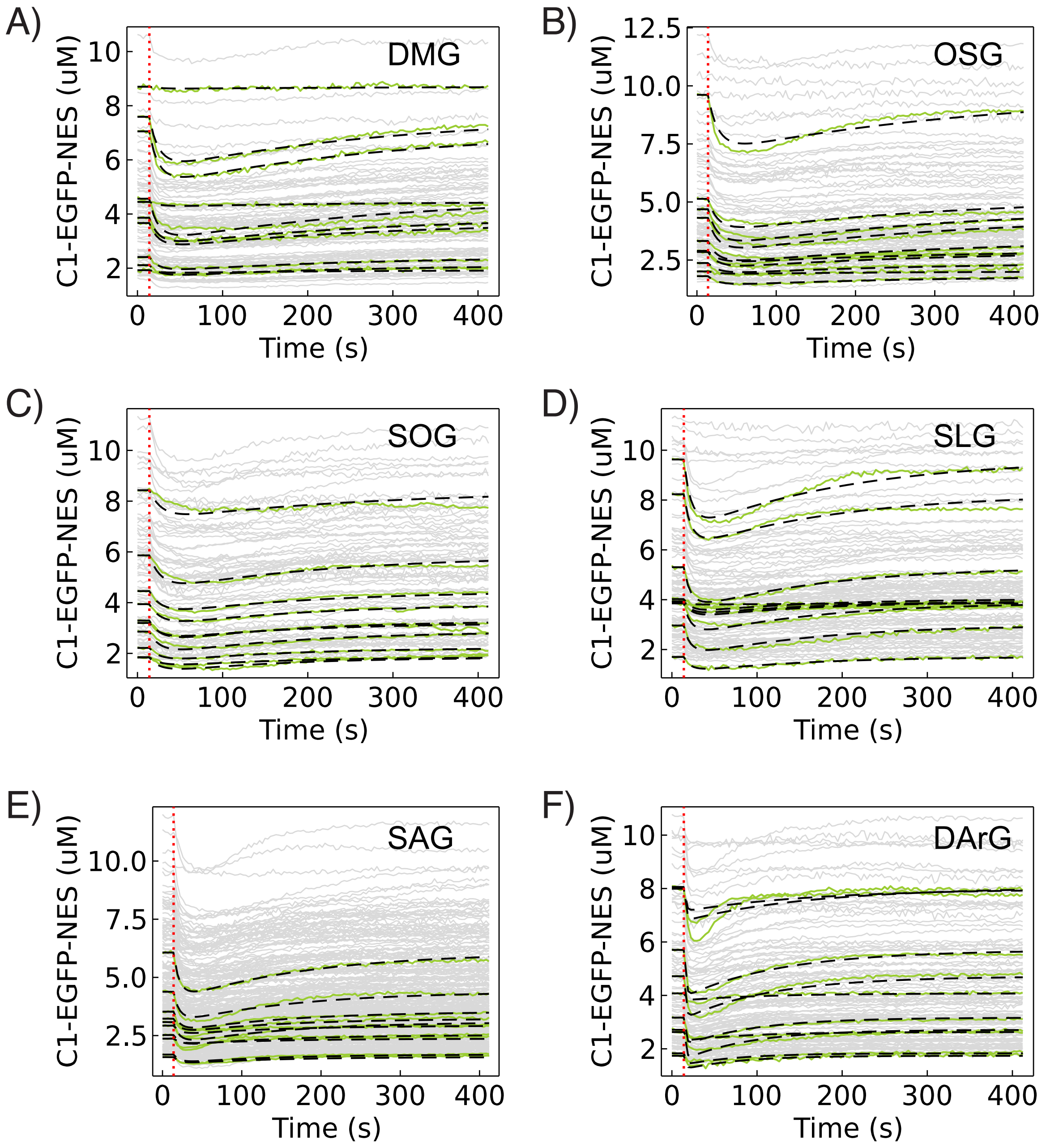


**Figure S5. Fitting of different experimental DAG traces after uncaging.** Single cell traces of C1-EGFP-NES dynamics (grey lines) after uncaging of caged (A-F) DMG, OSG, SOG, SLG, SAG, and DArG lipids. Ten representative cell traces (green lines) and their respective fits (dashed black lines) are shown. Red vertical dotted line indicates the time of UV exposure that results in DAG uncaging. Total number of cell traces used are 97, 105, 86, 108, 273, and 120 for DMG, OSG, SOG, SLG, SAG, and DArG, respectively.

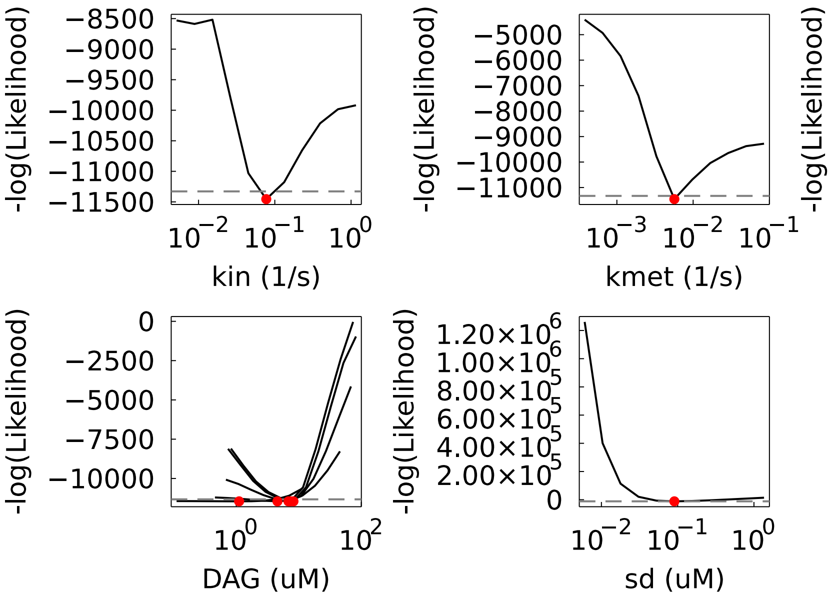


**Figure S6. Profile likelihoods of model parameters for DMG.** Grey horizontal dashed lines indicate the 95% likelihood-based confidence interval threshold and red circles indicate the fit MLE of the parameter. Profile likelihoods of uncaged DAG are shown for the same five representative cells in the population. The total number of cells used for the analysis is 97.

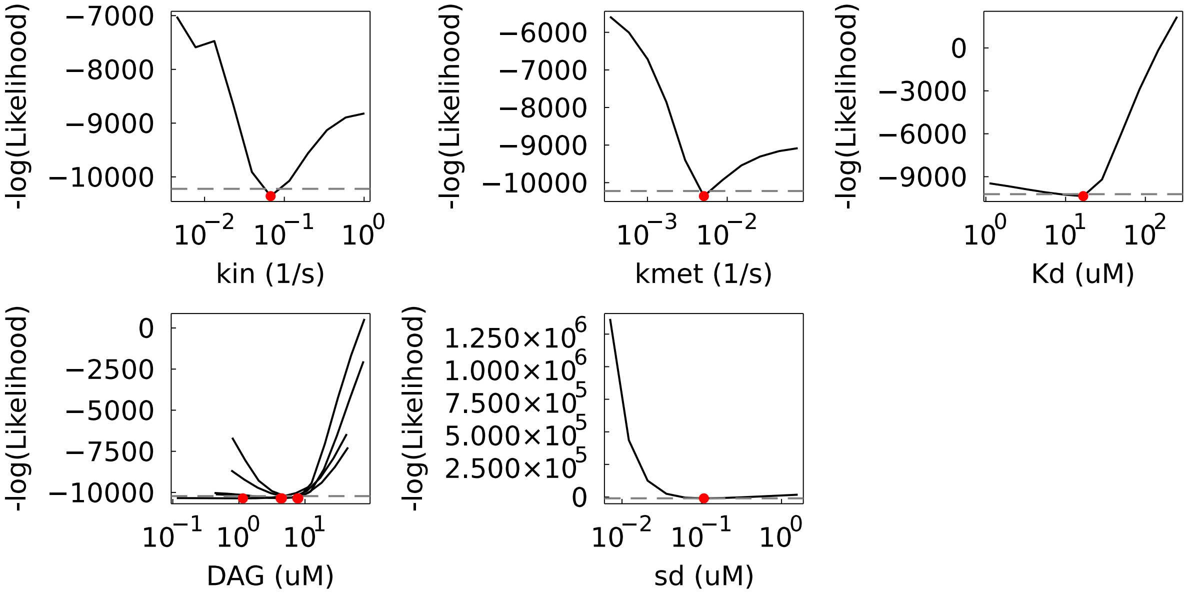


**Figure S7. Profile likelihoods of model parameters for OSG.** Grey horizontal dashed lines indicate the 95% likelihood-based confidence interval threshold and red circles indicate the fit MLE of the parameter. Profile likelihoods of uncaged DAG are shown for the same five representative cells in the population. The total number of cells used for the analysis is 105.

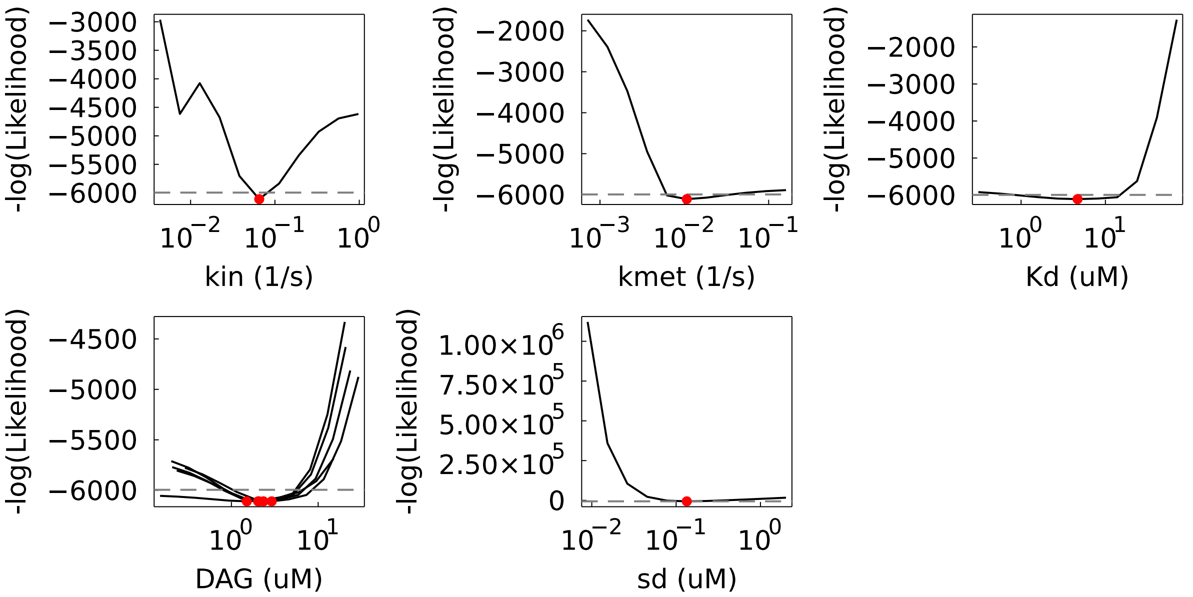


**Figure S8. Profile likelihoods of model parameters for SOG.** Grey horizontal dashed lines indicate the 95% likelihood-based confidence interval threshold and red circles indicate the fit MLE of the parameter. Profile likelihoods of uncaged DAG are shown for the same five representative cells in the population. The total number of cells used for the analysis is 86.

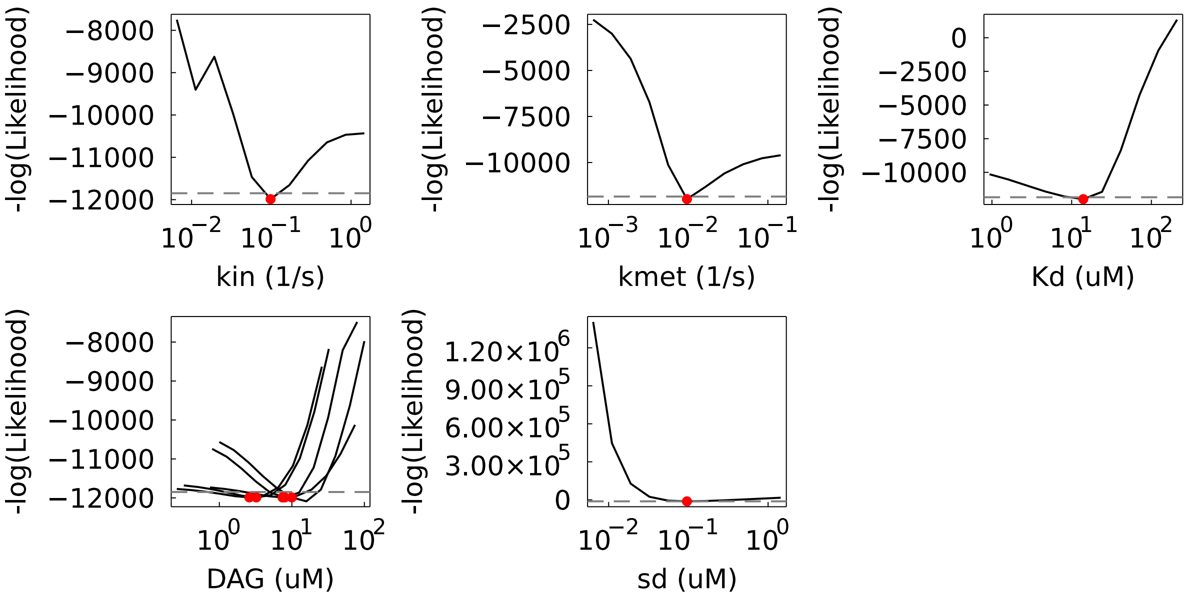


**Figure S9. Profile likelihoods of model parameters for SLG.** Grey horizontal dashed lines indicate the 95% likelihood-based confidence interval threshold and red circles indicate the fit MLE of the parameter. Profile likelihoods of uncaged DAG are shown for the same five representative cells in the population. The total number of cells used for the analysis is 108.

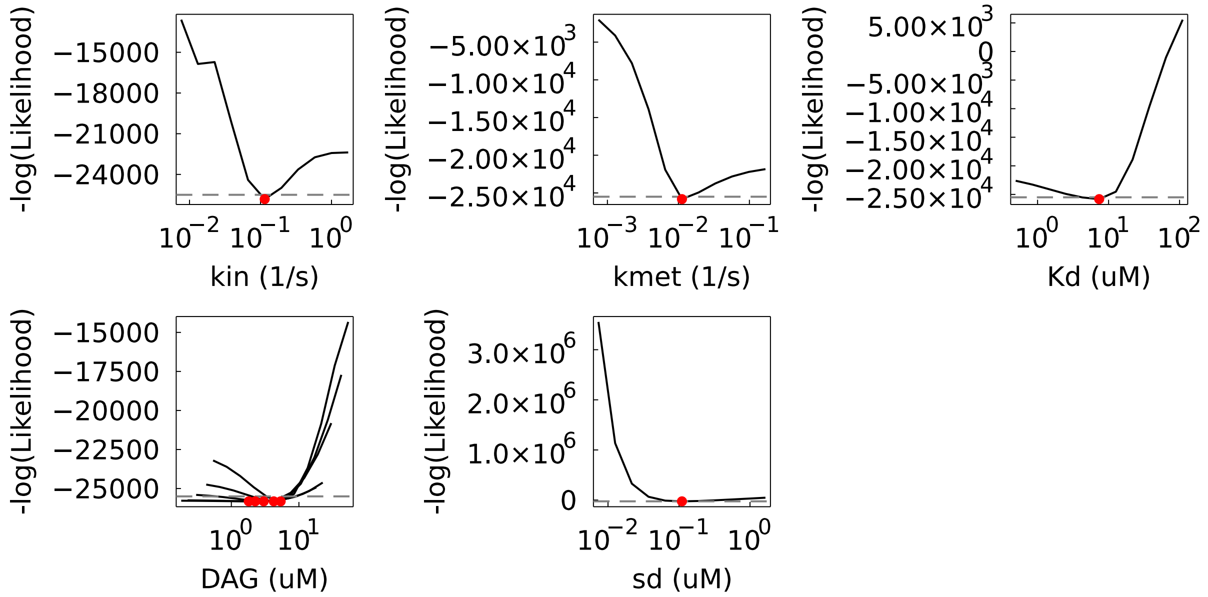


**Figure S10. Profile likelihoods of model parameters for SAG.** Grey horizontal dashed lines indicate the 95% likelihood-based confidence interval threshold and red circles indicate the fit MLE of the parameter. Profile likelihoods of uncaged DAG are shown for the same five representative cells in the population. The total number of cells used for the analysis is 273.

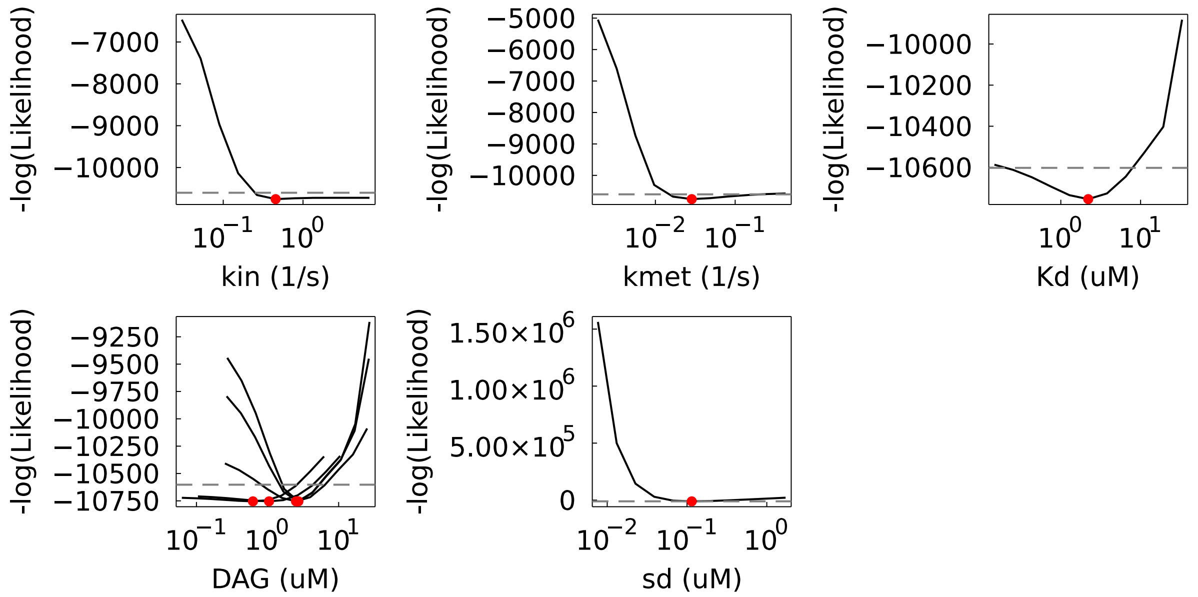


**Figure S11. Profile likelihoods of model parameters for DArG,** Grey horizontal dashed lines indicate the 95% likelihood-based confidence interval threshold and red circles indicate the fit MLE of the parameter. Profile likelihoods of uncaged DAG are shown for the same five representative cells in the population. The total number of cells used for the analysis is 120.

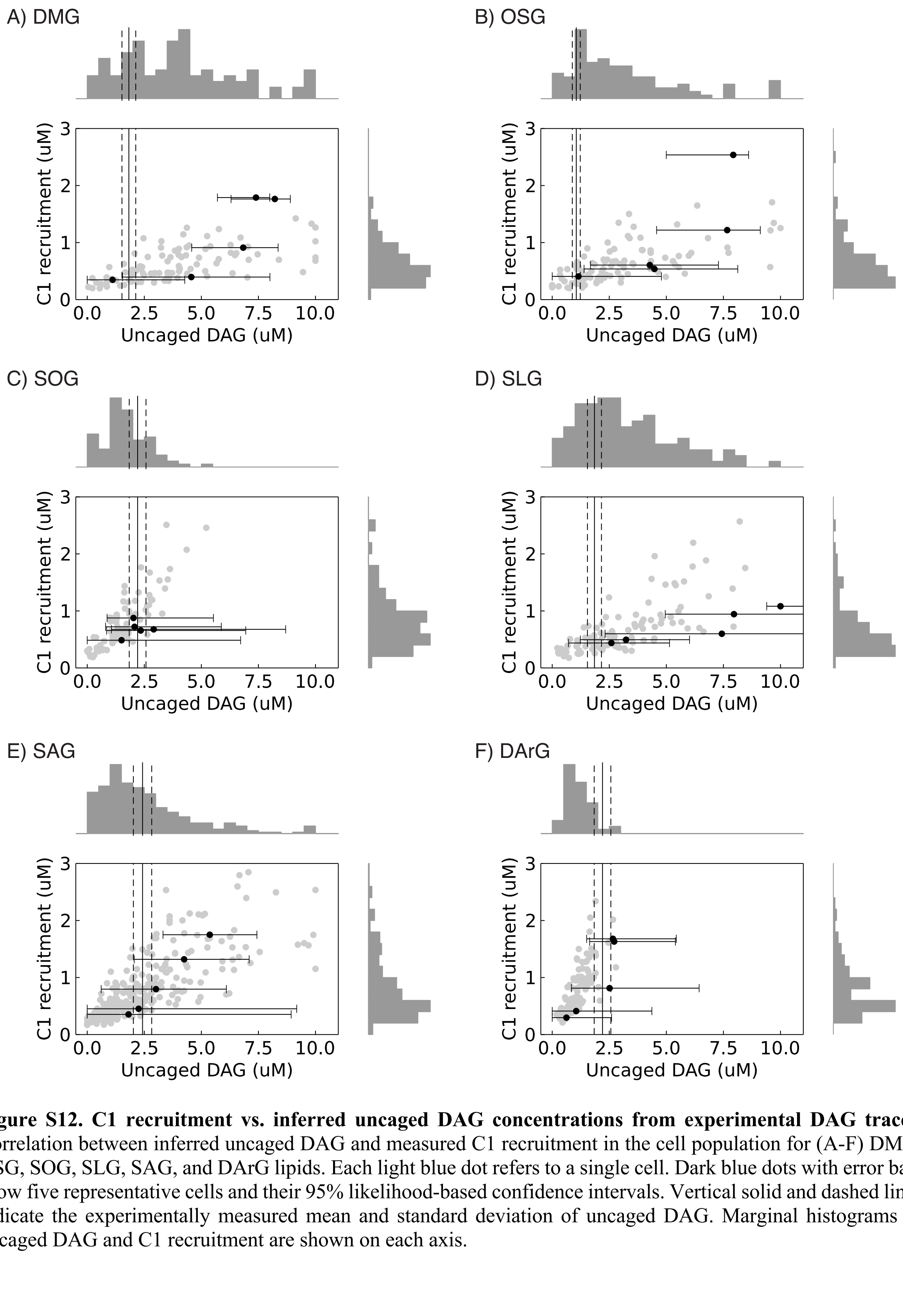


**Figure S12. C1 recruitment vs. inferred uncaged DAG concentrations from experimental DAG traces.** Correlation between inferred uncaged DAG and measured C1 recruitment in the cell population for (A-F) DMG, OSG, SOG, SLG, SAG, and DArG lipids. Each light blue dot refers to a single cell. Dark blue dots with error bars show five representative cells and their 95% likelihood-based confidence intervals. Vertical solid and dashed lines indicate the experimentally measured mean and standard deviation of uncaged DAG. Marginal histograms of uncaged DAG and C1 recruitment are shown on each axis.

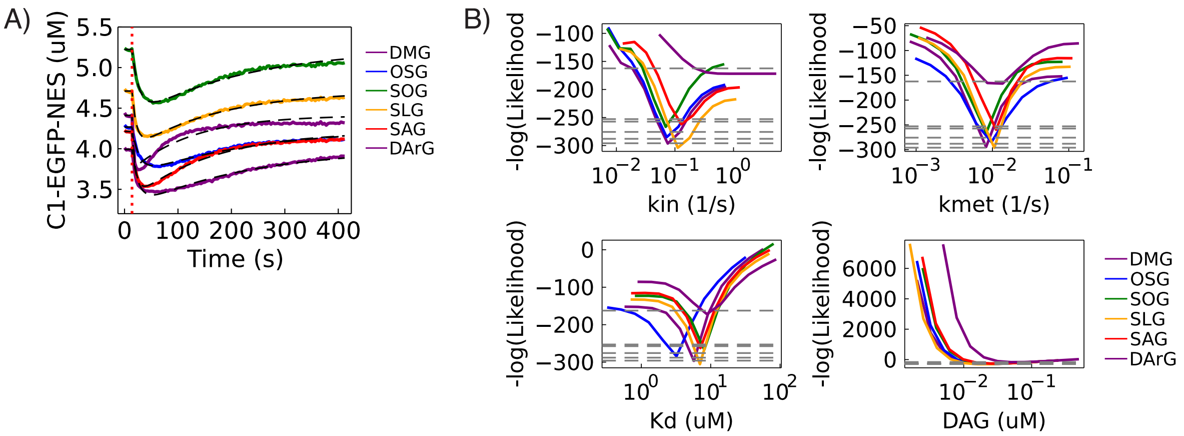


**Figure S13. Fitting and profile likelihoods using mean traces only.** (A) Mean traces of C1-EGFP-NES dynamics after uncaging of the different DAG lipids (colored lines). Black dotted lines are the fits of each DAG lipid using the model without $k_{out}$. Red vertical dotted line indicates the time of UV exposure that results in DAG uncaging. (B) Profile likelihoods of the model parameters using mean traces of the different DAG lipids. Grey horizontal dashed lines indicate the 95% likelihood-based confidence interval threshold.

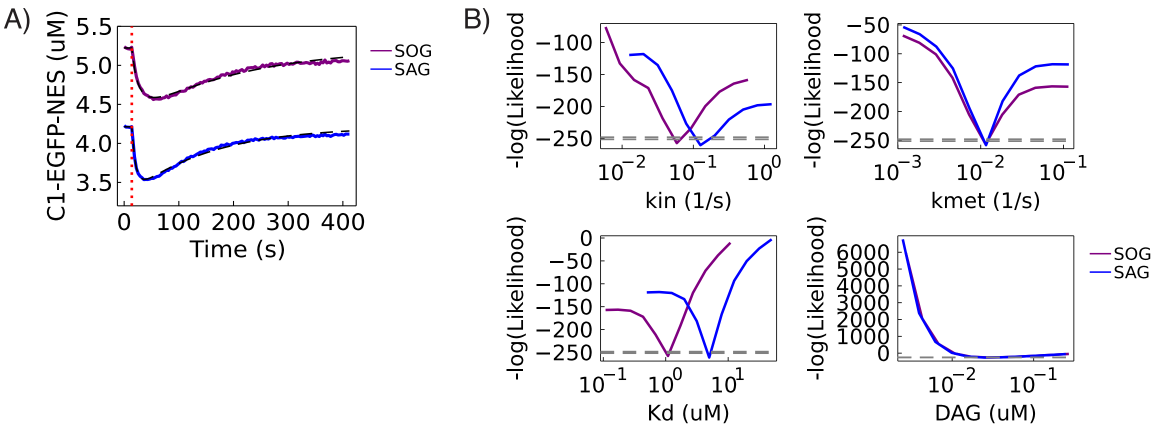


**Figure S14. Fitting and profile likelihoods using mean traces and corrected C1 concentrations.** (A) Mean traces of C1-EGFP-NES dynamics after uncaging of the SOG and SAG (colored lines). Available C1-EGFP-NES concentrations for DAG binding are corrected to be 23.4% and 75% of initial measured C1-EGFP-NES for SOG and SAG. Black dotted lines are the fits of each DAG lipid using the model without $k_{out}$. Red vertical dotted line indicates the time of UV exposure that results in DAG uncaging. (B) Profile likelihoods of the model parameters using mean traces of the different DAG lipids. Grey horizontal dashed lines indicate the 95% likelihood-based confidence interval threshold.

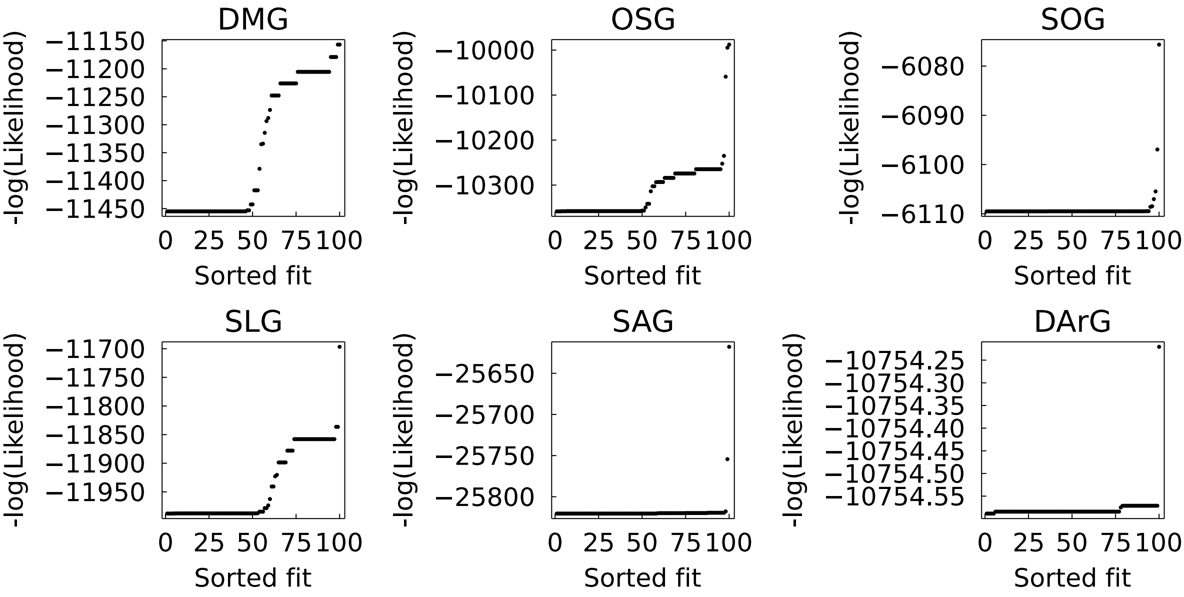


**Figure S15. Minimum negative loglikelihoods using random initial conditions for single cell DAG traces.** MLE results from 100 random initial conditions within the ranges 0-1 $s^{-1}$, 0-1 $s^{-1}$, 0-20 μM, and 0-0.5 μM for $k_{in}$, $k_{met}$, $K_d$, and σ, respectively.

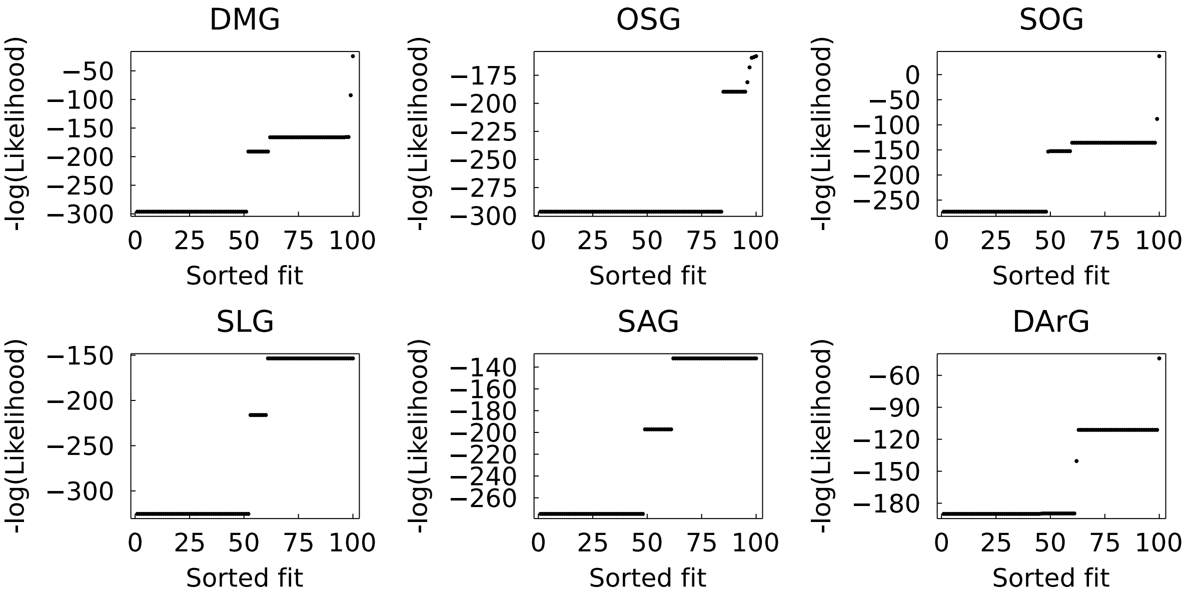


**Figure S16. Minimum negative loglikelihoods using random initial conditions for population mean traces.** MLE results from 100 random initial conditions within the ranges 0-1 $s^{-1}$, 0-1 $s^{-1}$, 0-20 µM, and 0-0.5 µM for $k_{in}$, $k_{met}$, $K_d$, and σ, respectively.

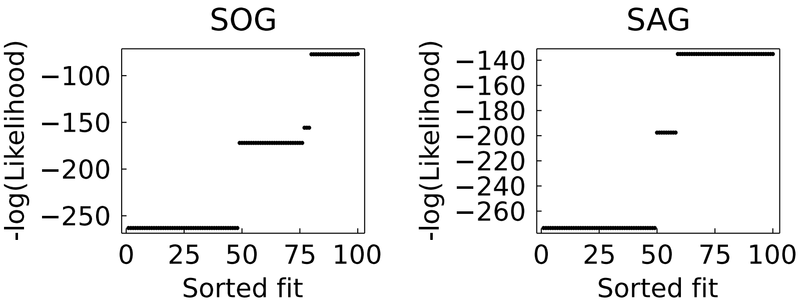


**Figure S17. Minimum negative loglikelihoods using random initial conditions for population mean traces with corrected C1 concentrations for SOG and SAG.** MLE results from 100 random initial conditions within the ranges 0-1 $s^{-1}$, 0-1 $s^{-1}$, 0-20 μM, and 0-0.5 μM for $k_{in}$, $k_{met}$, $K_d$, and σ, respectively.

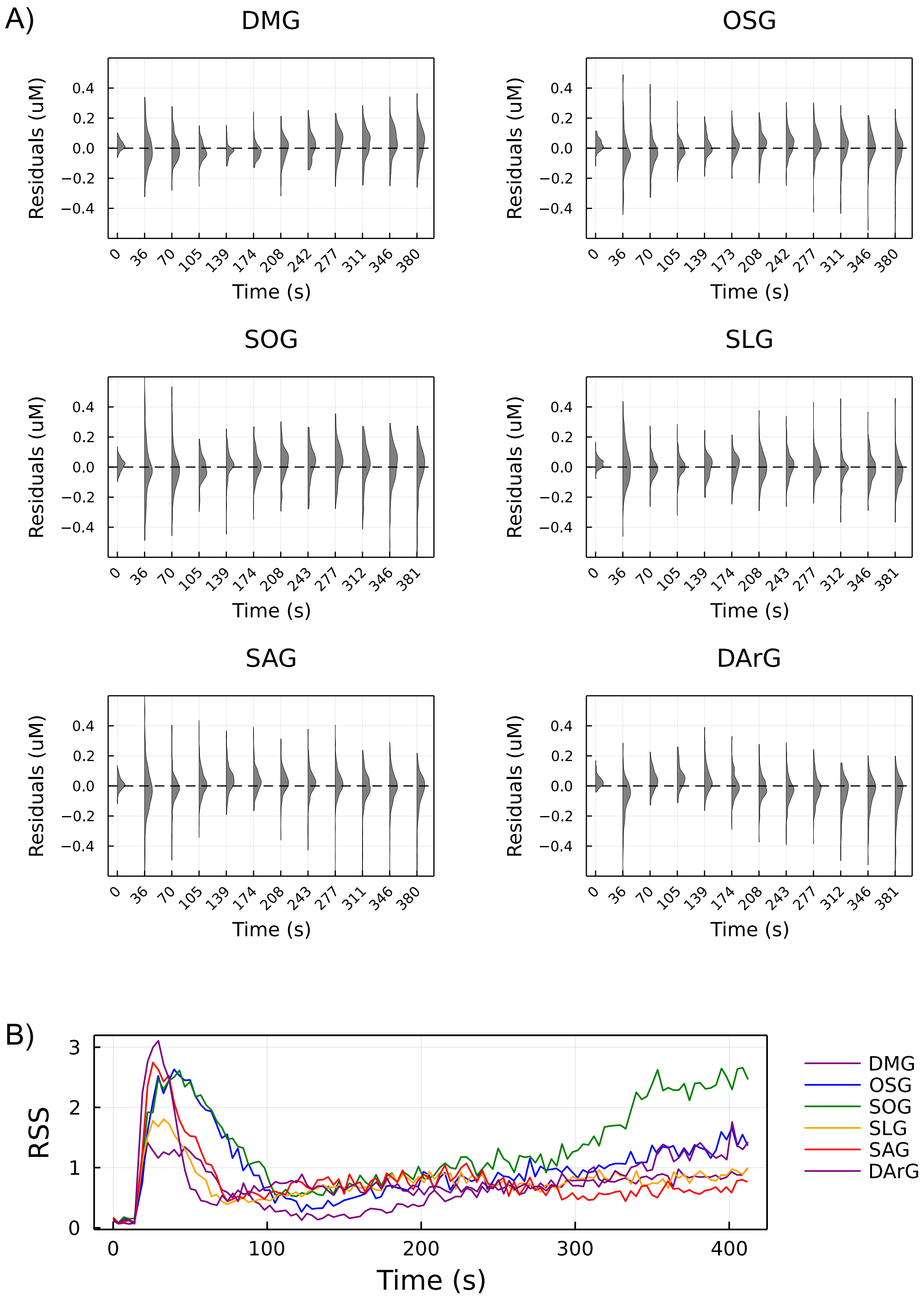


**Figure S18. Residual analysis of fitting on experimental data using the model without $k_{out}$.** (A) Histogram of residuals between single cell data and fit of each DAG over time. (B) Residual square of sum (RSS) of each DAG over time from single cell data and fit.

A)

| cgDAG | fatty acid composition |
|---|---|
| cgDMG | SN1: 14:0; SN2: 14:0 |
| cgOSG | SN1: 18:1; SN2: 18:0 |
| cgSOG | SN1: 18:0; SN2: 18:1 |
| cgSLG | SN1: 18:0; SN2: 18:2 |
| cgSAG | SN1: 18:0; SN2: 20:4 |
| cgDArG | SN1: 20:4; SN2: 20:4 |
| cg1,3DOG | SN1: 18:1; SN3: 18:1 |

B)

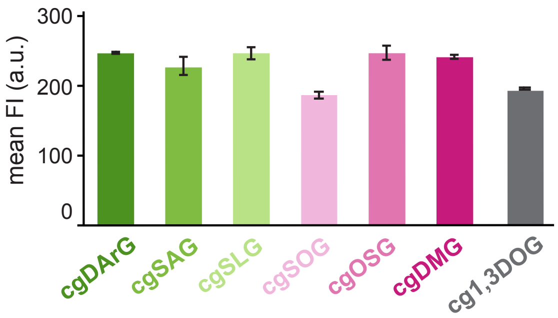


**Figure S19. Caged DAG composition and loading on cell membrane.** (A) Chain length and saturation degree of the different caged DAGs used in this study. (B) Uniform fluorescence signal of the coumarin cage in different DAGs after loading on the cell membrane.

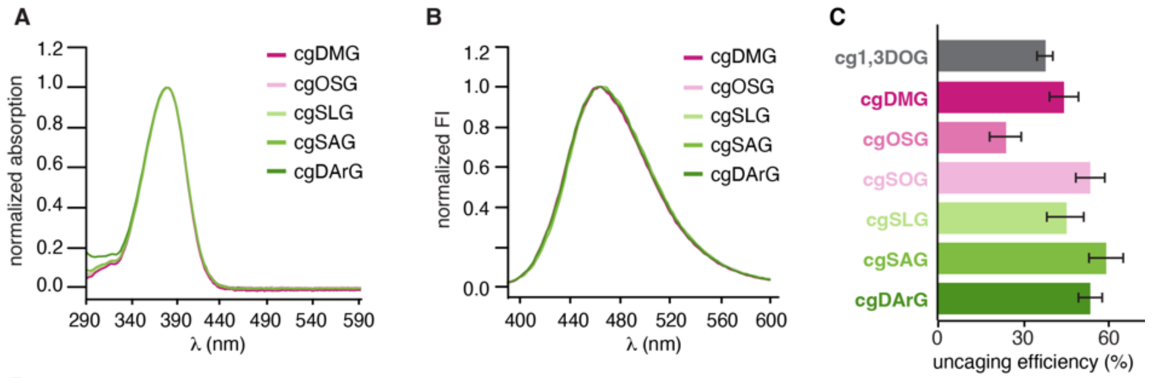


**D**

| cgDAG | FI[before] | FI[after] | E | E(corr) | FI(base) |
|---|---|---|---|---|---|
| cgSOG | 386.22 ± 5.68 | 336.35 ± 4.47 | 13 ± 0.31% | 54 ± 5% | 0.76 ± 0.03 |
| cgSAG | 551.28 ± 9.31 | 437.79 ± 6.57 | 0.21 ± 0.50% | 59 ± 6% | 0.65 ± 0.03 |
| cg1,3DOG | 358.47 ± 4.84 | 306.87 ± 3.86 | 14 ± 0.27% | 38 ± 3% | 0.62 ± 0.03 |
| cgOSG | 303.85 ± 3.16 | 278.14 ± 2.59 | 8 ± 0.18% | 24 ± 5% | 0.65 ± 0.04 |
| cgSLG | 392.04 ± 4.62 | 341.01 ± 3.82 | 13 ± 0.20% | 45 ± 6% | 0.71 ± 0.02 |
| cgDMG | 450.92 ± 6.00 | 377.07 ± 4.72 | 16 ± 0.29% | 44 ± 5% | 0.63 ± 0.02 |
| cgDArG | 507.50 ± 7.65 | 380.01 ± 5.00 | 25 ± 0.52% | 53 ± 4% | 0.53 ± 0.02 |

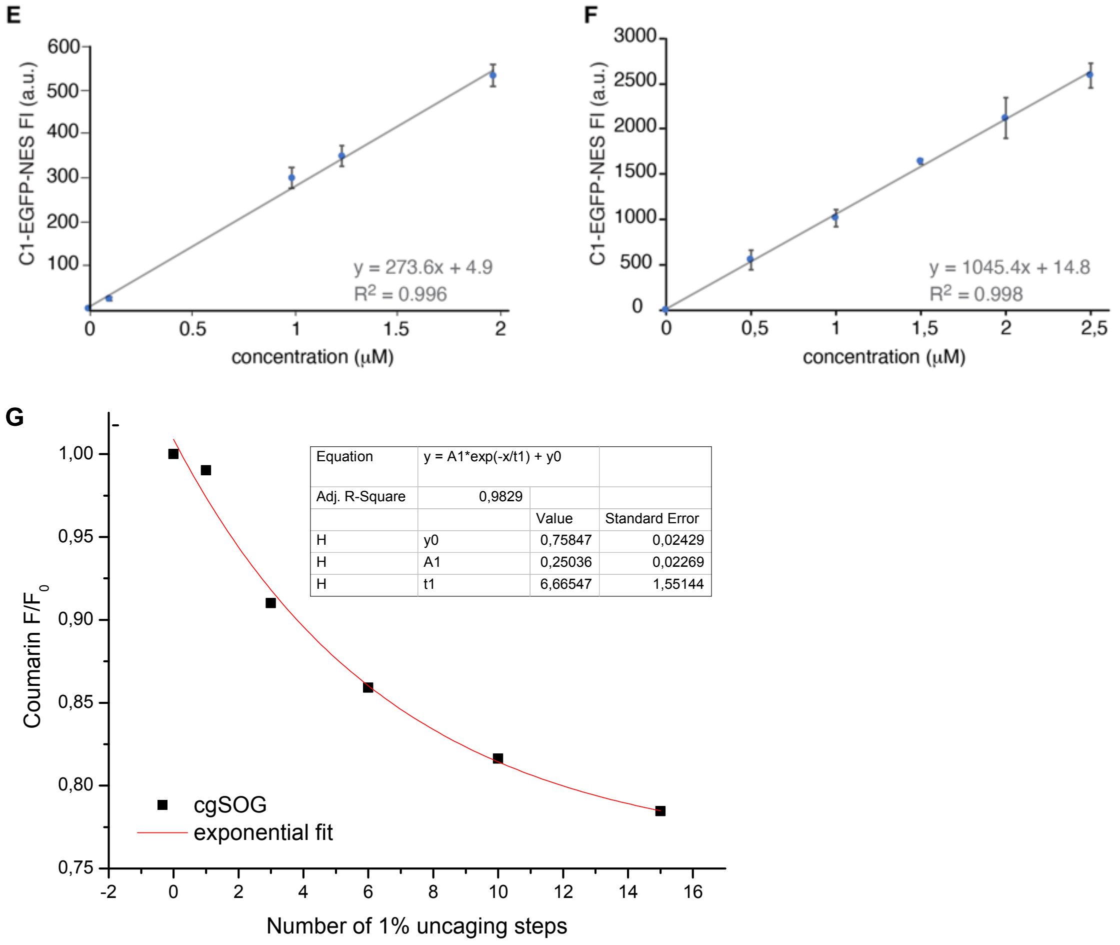


**Figure S20. Characterization of DAGs and C1-EGFP-NES.** (A) Absorption spectra of the different caged DAGs. (B) Fluorescence spectra of the different caged DAGs at 380 nm excitation. (C) Uncaging efficiencies of the different caged DAGs after uncaging with a 405 nm laser at 10% intensity. (D) Calculation of uncaging efficiencies accounting for background fluorescence. (E-F) Calibration curves of C1-EGFP-NES in the same imaging settings as our live cell experiments to obtain single cell traces of C1-EGFP-NES and cell surface and volume determination, respectively. (G) Monoexponential decay fit of multiple 1% uncaging steps to determine the asymptotic value ($FI_{base}$) indicative of the background fluorescence.

**Synthesis of DMG and DAG**

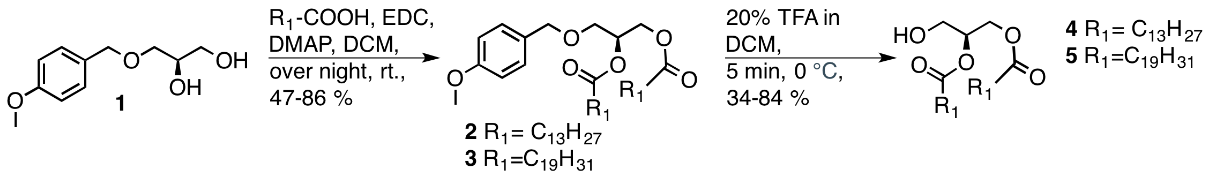


**Synthesis of OSG**

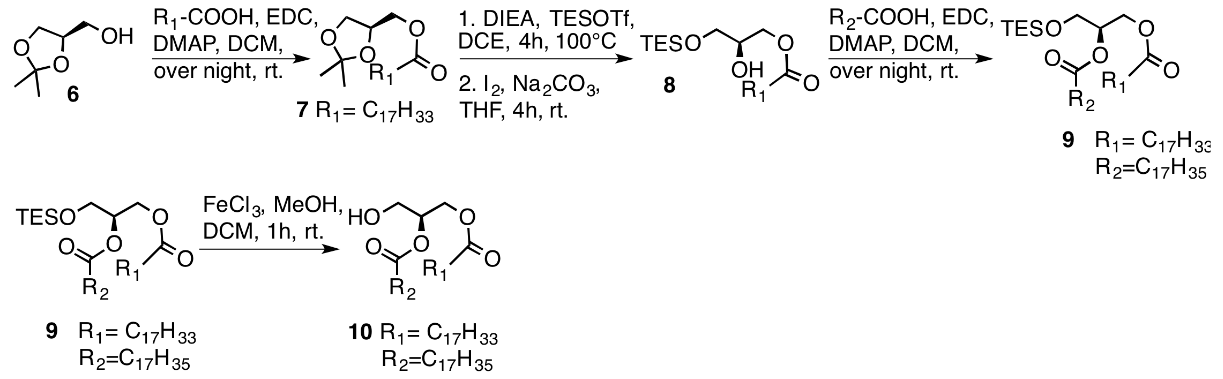


**Synthesis of caged natural DAGs**

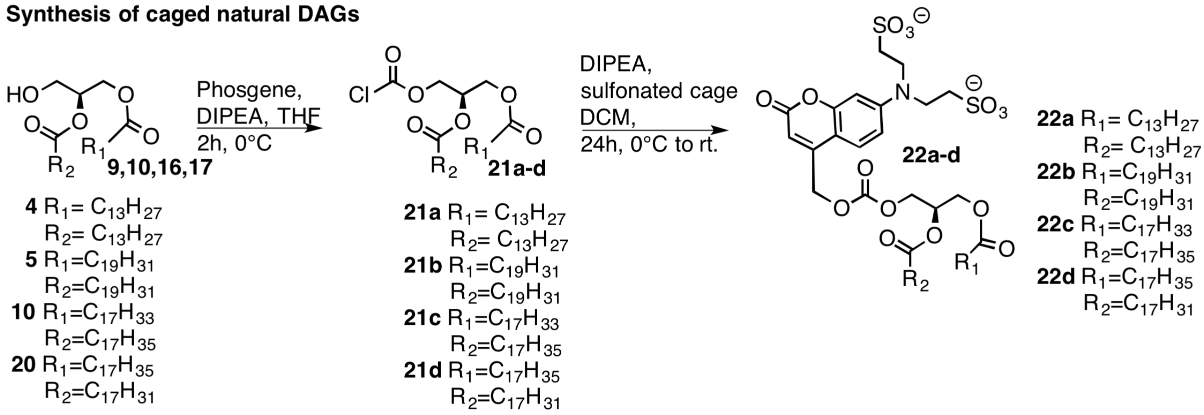


**Figure S21. Overview of synthesis of DAGs and caged DAGs.**

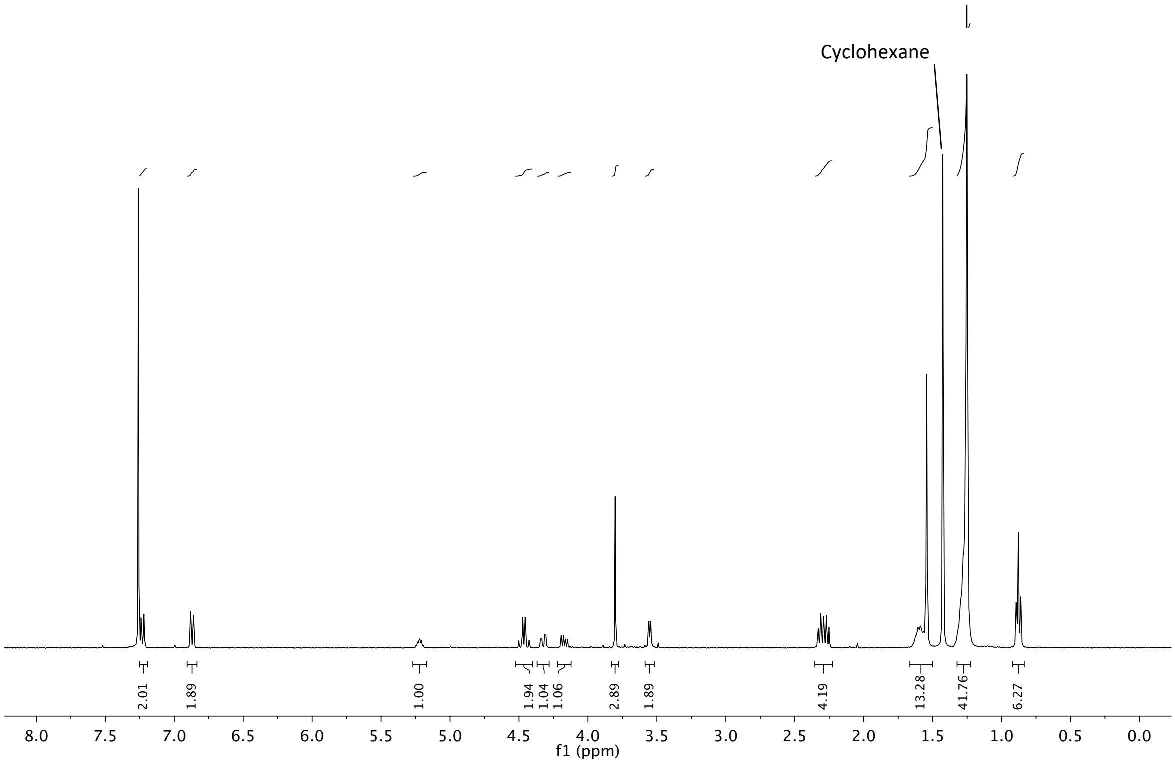


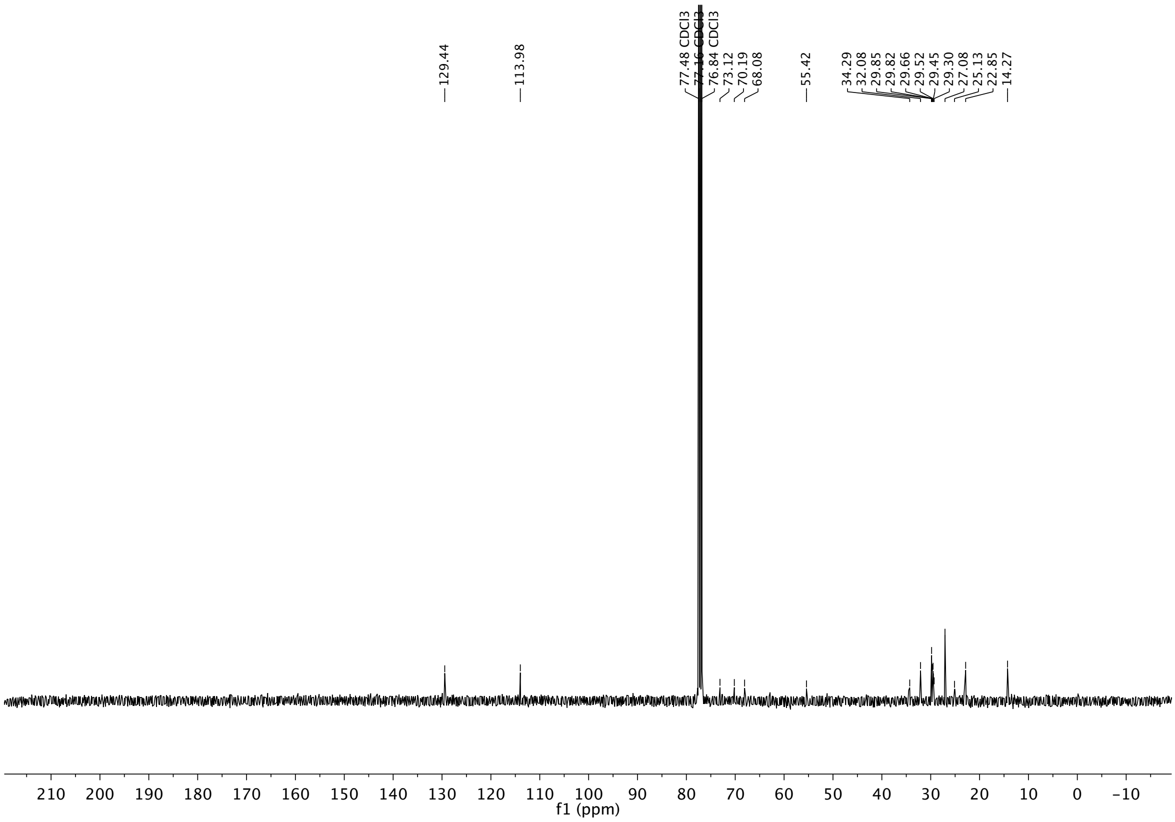


**Figure S22. NMR spectra of (2)**

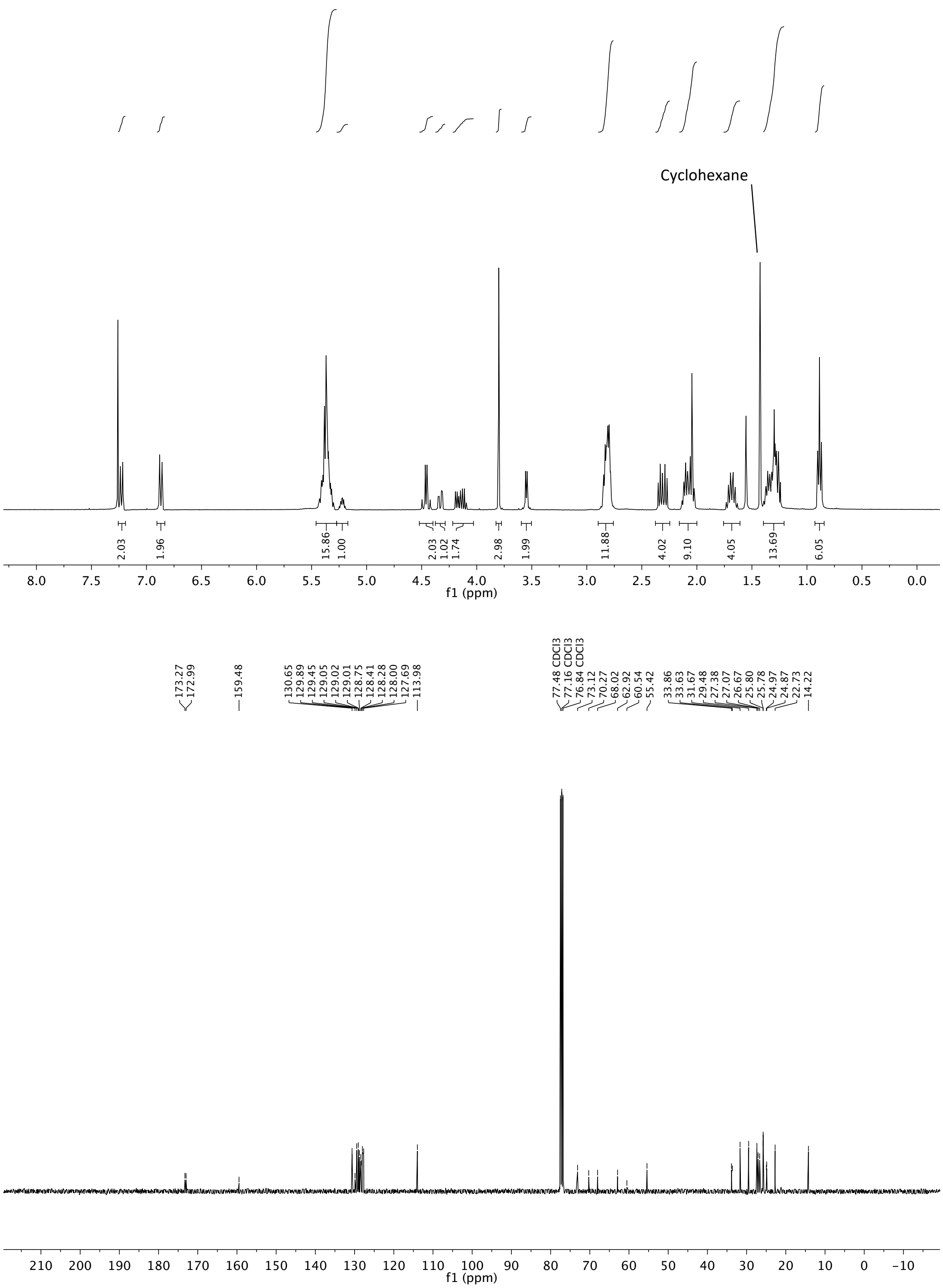


**Figure S23. NMR spectra of (3)**

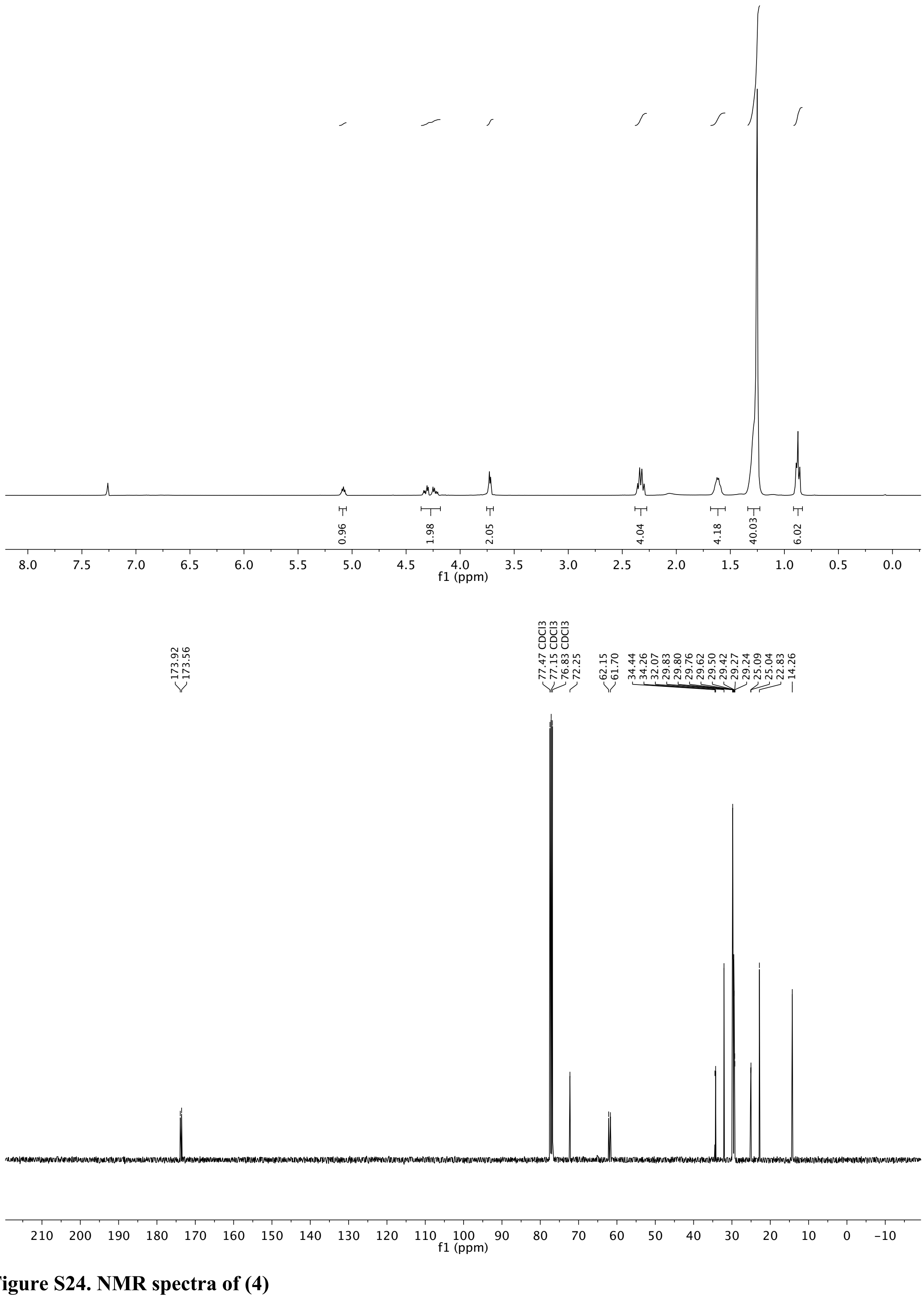


**Figure S24. NMR spectra of (4)**

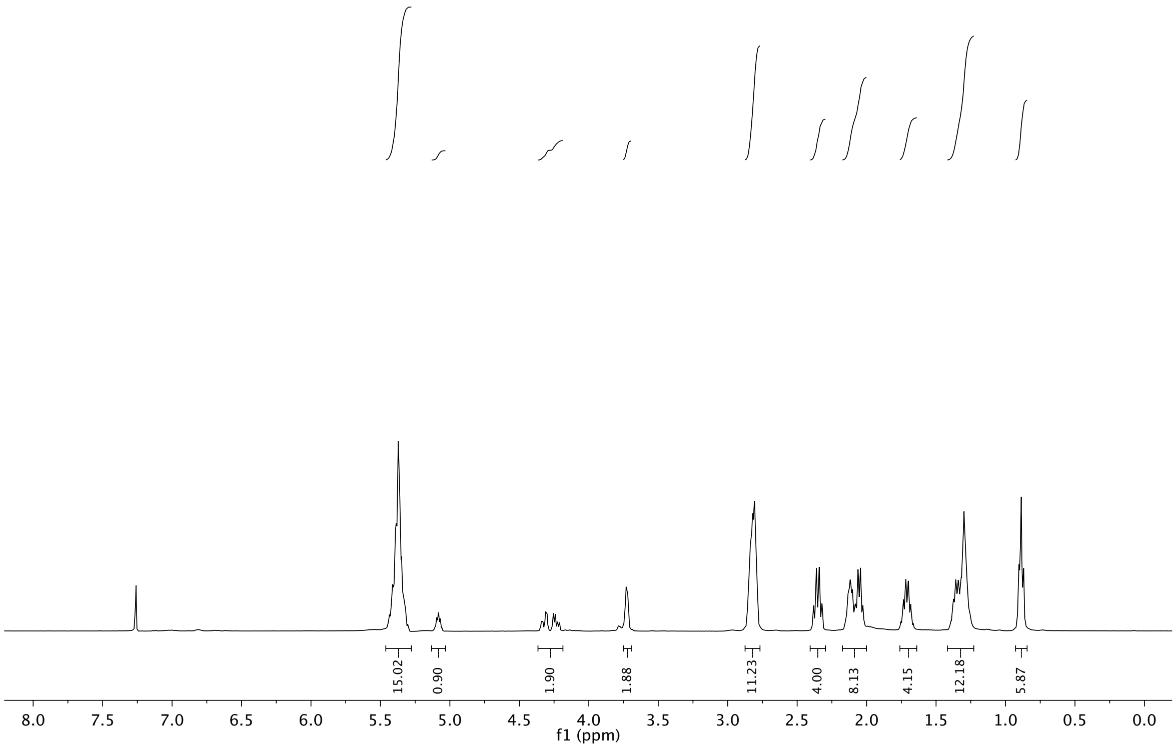


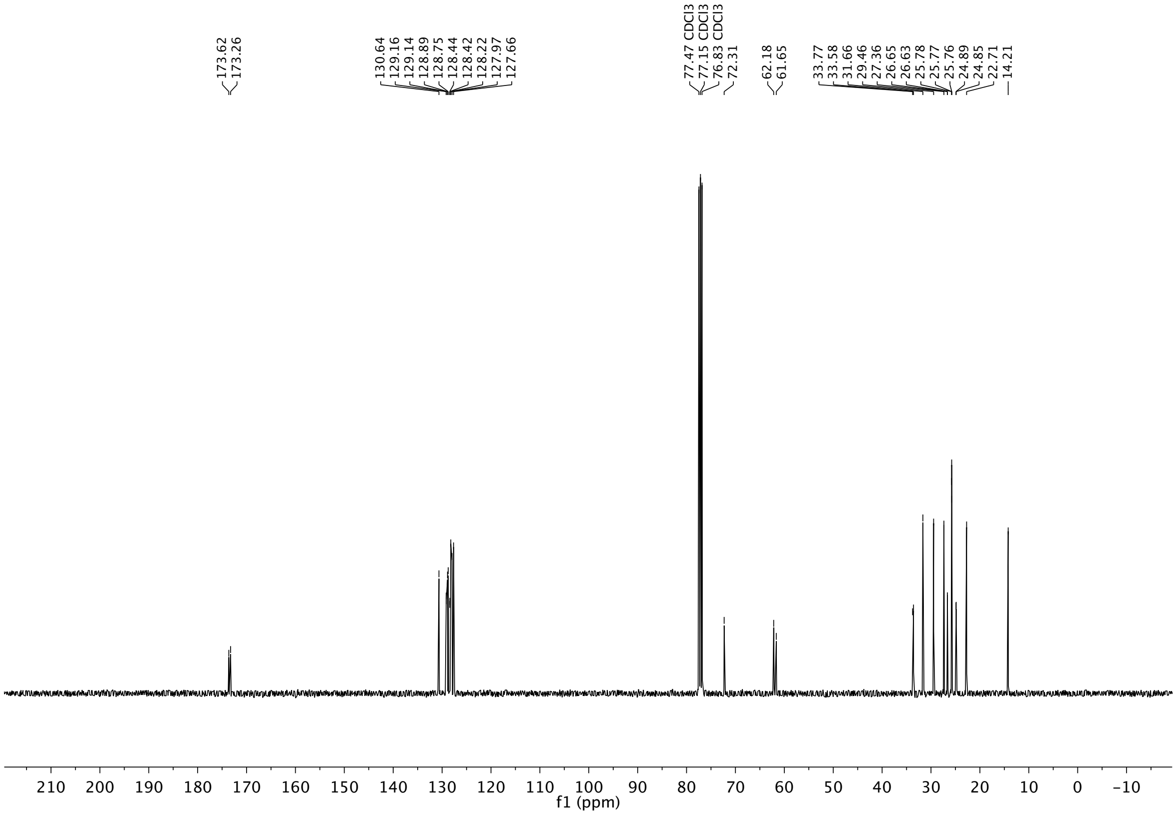


**Figure S25. NMR spectra of (5)**

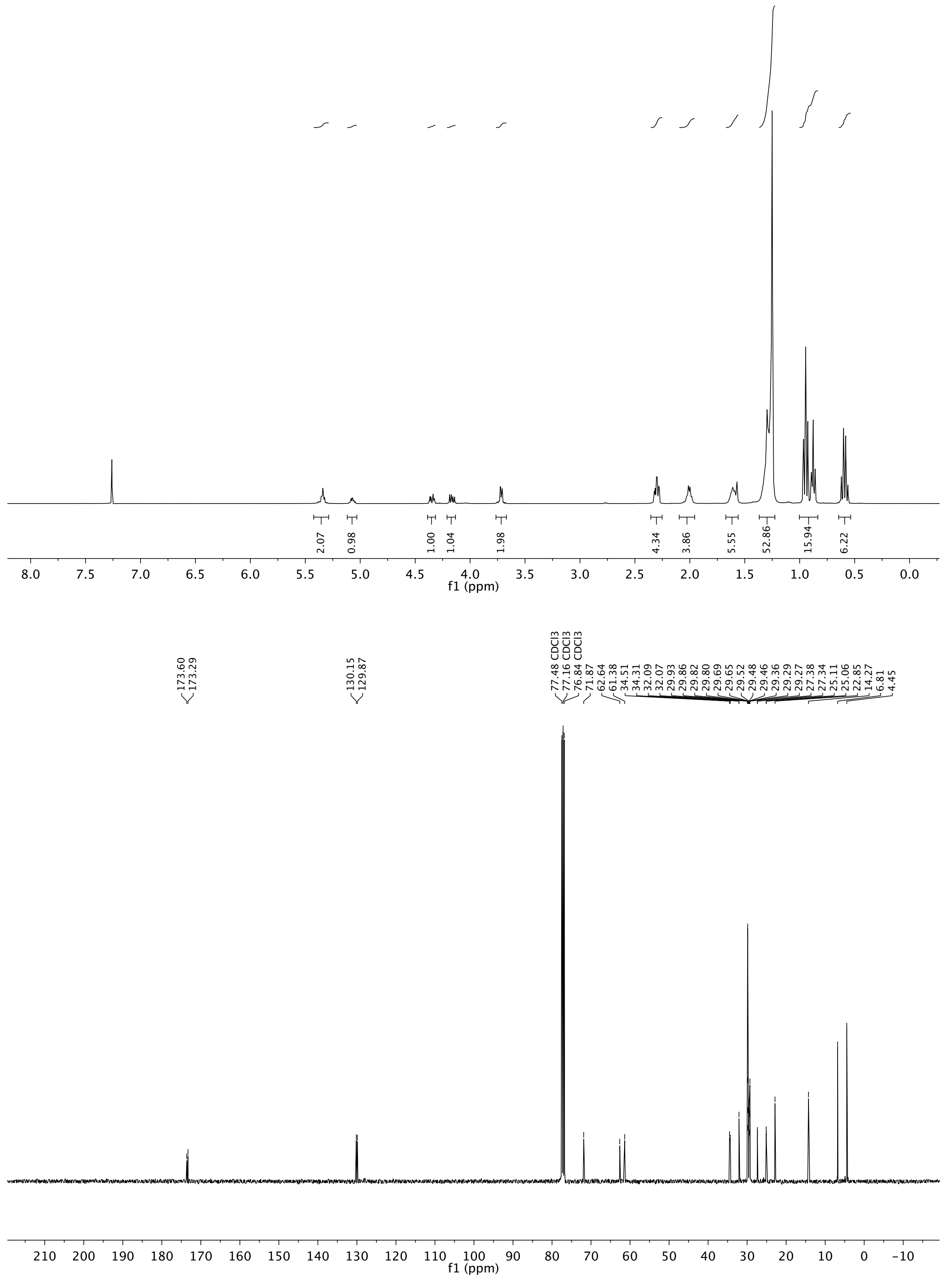


**Figure S26. NMR spectra of (9)**

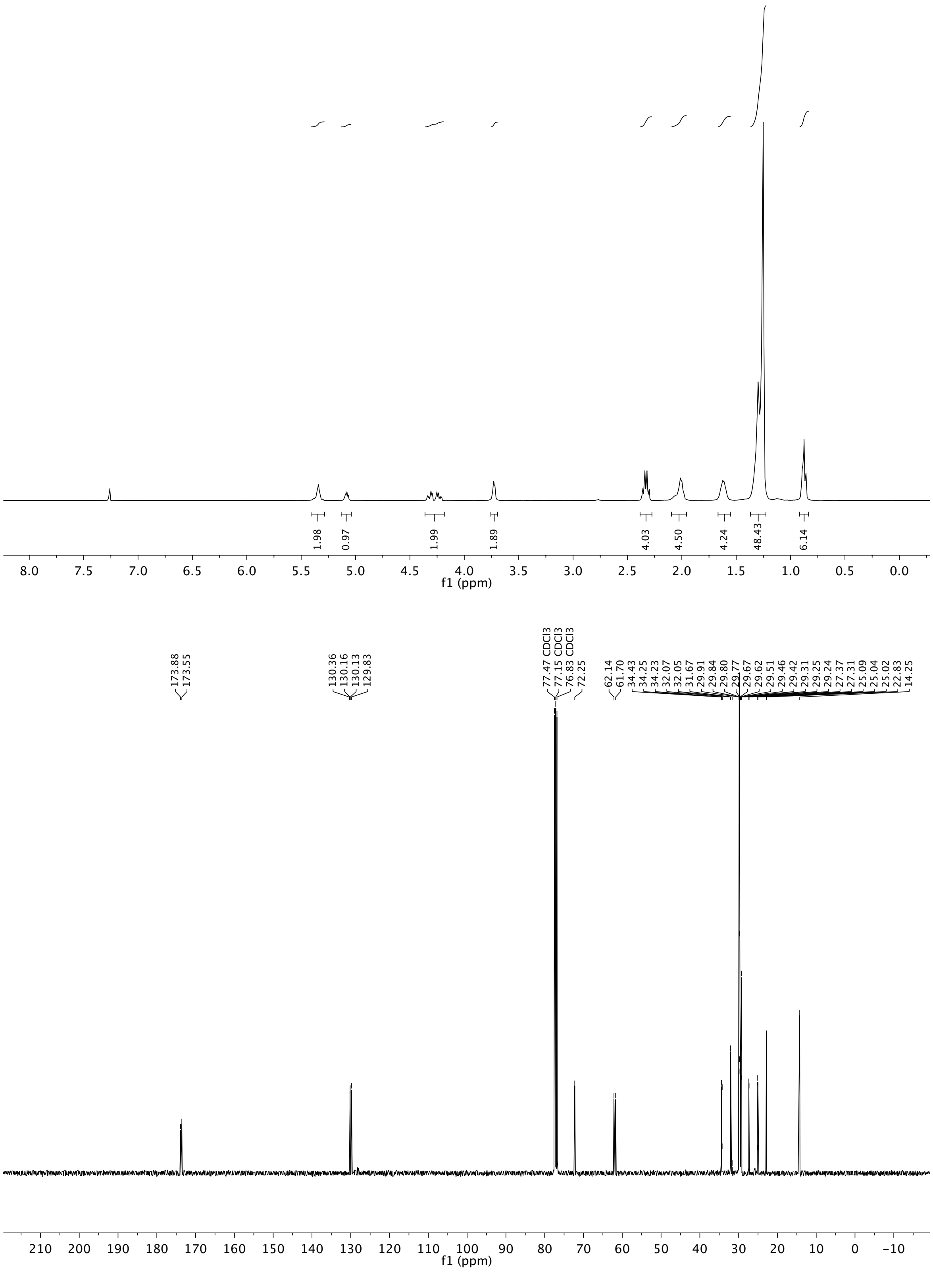


**Figure S27. NMR spectra of (10)**

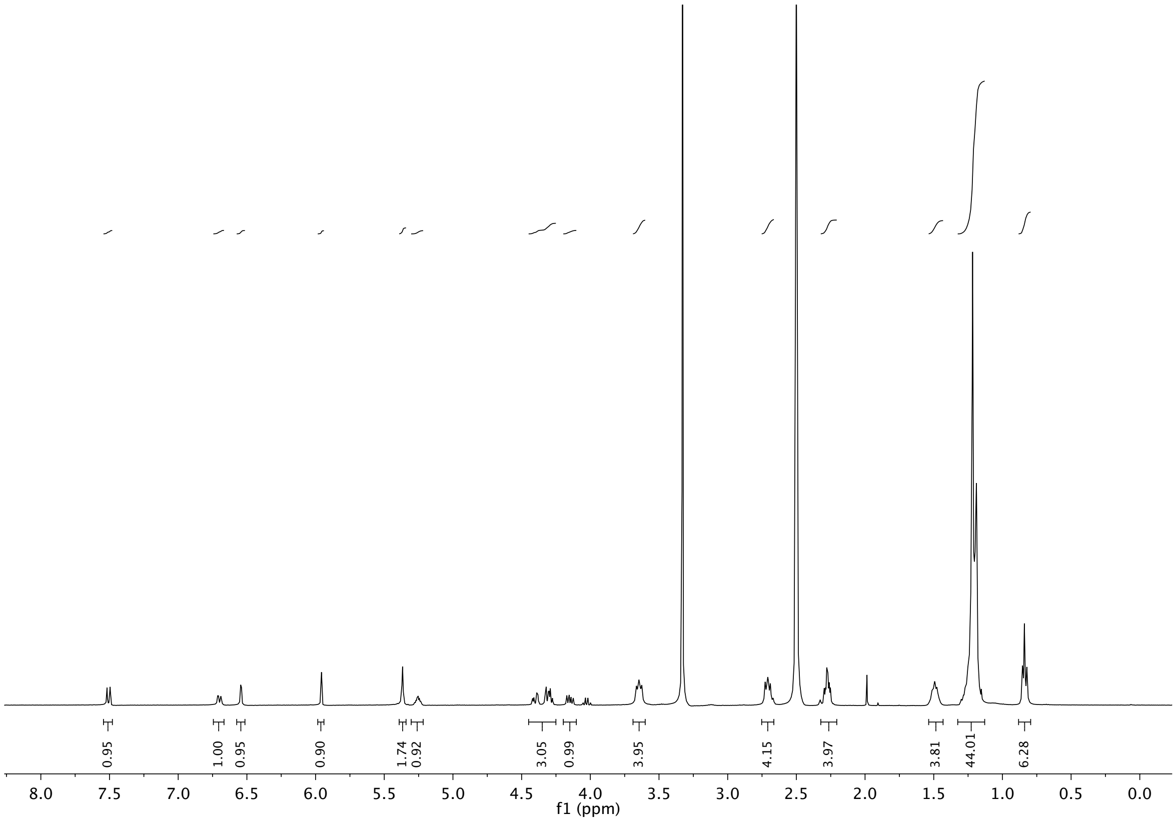

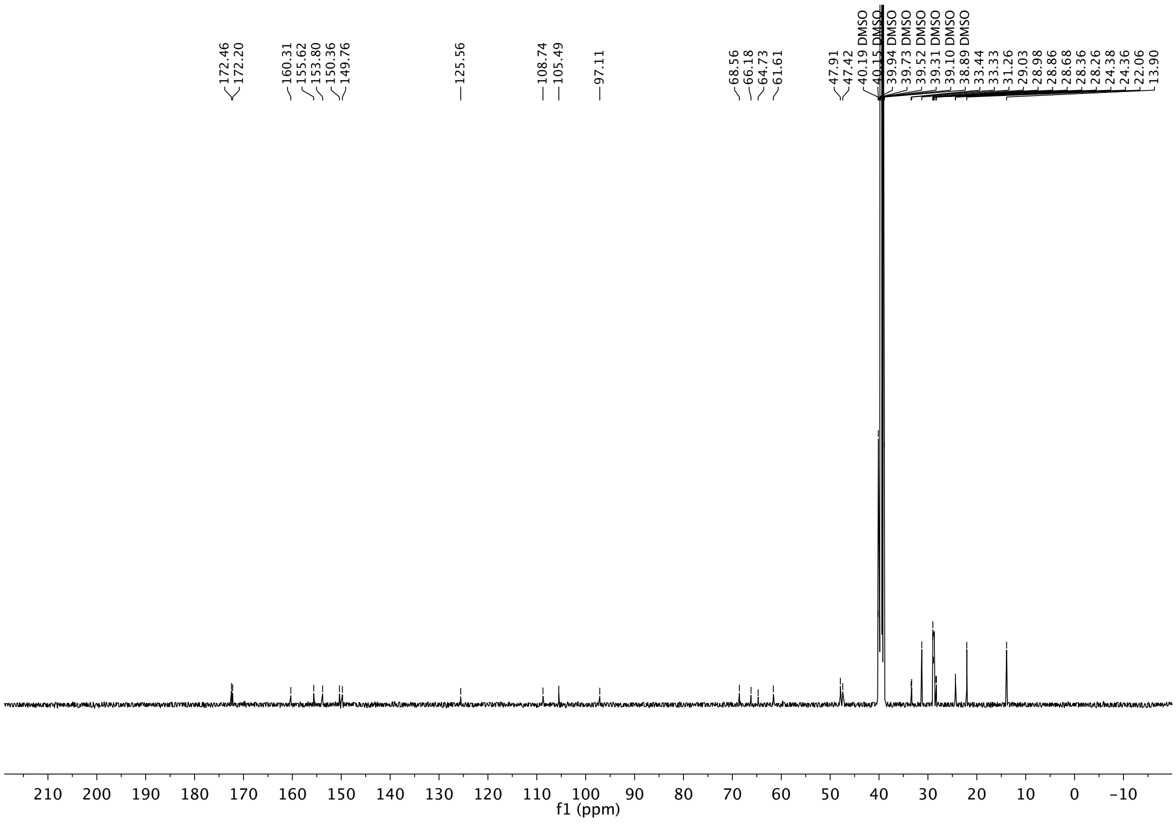

**Figure S28. NMR spectra of (22a)**

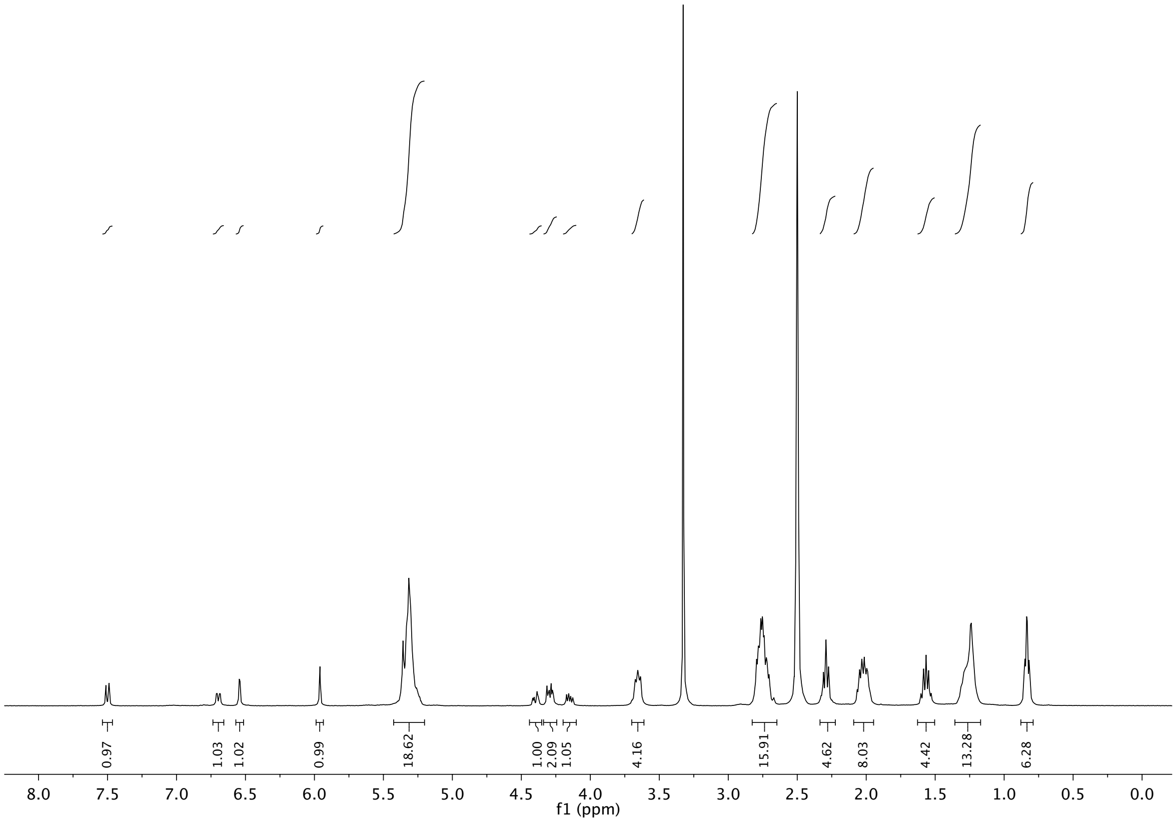

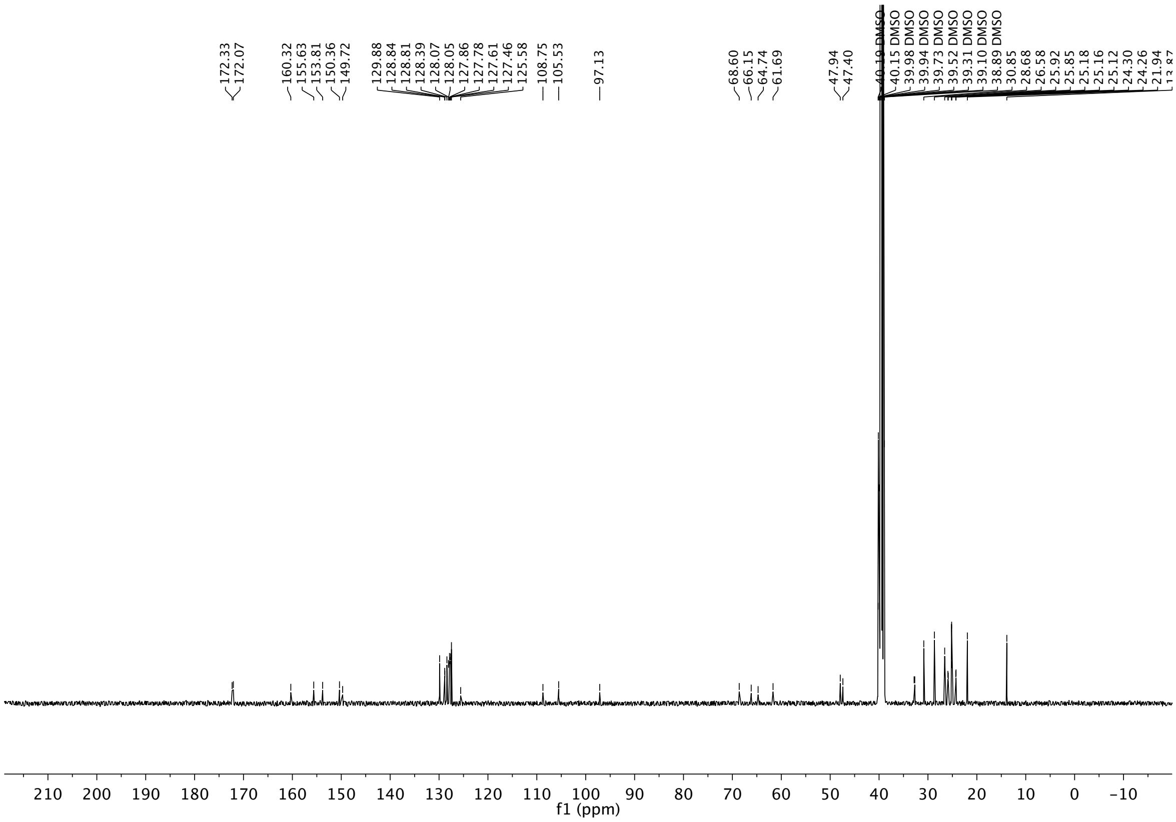

**Figure S29. NMR spectra of (22b)**

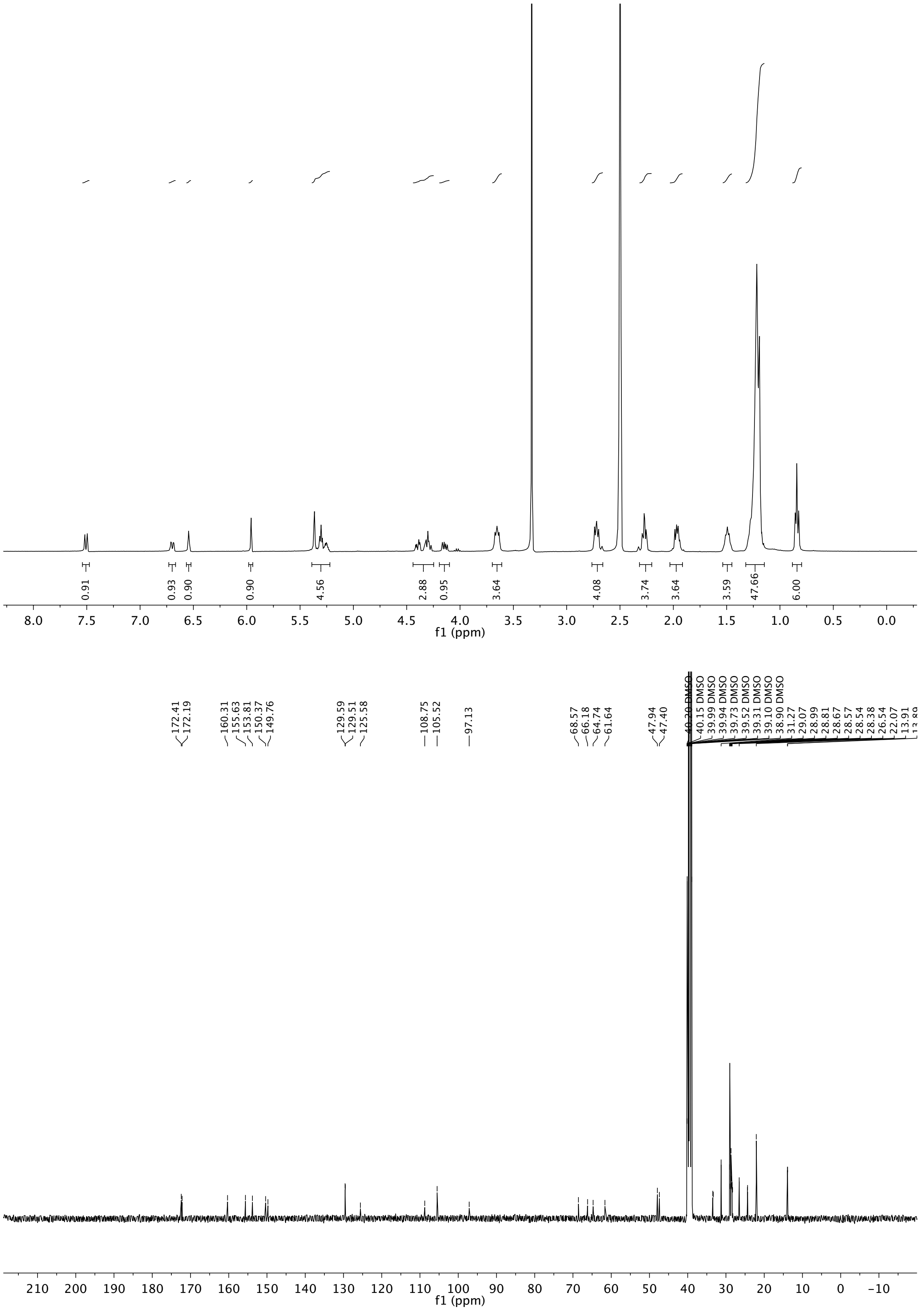

**Figure S30. NMR spectra of (22c)**

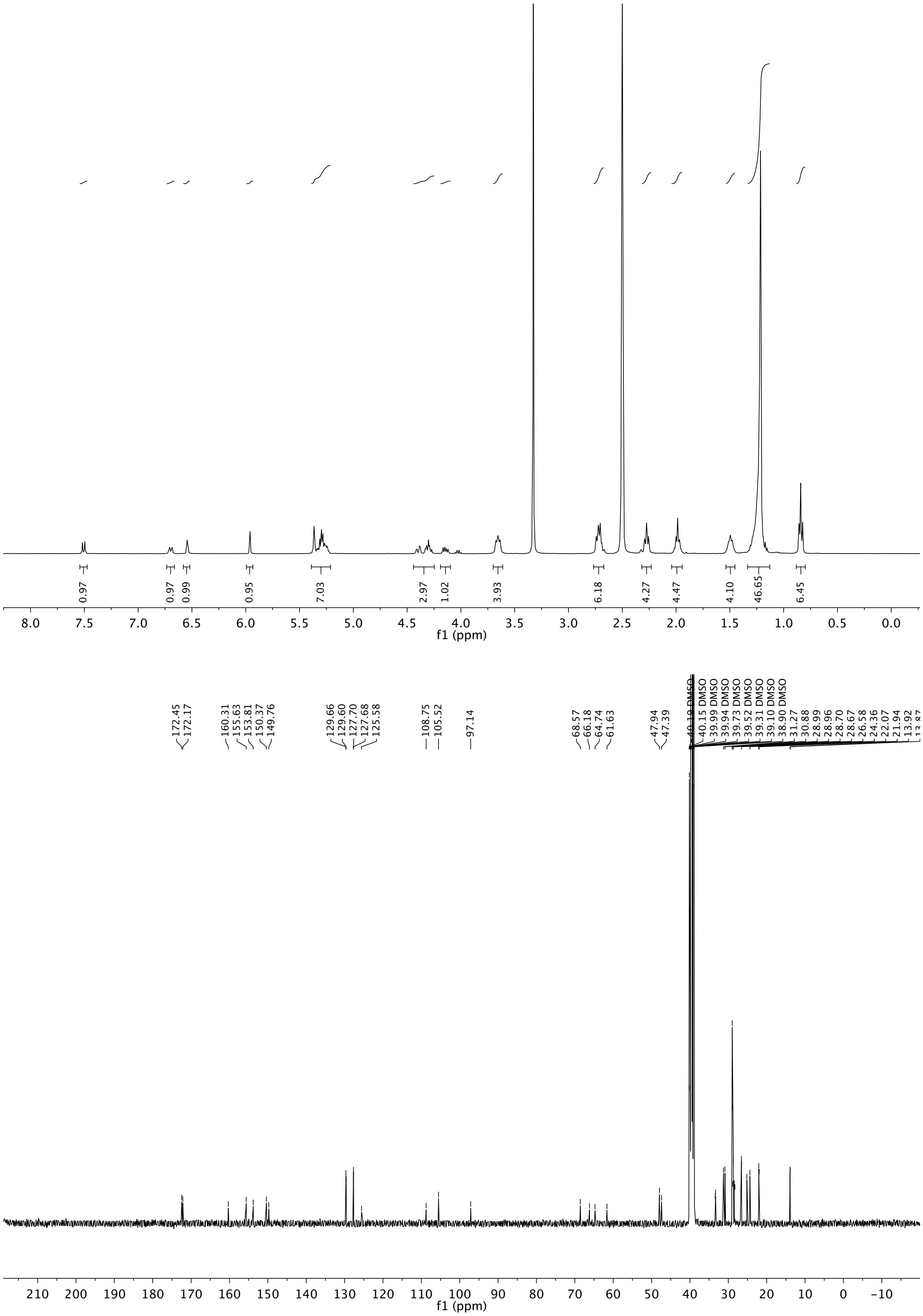

**Figure S31. NMR spectra of (22d)**
